# Distinct genomic architectures but the same gene underlie the convergent evolution of a plant supergene

**DOI:** 10.1101/2025.10.01.679701

**Authors:** Giacomo Potente, Narjes Yousefi, Rimjhim Roy Choudhury, Stefan Grob, Irina A. Gavrilina, Barbara Keller, Emiliano Mora-Carrera, Péter Szövényi, Rebecca L. Stubbs, Hanna Weiss-Schneeweiss, Eva M. Temsch, Gerald M. Schneeweiss, Matthias H. Hoffmann, Giulio Formenti, Ann M. Mc Cartney, Alice Mouton, Henrique G. Leitão, Genevieve Diedericks, Hannes Svardal, Maria Angela Diroma, Chiara Natali, Claudio Ciofi, Étienne Léveillé-Bourret, Elena Conti

## Abstract

Evolution reflects a balance between innovation and constraint, often repurposing existing components in new contexts. Convergent evolution exemplifies this interplay, with similar traits evolving independently in different species, yet the genomic mechanisms enabling such repeatability remain poorly understood. Here, by analyzing ten chromosome-scale genome assemblies, including seven newly generated, we discovered that the *S*-locus supergene (a cluster of tightly linked genes controlling a floral dimorphism called distyly) arose independently multiple times within the primrose family, Darwin’s iconic system for studying distyly. In each case, the same gene was independently duplicated and co-opted, yet the resulting genomic architectures differed, ranging from hemizygous (present on one chromosome copy) to heterozygous (on both copies). These diverse architectures shaped supergene evolution differently, with genetic degeneration occurring only in the heterozygous case. By uncovering multiple mechanisms for supergene origins, our work shows how convergent evolution can produce similar phenotypes by reusing the same genetic building blocks while exploring distinct genomic configurations.

## Introduction

Convergent evolution, defined as the independent acquisition of similar traits in distinct lineages (*1*), is a central topic in evolutionary biology, for it showcases the power of natural selection in driving the repeated emergence of similar adaptive traits under similar selective pressures (*2*). Understanding the genetic basis of convergence is a central question in evolutionary biology, and although recent advances in genomics have made it more accessible, such investigations remain challenging when convergent traits are controlled by multiple, often dispersed, genes across the genome. Conversely, supergenes, i.e. clusters of tightly linked genes controlling a set of co-adapted polymorphic traits (*3*, *4*), offer a two-fold advantage for studying convergent evolution: as simple Mendelian loci, they can be readily associated with phenotypic traits, and enable exploring the role of genomic architecture in adaptation and convergence (*5*). When candidate genes of convergent traits are known, it is possible to discern whether phenotypic convergence arises from mutations in the same genes, mutations in different genes with a shared biochemical pathway, or the recruitment of entirely different genes producing the same phenotypic outcome (*6*). Additionally, while extensive research has focused on identifying genes responsible for phenotypic convergence, the influence of genomic architecture on convergent evolution remains underexplored.

One of the best studied supergenes is the *S*-locus controlling distyly, a floral dimorphism characterized by the coexistence within the same species of two floral morphs, called “pin” and “thrum”, with reciprocally positioned sexual organs promoting cross-pollination (*7–9*) (**Fig. 1b**). Having evolved independently multiple times across angiosperms (*10*), distyly represents a prime example of phenotypic convergence. Recently, *S*-loci were characterized in several species from distantly related angiosperm taxa, revealing diverse genes controlling distyly, but a convergent genomic architecture, with the *S*-locus being hemizygous in thrums (*S*/*s*) and absent in pins (*s*/*s*)(*11–20*) (**Fig. 1c**). Furthermore, in five species that independently evolved distyly, style length dimorphism is controlled by an *S*-locus gene (hereafter *S*-gene) that inactivates brassinosteroids (*16–24*). In Primulaceae, phylogenetic analysis inferred three independent origins of distyly: in the ancestor of *Primula*, in *Hottonia palustris*, and in *Androsace vitaliana* (**Fig. 1a**) (*25*). The *Primula S*-locus contains four core genes (*CYP^T^*, *GLO^T^*, *KFB^T^*, and *PUM^T^*) (*13*, *26*, *27*), two of which have been functionally characterized: in thrums, *CYP^T^* determines short style (*20*) and female incompatibility (*21*) by inactivating brassinosteroids, while *GLO^T^* elevates anthers (*28*).

**Fig. 1:**
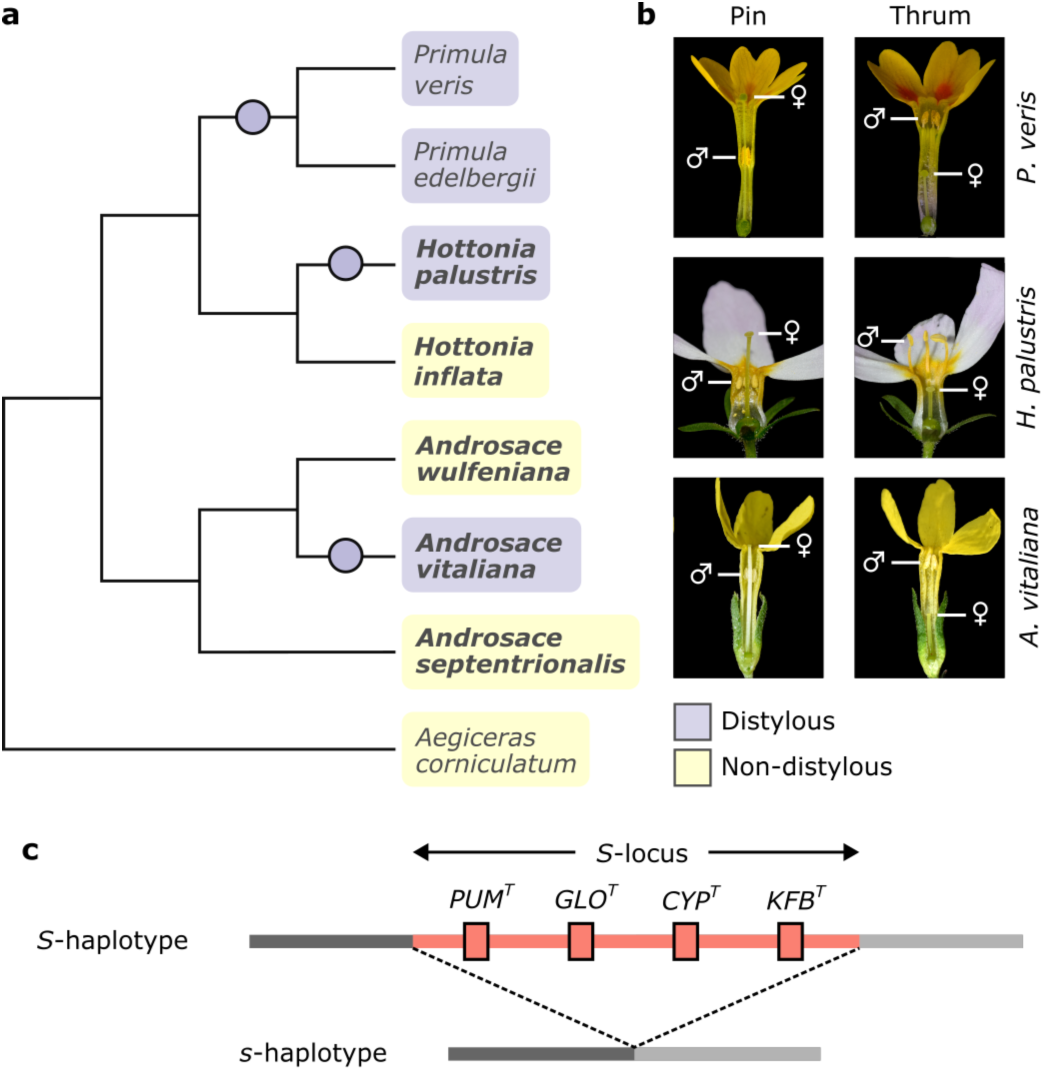
Multiple inferred origins of distyly in Primulaceae. **a.** Cladogram describing relationships among the eight Primulaceae species whose chromosome-scale assemblies were used in this study. Distylous and non-distylous species are highlighted in purple and yellow, respectively; species whose assembly are presented here for the first time are boldfaced. Haplotype-phased assemblies were generated for the distylous *H. palustris* and *A. vitaliana*. Purple circles indicate the three origins of distyly previously inferred in a phylogenetic study of 265 Primulaceae species; *Primula* comprises 526 species, of which 82% distylous; *Hottonia* comprises only two species, of which only *H. palustris* is distylous; *Androsace* comprises 175 species, of which only *A. vitaliana* is distylous (*25*). **b.** Flowers of three distylous species: pins (left) have stigma above anthers, while thrums (right) have anthers above stigma. **c.** Schematic representation of the *S*-locus in *Primula*, in which it is a genomic region containing four core genes present only in the dominant *S*-haplotype and absent from the recessive *s*-haplotype. In all distylous species studied so far, the *S*-locus is hemizygous in thrums (*S*/*s*), and absent in pins (*s*/*s*).

Here, we analyze ten chromosome-scale genome assemblies, seven newly generated, from eight Primulaceae species to ask the following questions: 1) Since distyly is inferred to have evolved multiple times within Primulaceae (*25*), did the *S*-locus also evolve repeatedly, or did it originate once deep in the phylogeny, followed by independent losses or modifications in non-distylous lineages? 2) Do the same genes and genomic architectures underpin distyly in Primulaceae? 3) Do supergenes evolve via colocalization of already functionally interacting genes, or via neofunctionalization of already colocalized genes? 4) Which evolutionary processes drive the expansion of suppressed recombination? 5) Do supergenes with different genomic architectures undergo different evolutionary trajectories? This study significantly advances the understanding of the relationship between phenotypic and genotypic convergence and sheds light on fundamental questions on supergene evolution.

## Results and Discussion

### Comparative genomic analyses of distylous and non-distylous species reveal a lack of inter-specific synteny

To investigate the *S*-locus origins in Primulaceae, we assembled a global data set comprising ten genomes from eight species. Of these, seven chromosome-scale assemblies from five Primulaceae species were newly generated using a combination of short- and long-read sequencing and Hi-C scaffolding. The new assemblies comprised the distylous *Androsace vitaliana* and *Hottonia palustris* (pin and thrum haplotypes were assembled separately for both species), and the closely related, non-distylous *H. inflata* (sister of *H. palustris*), *A. wulfeniana*, and *A. septentrionalis* (**Table 1; Fig. S1-S3; Table S1-S13**). High BUSCO completeness scores were obtained for all assemblies and gene annotations (**Fig. S4**). Comparative genomic analyses revealed an overall lack of whole-genome synteny among genera (**Fig. S5**). Furthermore, we identified a whole-genome duplication (WGD) that likely occurred in the common ancestor of *A. vitaliana* and *A. wulfeniana* but was not shared with *A. septentrionalis* (**Fig. S6**), confirming previous studies (*29*).

**Table 1:**
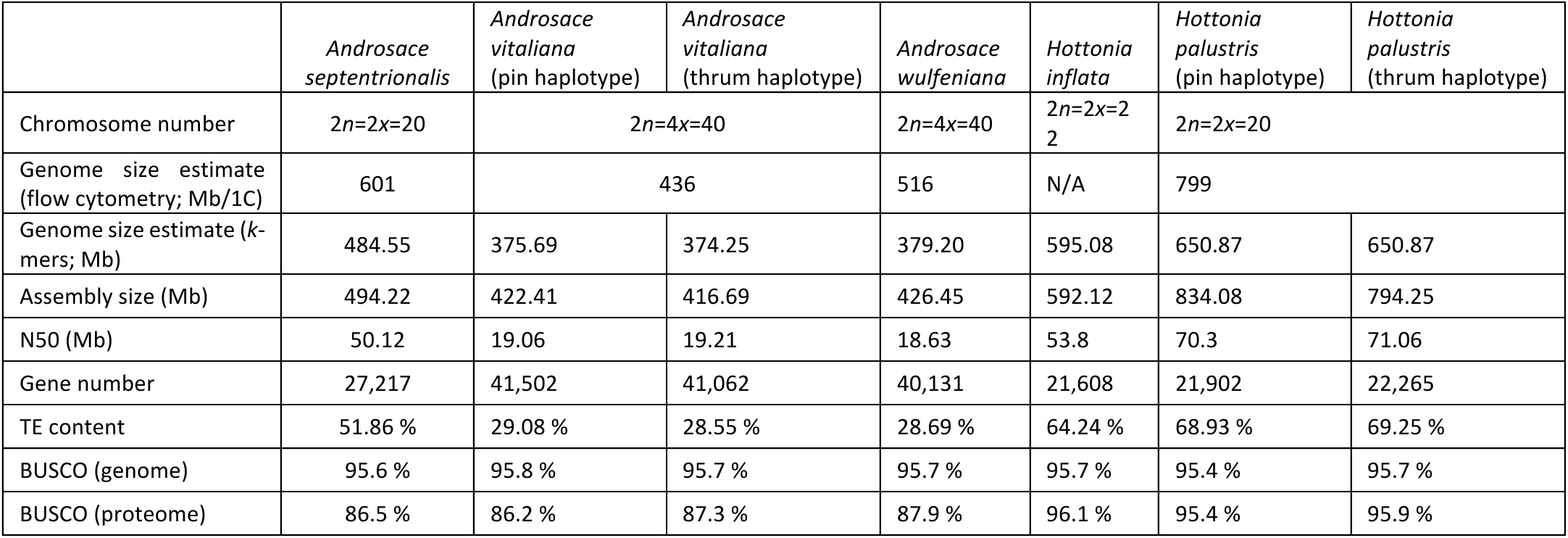
Summary of the genome assemblies generated for this study.

### The *S*-locus of *Hottonia* and *Primula* evolved in their common ancestor

To discover the genetic basis of distyly in *H. palustris*, we first determined which morph contains the *S*-locus by searching for morph-specific *k*-mers, i.e. unique sequences present in one floral morph but absent in the other. We identified 7,023,864 thrum-specific and only 107 pin-specific *k*-mers, suggesting that thrums bear the *S/s* genotype (**Fig. 2a**), as in all distylous species studied to date (*11–13*, *15–17*, *19*). Since the *S*-locus co-segregates with the thrum phenotype, we searched for a region showing high differentiation between morphs. We therefore aligned the thrum-specific *k*-mers onto the genome assembly and discovered that most (99.99%) mapped to a 12.77-Mb region that includes 115 genes and spans the putative centromere of chromosome 9 (36.81-49.56 Mb; **Fig. 2b**). The same genomic region was characterized by elevated differentiation between morphs (expressed as higher *F_ST_* and *D_XY_*), increased heterozygosity in thrums versus pins, and reduced recombination in thrums (**Fig. 2c-f, S7, S8**). The high genetic divergence between floral morphs unambiguously pinpoints this region as the *S*-locus. A synteny analysis between the two haplotypes revealed that the *S*-locus was mostly heterozygous; however, 33 and 25 genes were unique to the *s*- and *S*-haplotype, respectively (**Fig. 2g, S9, S10**). Notably, five of the 25 *S*-haplotype-specific hemizygous genes were orthologous to the *Primula S*-genes: *HpGLO^T1^* and *HpGLO^T2^* (tandemly duplicated copies of *Primula GLO^T^*), *HpPUM^T^*, *HpCYP^T^*, and *HpKFB^T^* (**Table S14**). These five hemizygous genes spanned 5.14 Mb and were intermingled with heterozygous genes.

**Fig. 2:**
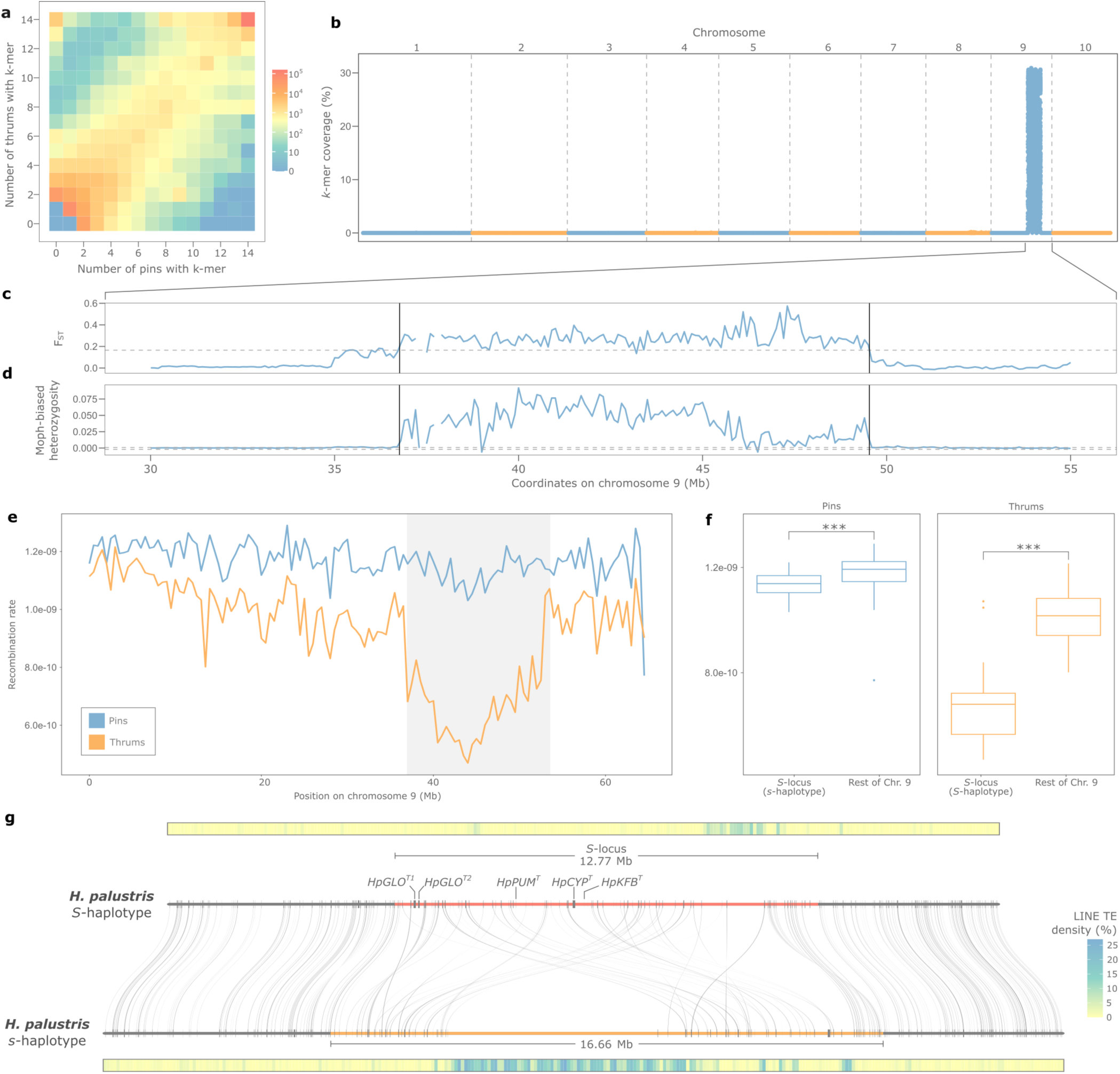
The *Hottonia palustris S*-locus is heterozygous but contains hemizygous genes, including the orthologs of *Primula S*-genes. **a.** Heatmap illustrating *k*-mer distribution across combinations of pin (x-axis) and thrum (y-axis) individuals. Each cell’s color represents the *k*-mer count for that specific combination of thrums and pins. The increased *k*-mer count in the top-left corner compared to the bottom-right corner indicates an abundance of thrum-specific over pin-specific *k*-mers. **b.** Percentage of sequence covered by thrum-specific *k*-mers (calculated in 5-kb windows) across the *H. palustris* assembly. **c-d.** Distribution of *F_ST_* (**c**) and morph-biased heterozygosity (**d**), in 100-kb windows across chromosome 9 (30-55 Mb). The *S*-locus borders are marked by black vertical lines. Dotted horizontal lines represent the 95th percentile of the two distributions. **e.** Recombination rate in 14 pin (blue) and 14 thrum (orange) individuals, in 50-kb windows across chromosome 9. The *S*-locus is highlighted in grey. **f.** Box plots showing that recombination rate is significantly lower in the *S*-locus than in the rest of chromosome 9 for both pins (left) and thrums (right) due to its pericentromeric location, but more pronounced in thrums, indicating that the *S*-haplotype is not recombining (***, p-value < 0.001; Wilcoxon rank-sum test. **g.** Microsynteny between the *S*- and *s*-haplotypes. The orthologs to *Primula S*-genes are indicated in the figure. On top and bottom of the two haplotypes are density plots representing the proportion of sequence covered by LINE TEs (calculated in 100-kb windows), which are enriched in centromeric regions (see also Fig. S3).

The occurrence of the same *S*-genes in the *S*-haplotypes of *Primula* and *H. palustris* suggests that the *S*-locus originated before the divergence between *Primula* and *Hottonia*. If so, the absence of distyly in *H. inflata* would reflect a secondary loss caused by loss-of-function mutations in its *S*-locus. Indeed, we found that the *S*-locus was present but degenerated in *H. inflata*, as it lacked both *CYP^T^* and *GLO^T^*, while containing *PUM^T^* and *KFB^T^* (**Fig. S11**). Given *CYP^T^* and *GLO^T^* functions (*20*, *21*, *28*), individuals with an *S*-locus lacking these two genes are expected to have the stigma above the anthers and be self-compatible, consistent with the floral morphology of *H. inflata* and its tendency to self-pollinate (*30*). *Hottonia inflata* therefore showcases a novel way of losing distyly, since the loss of distyly previously described both in populations of *Primula vulgaris* (*31*, *32*) and in the non-distylous *Primula grandis* (*33*) involved loss-of-function mutations affecting only *CYP^T^*, but not *GLO^T^*.

Our results show that the *S*-locus of *Primula* and *Hottonia* originated in their common ancestor and was independently lost in non-distylous *Primula* species and *H. inflata*, contrary to the phylogenetically inferred hypothesis that distyly evolved independently in *Primula* and *H. palustris* (*25*). Unlike the fully hemizygous *S*-loci characterized so far (*11–13*, *15*, *17*, *19*), the *H. palustris S*-locus contains both heterozygous and hemizygous genes, like the distantly-related *Turnera*, whose *S*-locus consists of three hemizygous genes flanked by 17 heterozygous genes contained in two inversions (*16*). Furthermore, the genomic region containing the *S*-locus in *H. palustris* was not syntenic to the one containing the *S*-locus in either *P. veris* or *P. edelbergii*, suggesting that the *S*-locus translocated between chromosomes at least twice since its origin in the common ancestor of the two genera (**Fig. S12**). However, despite multiple translocations, the *S*-locus retained a pericentromeric location in all *Primula* and *Hottonia* species studied so far (*26*, *27*), suggesting strong selective pressure to maintain it in a region characterized by reduced recombination.

### The *Androsace vitaliana S*-locus originated independently from the *Primula*-*Hottonia S*-locus

To search for the *S*-locus of *A. vitaliana*, we adopted the same approach as for *H. palustris* (see above). We observed more thrum-specific than pin-specific *k*-mers (18,990 vs 2,896), indicating that thrums bear the *S/s* genotype also in this species (**Fig. 3a**). The majority of thrum-specific *k*-mers, as well as elevated *F_ST_*, were observed in a ca. 70-kb region at one end of chromosome 5 (975-1,045 kb; **Fig. 3b-c, S13-S16**). The same region was also characterized by increased heterozygosity in thrums compared to pins, hence representing the *S*-locus (**Fig. 3d**). Synteny analysis showed that three genes are contained in the *S*-locus and are present in both haplotypes in the same order and orientation (**Fig. 3e**). Following the same naming convention used in *Primula* (*13*), we name these genes *AvCSE^T^*, *AvEH^T^*, and *AvCYP^T^*for *S*-alleles, where “*T*” refers to thrums, and *AvCSE^P^*, *AvEH^P^*, and *AvCYP^P^* for *s*-alleles, where “*P*” refers to pins. Caffeoyl shikimate esterases (*CSE*) participate in lignin biosynthesis, while epoxide hydrolases (*EH*) are involved in lipid biosynthesis, and both are presumed or confirmed to be involved in defense against pathogens. (*34*, *35*) This could argue for a role in self-incompatibility, as defense, lipid and lignin biosynthesis genes are up- or down-regulated in response to incompatible pollination in maize (*36*).

**Fig. 3:**
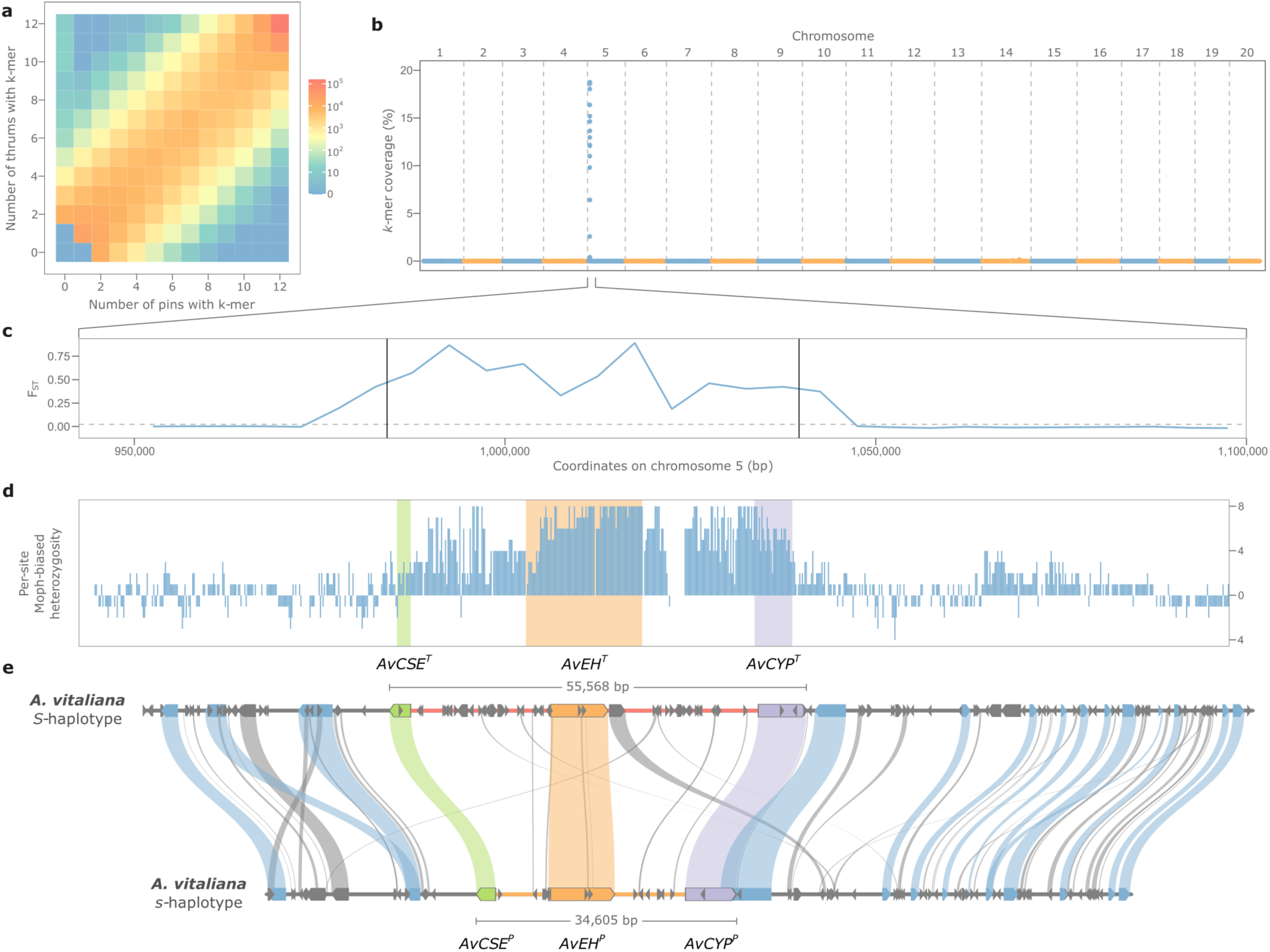
The *Androsace vitaliana S*-locus is heterozygous and consists of three genes, including one orthologous to a *Primula S*-gene (*CYP^T^*). **a.** Heatmap illustrating *k*-mer distribution across combinations of pin (x-axis) and thrum (y-axis) individuals. Each cell’s color represents the *k*-mer count for that specific combination of thrums and pins. The increased *k*-mer count in the top-left corner compared to the bottom-right corner indicates an abundance of thrum-specific over pin-specific *k*-mers. **b.** Percentage of sequence covered by thrum-specific *k*-mers (calculated in 5-kb windows) in the *A. vitaliana* assembly. **c.** *F_ST_* distribution, calculated in 5-kb windows, across chromosome 5. The increase in *F_ST_* overlaps with the region enriched for thrum-specific *k*-mers and represents the *S*-locus, whose borders are marked by black vertical lines. **d.** Bar plot representing morph-biased heterozygosity, calculated as the number of thrums carrying the heterozygous genotype minus the number of pins carrying the heterozygous genotype for 882 SNPs (chromosome 5: 950-1,100 kb) identified in 16 individuals (8 thrums and 8 pins). **e.** Microsynteny between the *S*- and *s*-haplotype. Genes and TEs are represented as blue and gray boxes, respectively; *S*-genes are colored in green (*AvCSE*), orange (*AvEH*), and purple (*AvCYP*); the *S*-haplotype is marked as a red line, while the *s*-haplotype as an orange line.

If the *S*-alleles played a role in controlling distyly, we would expect them to be upregulated compared to *s*-alleles in floral tissues. Indeed, we observed that, in thrum flowers, *AvEH^T^*and *AvCYP^T^* were upregulated compared to *AvEH^P^*and *AvCYP^P^* (**Fig. S17**), while *AvCSE* alleles were expressed at the same level. Our results therefore align with the hypothesis that the *S*-alleles *AvEH^T^* and *AvCYP^T^* have a function in controlling distyly by showing evidence of upregulation compared to their respective *s*-alleles in floral tissues. However, functional studies will be necessary to determine *AvCSE^T^*function and whether it plays a role in distyly.

The *A. vitaliana S*-locus represents the first described distyly *S*-locus that is entirely heterozygous, for it consists of three genes present in both haplotypes. The only detected difference between haplotypes is the accumulation of transposable elements (TEs) in the *S*-haplotype compared to the *s*-haplotype (**Fig. 3e**), but whether this is a cause or consequence of suppressed recombination remains to be investigated. Another distinctive feature of the *A. vitaliana S*-locus is its non-pericentromeric location, unlike the pericentromeric *S*-loci of *Primula* and *Hottonia*. It has been proposed that supergenes are more likely to emerge in regions already characterized by reduced recombination, such as pericentromeric regions (*3*, *4*, *37*). Indeed, several examples support this hypothesis, such as the sex-determining regions of kiwifruit (*38*), *Nepenthes* (*39*), and *Rumex* (*40*), and the self-incompatibility locus in *Petunia* (*41*). However, the *A. vitaliana S*-locus, along with other examples of supergenes located in actively recombining regions, e.g. the distyly *S*-locus of *Linum tenue* (*12*) and the sex-determining region of *Silene latifolia* (*42*), challenges this hypothesis, suggesting that supergene emergence is not restricted to regions of reduced recombination. Finally, we note that *A. vitaliana* represents another independently-evolved *S*-locus containing a gene involved in brassinosteroid inactivation (*AvCYP^T^*), as described in several other distylous species (*15–24*).

### The same gene (*CYP^T^*) was independently recruited in the *S*-loci of *Androsace* and *Primula-Hottonia*

Previous phylogenetic analyses on the *S*-locus in *Primula* estimated the duplication ages of *CYP^T^* and *GLO^T^* around 43 and 37 Mya, respectively (*13*, *26*, *28*). Such dates precede *Primula* divergence from *Hottonia* by 12-18 My, foreshadowing our current demonstration of the shared origin of distyly in the two genera. However, the sparse sampling of *Primula*’s closest relatives left much uncertainty regarding the evolution of neofunctionalization in these key *S*-genes.

The genomic data presented here allowed us to improve estimates of *S*-gene phylogenetic histories (**Fig. S18-S24**). We inferred node-dated phylogenies for all gene families comprising the *S*-genes identified in Primulaceae. Five gene phylogenies supported the previously identified Primulaceae WGD event (*Pv-α*) (*26*), dating it at 53 (35–81) Mya. Notably, *KFB^T^* and *PUM^T^* of *Primula* and *Hottonia* seemingly originated through this WGD event. The other *S*-genes in these two genera originated through more recent duplication events (*CCM^T^*: 7.7 Mya; *GLO^T^*: 36 Mya; *CYP^T^*: 45 Mya). Furthermore, our analyses revealed that the closest paralog of *CYP^T^*is not *CYP734A51*, as previously suggested (*20*), but a newly-discovered paralog present in *Hottonia* genomes but absent from all *Primula* genomes sequenced to date (**Fig. 4**). The gene phylogenies also strongly support a WGD event shared between *A. vitaliana* and *A. wulfeniana* around 8.0 (3.7-24.0) Mya, likely before the divergence between the *S-* and *s*-alleles of *A. vitaliana* dated at 7.3 (2.7-13.0) Mya for *EH*, 1.7 (0.6-2.9) Mya for *CYP*, and 1.4 (0.3-3.0) Mya for *CSE*.

**Fig. 4:**
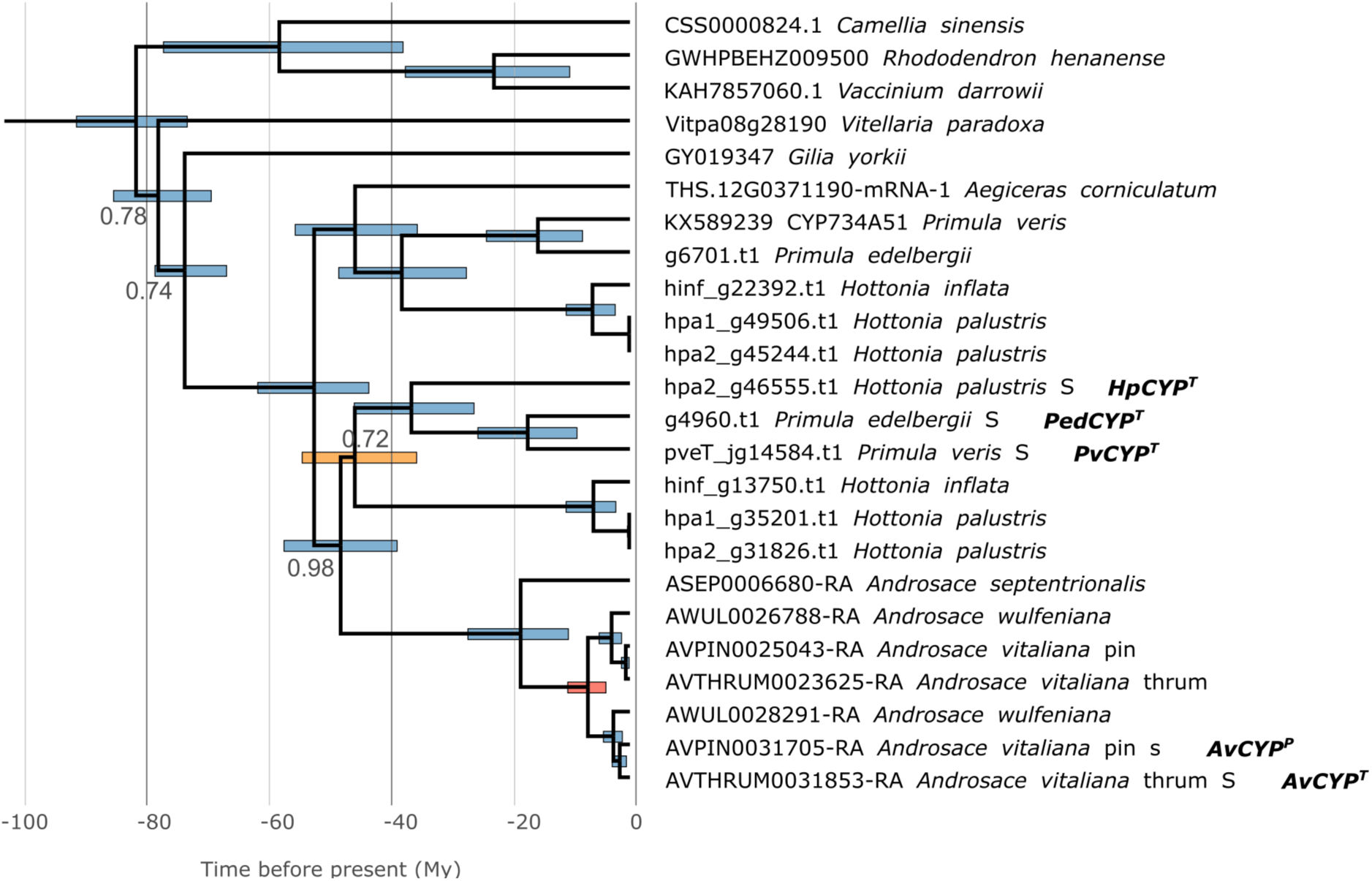
Phylogeny of the *S*-gene *CYP_T_* and close homologs. Bayesian chronogram of *CYP734* genes in selected genomes. Bottom scale bar indicates time before present in million years (My). Blue bars at nodes represent 95% Bayesian credibility intervals around age estimates, the red bar represents the same for the *Androsace* WGD that gave rise to *AvCYP* in *Androsace*, while the orange bar represents the same for the duplication event that gave rise to *CYP_T_* in *Primula* and *Hottonia* and its closest paralog. Branch labels represent posterior probabilities <1, while those without a numeric label have posterior probabilities of 1.

Interestingly, *AvCYP^T^* originated via duplication of the same gene whose duplication gave origin to the *Primula*-*Hottonia CYP^T^*. This means that duplicates of the same gene were independently and asynchronously recruited to create the distylous phenotype, first in the most recent common ancestor (MRCA) of *Primula* and *Hottonia*, and later in *A. vitaliana*. In both cases, neofunctionalization of *CYP^T^* was preceded by a duplication event: in the MRCA of *Primula* and *Hottonia* this was an isolated duplication followed by translocation to the *S*-locus, while in *A. vitaliana* this occurred in the *Androsace*-specific WGD. Yet, all these gene copies coalesce in the MRCA of Primulaceae, making them orthologs likely with the same or very similar functions.

In distantly related taxa, the convergent evolution of distyly often involves different genes acting within the brassinosteroid pathway (*15–24*). Here we showed that within Primulaceae, the convergent evolution of distyly occurred via repeated recruitment of the same brassinosteroid-inactivating gene, *CYP^T^*. These results align with the notion that gene reuse in convergence is more likely among closely related lineages (*43*, *44*).

### The *S*-locus of *Androsace vitaliana* originated via a selective sieve

Two main models have been proposed for the origin of supergenes. According to the “Turner’s sieve” model, physically linked genes eventually acquire mutations that make them functionally interact with each other (*45*). Conversely, the “translocation” model posits that functionally interacting but genomically dispersed genes are later brought into proximity via translocation (*46*).

To test whether the physical linkage among *AvCSE*, *AvEH*, and *AvCYP* precedes the emergence of distyly, we performed synteny analyses among nine Primulaceae genomes of both distylous and non-distylous species and observed that these three genes are collinear across all species (**Fig. S25**), suggesting that they already colocalized prior to the evolution of functional interaction among them. Thus, the *S*-locus of *A. vitaliana* evolved as described by “Turner’s sieve”. Notably, this occurred following a WGD (**Fig. S6**), whereby the *AvCSE*, *AvEH*, and *AvCYP* paralogs, likely owing to relaxed selection on gene duplicates (*47*), could accumulate mutations that ultimately led to their neofunctionalization and recruitment in the control of distyly.

The *S*-loci of most distylous species studied to date originated via stepwise gene duplications and translocations (*24*). Instead, *A. vitaliana* fits the so-called “segmental duplication” model (*48*), as its *S*-genes duplicated simultaneously through a WGD. This scenario parallels recent findings in Oleaceae, where the *S*-locus controlling self-incompatibility and distyly also arose via neofunctionalization of colocalizing genes following WGD, albeit involving a different gene set (*15*).

### Distinct genomic architectures affect selection on *S*-loci differentially

As non-recombining regions, supergenes are subject to reduced efficacy of selection and expected to undergo genetic degeneration, a phenomenon characterized by the accumulation of deleterious mutations and TEs (*49*). Degeneration is predicted to be more severe in heterozygous supergenes, in which recessive deleterious alleles can be masked by dominance, than hemizygous supergenes, where recessive deleterious alleles are always exposed (*3*). These theoretical predictions received support from forward simulations (*3*), but empirical validation in natural systems remains scarce. Primulaceae *S*-loci are an ideal system to test the above predictions by comparing selection on supergenes that are fully hemizygous (*P. veris*), fully heterozygous (*A. vitaliana*), or comprise both heterozygous and hemizygous genes (*H. palustris*).

We identified a clear distinction between selective forces acting on hemizygous versus heterozygous *S*-genes. In *H. palustris*, *S*-alleles accumulated more slightly deleterious mutations than the genomic background, i.e. they were enriched in genes with significant p-value for the McDonald-Kreitman (MK) test and negative direction of selection (DoS) (Fisher’s exact test; *P* < 0.001); conversely, all *s*-alleles evolved under neutrality, i.e. no *s*-allele had a significant p-value for the MK test (**Fig. 5b; Table S15**). DoS could not be computed for 19/20 hemizygous *S*-genes due to paucity of polymorphic and/or divergent sites; among these, *HpGLO^T1^*, *HpGLO^T2^*, *HpPUM^T^*, *HpCYP^T^*, and *HpKFB^T^* contained no polymorphic sites at all. Similarly, in *A. vitaliana*, *s*-alleles (*AvCSE^P^*, *AvEH^P^*, and *AvCYP^P^*) evolved under neutrality, while *S*-alleles (*AvCSE^T^*, *AvEH^T^*, and *AvCYP^T^*) accumulated slightly deleterious mutations (**Fig. 5c-e; Table S16-S17**). In *P. veris*, DoS could not be computed for any *S*-gene due to absence of polymorphic and/or divergent sites. Overall, our analyses revealed that only heterozygous *S*-genes accumulated slightly deleterious mutations, and only on thrum-specific *S*-alleles, which never recombine. Conversely, hemizygous *S*-genes were characterized by extremely reduced nucleotide diversity, indicating strong purifying selection. Such pattern was observed for hemizygous *S*-genes regardless of whether they were contained in a fully hemizygous supergene (*P. veris*) or in a partially heterozygous supergene (*H. palustris*). Our empirical results thus corroborate theoretical predictions and results of forward simulations (*3*).

**Fig. 5:**
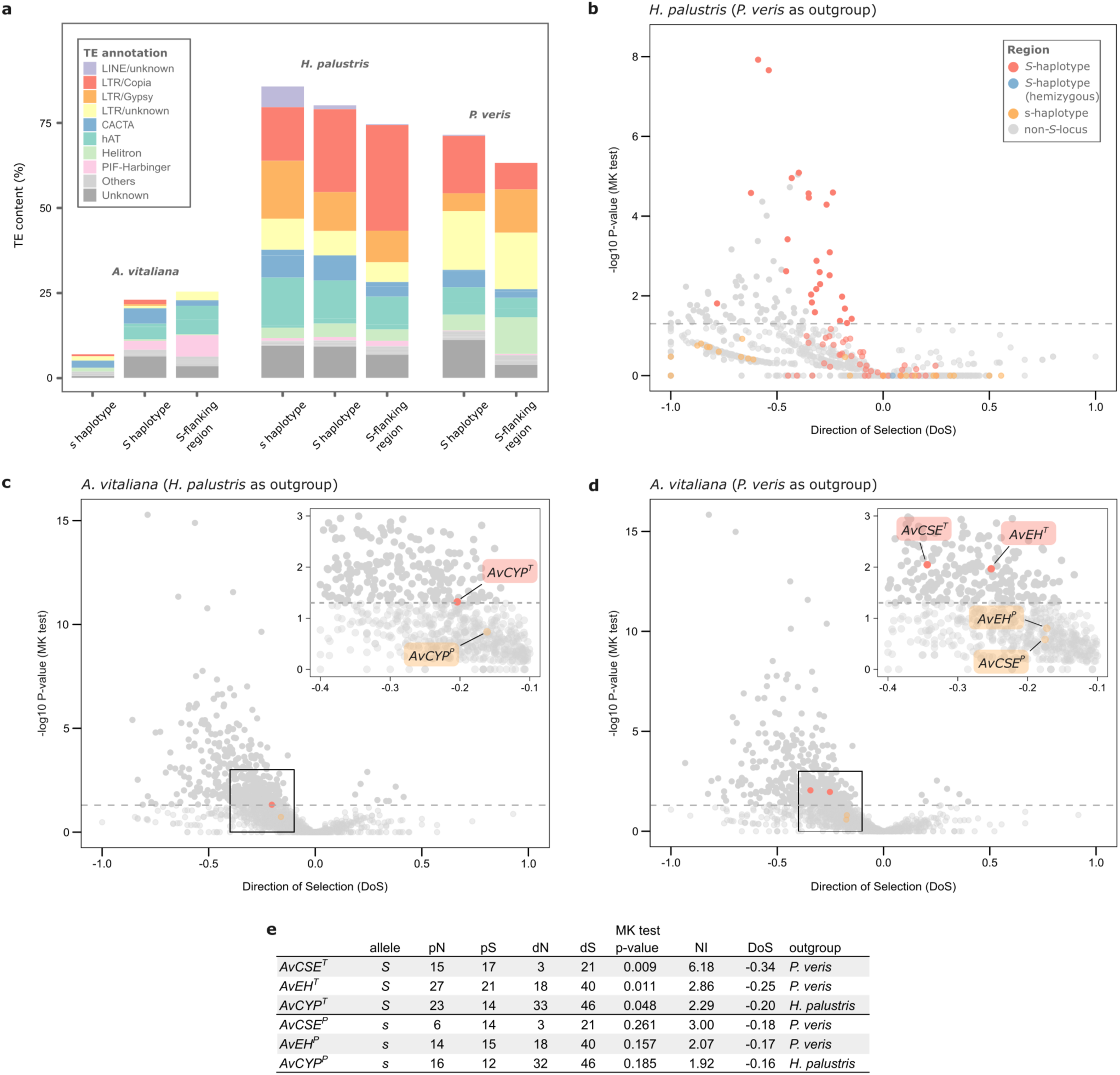
Contrasting patterns of degeneration in heterozygous and hemizygous *S*-loci. **a.** Transposable element (TE) abundance, expressed as percentage of sequence covered by each TE superfamily as in legend, for three distylous species, comparing the *S*-locus (*S*- and *s*-haplotype, when both available) with their flanking regions. **b-d.** Volcano plots representing the direction of selection (DoS, x-axis) and the −log_10_ of the p-value of a McDonald-Kreitman test (y-axis) calculated for all genes in the chromosome carrying the *S*-locus of *H. palustris* (**b**, *P. veris* used as outgroup; n=957), *A. vitaliana* (**c**, *P. veris* used as outgroup; n=1,302), and *A. vitaliana* (**d**, *H. palustris* used as outgroup; n=1,322). Each dot in the volcano plot represents a gene: *S*-genes are red (*S*-alleles), orange (*s*-alleles), or blue (hemizygous *S*-genes), while genes in the rest of the genome are gray. A dashed horizontal line marks the significance threshold of the MK test at p-value = 0.05. The inset plots in (**c**) and (**d**) highlight *S*-genes. **e.** Table summarizing the MK and DoS results for the three *A. vitaliana S*-genes in both haplotypes. pN, polymorphic non-synonymous sites; pS, polymorphic synonymous sites; dN, non-synonymous substitutions; dS, synonymous substitutions; NI, neutrality index. The volcano plots show that the heterozygous *S*-alleles tend to accumulate slightly deleterious mutations, unlike the *s*-alleles and the hemizygous *S*-genes.

Another expected feature of supergenes is an enrichment of TEs (*3*), a pattern observed in most *S*-loci (*12*, *14–18*). Our analyses showed that the Primulaceae *S*-loci were not enriched in TEs compared to the genomic background, and only in *H. palustris* the *S*-locus contained more TEs than its flanking regions (**Fig. 5a; Table S18; Fig. S26-S27**). When testing whether TE abundance was higher in the non-recombining *S*-haplotypes than in the freely recombining *s*-haplotype of heterozygous *S*-loci, differences were detected among species. In *A. vitaliana*, the *S*-haplotype contained 3.7 times more TEs than the *s*-haplotype (25.40% vs 6.94%), consistent with the expectation of non-recombining regions accumulating repetitive elements, while in *H. palustris* the *s*-haplotype contained more TEs than the *S*-haplotype (85.86% and 80.31%, respectively; **Fig. 27**). These findings highlight that the prediction of TE accumulation in supergenes should be approached with caution, as TE accumulation is influenced by multiple factors, including the species’ evolutionary history and the age, architecture, and genomic location of the supergene (*50–52*).

### The *S*-locus expansion in *Hottonia palustris* was driven by non-selective processes

The *S*-locus is much larger in *H. palustris* (12.75 Mb) than in *Primula* (ca. 260 kb), despite sharing a common origin. Two scenarios could explain this: either suppressed recombination expanded beyond the original *S*-locus in *H. palustris*, *de facto* increasing *S*-locus size, or the *S*-locus was originally larger and later contracted in *Primula*. To disentangle between these two possibilities, we calculated synonymous divergence (d_S_) between the two *S*-locus haplotypes of *H. palustris* and between *H. palustris* and *P. veris*. The results showed that recombination suppression between *S*-locus haplotypes in *H. palustris* occurred after the split from *Primula*, thus the *S*-locus expanded in *H. palustris* (**Fig. 6a**). This represents the first description of suppressed-recombination expansion beyond a hemizygous supergene.

**Fig. 6:**
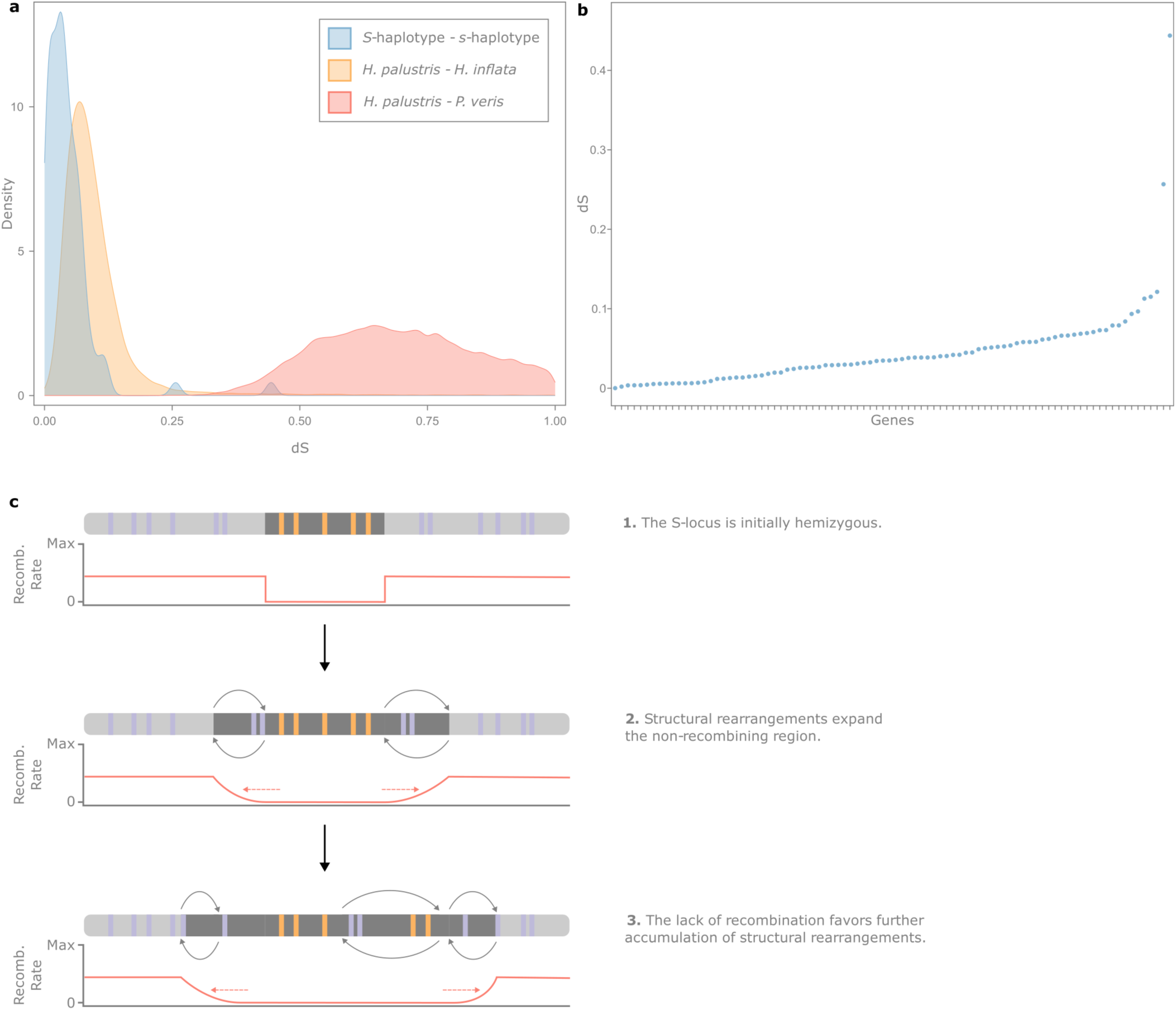
Expansion of suppressed recombination in the *Hottonia palustris S*-locus. **a.** Distributions of d_S_ calculated between syntenic orthologs of *H. palustris - P. veris* (n=15,964; red curve), *H. palustris - H. inflata* (n=18,830; orange curve), and between the *S*- and *s*-haplotype of *H. palustris* (n=88; blue curve). The suppression of recombination in the *H. palustris S*-locus (blue curve) is more recent than the divergence between *H. palustris* and *P. veris* (red curve) and overlapping with the divergence between *H. palustris* and *H. inflata* (orange curve). **b.** d_S_ values obtained between the *S*- and *s*-alleles for the 88 *S*-genes present in both haplotypes. Genes were ordered on the x-axis by increasing d_S_ value. The gradual, rather than stepwise, increase in d_S_ within the *S*-locus indicates the lack of evolutionary strata. **c.** Possible model for the expansion of suppressed recombination in the *H. palustris S*-locus. Regions of suppressed recombination are indicated by dark grey boxes; *S*-genes are colored in orange; other genes are colored in purple. The *S*-locus is initially hemizygous (1) and structural rearrangements (e.g. inversions) occur in its flanking regions, expanding the non-recombining region (2). This process may proceed iteratively, progressively expanding the region of suppressed recombination (3).

This finding raised the question of what drove *S*-locus expansion. A long-standing explanation for the progressive loss of recombination in supergenes and sex chromosomes is antagonistic selection, whereby suppressed recombination locks together beneficial combinations of alleles with antagonistic effects (*53*, *54*). However, empirical support for this model remains limited and alternative, non-selective models have recently been proposed (*55*). Among them, a neutral model (*56*) suggests that mutations can accumulate at the supergene borders and get linked to it by drift, increasing haplotype divergence and further reducing recombination. Importantly, this process is more likely if the supergene lies in a low-recombining region, such as the pericentromeric region containing the *S*-locus in *H. palustris*.

If antagonistic selection drove *S*-locus expansion, we would expect this region to be enriched in genes involved in distyly and show signs of stepwise expansion, such as evolutionary strata (*54*, *57*). Conversely, we wouldn’t expect such signatures if the *S*-locus expanded via non-selective processes (*55*). We found no enrichment in genes associated with distyly in the *H. palustris S*-locus (i.e. it did not contain orthologs of genes that were differentially expressed between pins and thrums in *P. veris* (*58*)), suggesting that the expanded *S*-locus of *H. palustris* does not contain genes with antagonistic effects (**Table S19**). Furthermore, no evolutionary strata were detected (i.e. the distribution of d_S_ calculated between *S*- and *s*-alleles showed a continuous cline rather than a stair-like pattern), pointing to a gradual expansion of suppressed recombination, likely caused by multiple, small structural rearrangements (**Fig. 6b-c**). Therefore, the *S*-locus expansion in *H. palustris* was likely not driven by antagonistic selection, but rather by non-selective processes.

## Conclusions

By investigating the convergent evolution of the *S*-locus controlling distyly in Primulaceae, our study illustrates how evolution operates as an exploration of genomic possibilities, evidenced by the diverse architectures and genes underpinning distyly across the examined species (**Fig. 7**). Yet, this flexibility is tempered by molecular and evolutionary constraints. The results presented here align with François Jacob’s concept of evolutionary tinkering (*59*), which suggests that evolution operates by repurposing existing components, and support Susumu Ohno’s hypothesis highlighting the importance of gene duplications in the evolution of novel phenotypes (*60*). Within this theoretical framework, the repeated co-option of *CYP^T^* in controlling distyly underscores how evolutionary innovation is often context-dependent and constrained by the available genetic toolkit, leading to convergent outcomes through the modification of pre-existing elements. Together, our study not only illuminates the mechanisms underlying supergene evolution but also contributes to a broader understanding of how genomic innovation and constraint interact in convergent evolution.

**Fig. 7:**
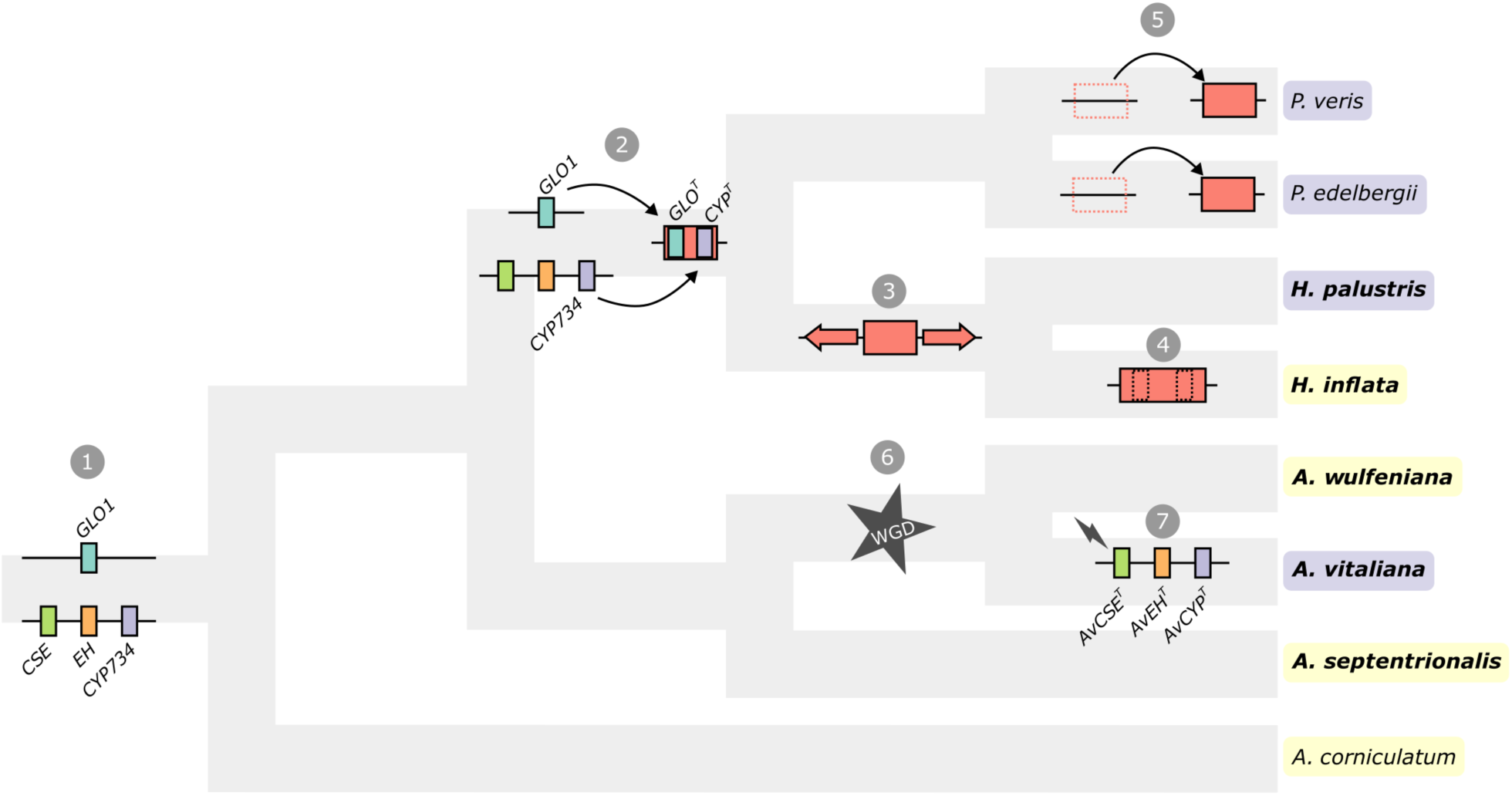
Model of *S*-locus evolution in Primulaceae. The ancestral genomic configuration (1) involved three colocalized (*CSE*, *EH*, *CYP734*) and one non-colocalized gene (*GLO1*). Subsequently, duplications of *GLO1* and *CYP734* in the common ancestor of *Primula* and *Hottonia* (2) gave origin to *GLO^T^* and *CYP^T^*, which were then translocated to the same region, forming the core genes of the hemizygous *S*-locus (red block). In *Hottonia*, the *S*-locus then expanded, incorporating other heterozygous genes (3), while in *H. inflata* the loss of *CYP^T^* and *GLO^T^* led to the loss of distyly (4). The genomic location of the *S*-locus differs among *P. veris*, *P. edelbergii*, and *Hottonia* species, implying at least two translocation events (5). Conversely, in *Androsace*, *CYP734* was duplicated along *CSE* and *EH* in a WGD that occurred in the common ancestor of *A. wulfeniana* and *A. vitaliana* (6) but only in in the latter species these genes neo-functionalized (here symbolized by a bolt), acquiring a role in controlling distyly (7).

## Materials and Methods

### Material collection

The pin and thrum plants used to generate the *Androsace vitaliana* subsp. *lepontina* reference genomes were collected by the Gibidumsee lake of Visperterminen in Canton Wallis (coordinates 46.2577,7.9396) and voucher specimens were deposited at the Zurich University Herbarium (accession number: *Z-000227551*). An additional 24 accessions (comprising 12 pins and 12 thrums) were collected from the Visperterminen and Zermatt areas in Canton Wallis (accession numbers: *Z-000227549*, *Z-000227550*, *Z-000227552*) and 14 accessions (7 pins and 7 thrums) sampled directly from herbarium specimens of the Zurich University Herbarium and Real Jardín Botánico Madrid Herbarium collected in the French Alps (pin & thrum: *Z-000195747*), Pyrenees (pin: *MA 320297*, thrum: *MA 320297*), Cantabrian (pin: *MA 493665*, thrum: *MA 493665*), Spanish Central System (pin: *MA 532234*, thrum: *MA 560365*), Nevada Mountains (pin: *MA 888351*, thrum: *MA 889193*), Montes Aquilanos (pin: *MA 280017*, thrum: *MA 280016*), the Apennines (pin: *MA 698758*) and Dolomites (thrum: *MA 353134*) for Whole Genome Resequencing (WGS) with Illumina (see below). For RNA sequencing, we collected leaves and flowers in RNAlater (ThermoFisher) from the Canton Wallis population.

The plant used to generate the *Hottonia palustris* reference genome assembly was collected in the Burgwies pond in Zurich (coordinates 47.3569,8.5747), and documented on iNaturalist (https://www.inaturalist.org/observations/41758573). An additional 28 accessions (comprising 14 pins and 14 thrums) were collected in Switzerland, France and Germany for WGS with Illumina (see below).

The *Hottonia inflata* individual used to generate the reference genome assembly was collected from Carbondale, Illinois, USA (specimen deposited at the Marie-Victorin Herbarium, *Lacroix-Carignan 1467*, MT) and shipped to Zurich for HiFi and RNA sequencing and to Arima Genomics (California, USA) for Hi-C sequencing. For RNA sequencing, we used leaves, buds, and flowers that were collected in RNAlater (ThermoFisher) or fresh material.

### DNA extraction and sequencing

To generate the genome assembly of each species, DNA was extracted from fresh leaves using a modified CTAB protocol, specific for high molecular weight DNA isolation (*26*). DNA sequencing was then performed with Oxford Nanopore Technologies (ONT) and Illumina platforms for *Androsace* species and PacBio HiFi for *Hottonia* species.

For *Androsace* species, ONT libraries were prepared using the SQK-LSK108 kit and sequenced on MinION and PromethION R9 flow cells for 48-72h to achieve a minimum of 60× coverage. Basecalling was done using the methylation aware (MinION) and high accuracy (PromethION) models in Guppy v.3.3.3. In addition, ca. 110× of Illumina 150 bp paired-end (PE) reads (300 bp insert size) was generated on a NovaSeq 6000 for the *Androsace* species. PacBio HiFi sequencing was done using the Sequencer PacBio II and PacBio IIe for *H. palustris* and *H. inflata*, respectively, at the Functional Genomics Center Zurich (FGCZ), Switzerland. For WGS data generation, DNA was extracted from dried samples or herbarium specimens (for *A. vitaliana*) using a modified CTAB protocol (*61*). TrueSeq Illumina libraries were generated and sequenced for PE 150 bp on a NovaSeq platform at the FGCZ.

Chromatin-conformation capture methods were used to aid genome scaffolding. Specifically, for *Androsace* genomes, Hi-C libraries were generated using a previously-published protocol (*62*) and sequenced on an Illumina NovaSeq 6000; for *H. palustris,* Omni-C® libraries were prepared using the Omni-C® library preparation kit (Dovetail, USA), loaded for paired-end sequencing on an Illumina NovaSeq 6000 system and run in XP mode using a NovaSeq 6000 SP Reagent Kits v1.5 (300 cycles). We set a single index running mode to 6:151:151:0 cycles. For *H. inflata*, Hi-C libraries were prepared using the Arima library preparation kit (Arima Genomics, USA) and sequenced on an Illumina NovaSeq.

Statistics on short- and long-read sequencing data were obtained with SeqKit (*63*) v2.9.0 and NanoStat (*64*) v1.1.2, respectively.

### RNA extraction and sequencing

For *H. palustris*, we extracted RNA from leaves, buds (young and old) and flowers using the Spectrum Plant Total RNA Kit (Sigma-Aldrich). TrueSeq Stranded mRNA libraries were generated for each sample and sequenced on a NovaSeq 6000 at the Functional Genomics Center Zurich (FGCZ, Switzerland) to generate 20 M 150-bp PE reads per sample.

RNA was isolated from vegetative (leaves and stem) and reproductive (flowers and flower buds) tissue of *Androsace vitaliana* pins and thrums, vegetative tissue of *A. wulfeniana*, as well as vegetative and reproductive tissue of *A. septentrionalis* with a Spectrum Plant Total RNA Kit (Sigma-Aldrich), and RNA integrity was checked on a TapeStation 6000 (Agilent Technologies). TruSeq Stranded mRNA libraries were prepared and sequenced on an Illumina NovaSeq 6000 to generate 100 M 150-bp PE reads per sample.

### Genome profiling

We estimated genome sizes using both a *k*-mer based genome profiling and flow cytometry in order to assess whether our genome assembly sizes were close to the expected size.

Genome profiling (i.e. estimate of genome size, repeat content, heterozygosity and haplotype length) was performed via *k*-mer analysis on Illumina reads for *Androsace* and PacBio HiFi reads for *Hottonia*. First, *k*-mers (31-mers for *Androsace* and 21-mers for *Hottonia*) were counted with the *count* function of Jellyfish (*65*) v2.2.10 (-C, -m 21). Then we used the *histo* function of Jellyfish to generate a suitable input file for the online version of GenomeScope (*66*) (qb.cshl.edu/genomescope) with the following parameters: for *Androsace*, *k*-mer length=31; read length=150; max *k*-mer coverage=10,000; for *Hottonia*, *k*-mer length=21; read length=10,000; max *k*-mer coverage=10,000.

To verify the results obtained with this *k*-mer-based approach, we also estimated genome sizes of *Androsace vitaliana*, *A. wulfeniana*, *A. septentrionalis* and *Hottonia palustris* with flow cytometry using 1-4 individuals per species (**Table S1**), following a previously-published protocol (*67*). Briefly, fresh leaf material of each sample was co-chopped (*68*) with a reference (*Raphanus sativus* cv. ‘Saxa’ 1C=1.11 pg (*69*), *Solanum lycopersicum* cv. ‘Stupicke polni tyckove rane’ 2C=1.96 pg (*69*), or *Solanum pseudocapsicum* 2C=2.59 pg (*70*)) in Otto I buffer, the suspension filtrated, digested with RNase, mixed with Otto II buffer, and stained with propidium iodide in the dark at 4°C for 1–24 hours. At least 10,000 nuclei were analyzed on a Cytoflex S (Beckman Coulter), or a CyFlow ML (Partec; green laser 100mW, 532nm, Cobolt Samba, Cobolt AB) flow cytometer. Only nuclei peaks with coefficients of variation below 5% were analyzed.

### Genome assembly

*Androsace* assemblies were generated using both Illumina and nanopore reads and the hybrid approach in MaSuRCA (*71*) v3.4.1 with the longest 35× raw ONT reads and all the 110× Illumina reads, polished with one round of POLCA (*72*), followed by one round of Pilon (*73*) v.1.23, and by mapping the trimmed Illumina reads with BWA-MEM (*74*) v.0.7.17. Organellar scaffolds were identified with tBLASTn (*75*) (BLAST v2.8.1; -evalue 1e-25, -max_target_seqs 1) using the chloroplast proteome of *Primula veris* (Genbank accession: KX639823) and mitochondrial proteome of *Camellia sinensis* (Genbank accession: NC_043914.1) as queries, and contigs with >12 significant hits to plastid or mitochondrial genes were excluded from the main assembly. Allelic contigs (haplotigs) were identified and removed with Purge_dups (*76*) v1.2.3 using default settings except for a similarity threshold of 94%, minimum fraction of 70%, and manual coverage cutoffs determined from the coverage histogram. After removing contaminant, organellar and allelic contigs, the remaining contigs were scaffolded with Hi-C short reads with Juicer (*77*) v.1.5.7, 3d-dna (*78*) v180922 and HiC-Hiker (*79*) v.1.0.0, followed by manual adjustments in juicer v1.1 (Juicebox Assembly Tools; JBAT) (*80*). Finally, gaps in the assembly were closed with TGS-GapCloser (*81*) v.1.1.1 using uncorrected nanopore reads, polished with one round of Racon (*82*) v.1.4.3 and two rounds of Pilon (*73*) v.1.23.

During the Hi-C scaffolding phase of the *Androsace vitaliana* thrum genome assembly, an unscaffolded contig was identified as containing the *S*-alleles, based on the *k*-mer, *F_ST_*, and morph-biased heterozygosity analyses (see *S*-locus identification section). Hi-C contact maps further confirmed that this contig was positioned in the same genomic region where the *s*-alleles had been scaffolded on chromosome 5. Given that the *S*-alleles are thrum-specific, the contig containing them was manually placed in the corresponding position on chromosome 5, replacing the *s*-alleles also found in the pin genome assembly. This adjustment ensured that the reference *A. vitaliana* thrum genome assembly contained the *S*-alleles in its chromosome-scale scaffolds. The junction between the *S*-allele contig and adjoining contigs was subsequently gap-filled and polished to ensure assembly continuity. The *k*-mer, *F_ST_* and morph-biased heterozygosity results presented here are based on this final reference genome.

The *H. palustris* genome assembly was generated by combining PacBio HiFi long reads and Omni-C® data in HiFiasm (*83*) v0.16.1, resulting in a haplotype-phased assembly. Each haplotype was then scaffolded using Hi-C reads in YaHS (*84*) v1.2. The *H. inflata* assembly was generated using PacBio HiFi long reads in HiFiasm (*83*) v0.16.1 and scaffolded using Hi-C reads in YaHS (*84*) v1.2. A Hi-C contact map was then generated for each assembly and visualized it in JBAT (*80*), allowing for the manual curation of misassemblies.

Before scaffolding, each assembly was screened for contaminant sequences using Blobtools (*85*) v1.1.1 in combination with the UniProt database (*86*) (The UniProt Consortium 2017), removing few (<1% of assembly length) contigs.. Contigs representing contaminants were removed. Telomeric repeats were identified in each genome independently with the *TeloExplorer* tool of quarTeT (*87*) v1.2.0, which searched for the TTTAGGG monomer repeated in tandem at least 50 times (-c plant, -m 50). The completeness of the assemblies was assessed with BUSCO (*88*) v.5.6.1 (-m genome), using the 2,326 single-copy orthologs from the eudicot database (eudicots_odb10; creation date: 2024-01-08), while basic statistics on the assemblies were obtained with Quast (*89*) v5.0.2.

### Transposable element annotation

Repetitive elements were identified in all assemblies using EDTA (*90*) (v1.8.3 and v1.9.4 for *Androsace* and *Hottonia* genome assemblies, respectively), which combines structure- and homology-based approaches for *de novo* TE identification. Structural discovery of TEs was achieved using LTRharvest (*91*) and LTR_retriever (*92*) for LTR retrotransposons, TIR-Learner (*93*) for TIR transposons, and Helitronscanner (*94*) for helitrons, generating a refined, non-redundant TE library for each genome assembly. Additional repetitive sequences were identified using RECON (*95*) v1.08 and RepeatScout (*96*) v1.06 through RepeatModeler (*97*) v2.0, resulting in a curated TE library specific to each assembly. The *de novo* TE libraries produced by EDTA were then used to annotate the respective assemblies using RepeatMasker (*98*) v4.0.9.

### Gene annotation

Gene annotation was conducted individually for each assembly by using a combination of *ab initio* and evidence-based methods and supported by using both protein datasets and RNA-seq data. For *Androsace*, GeneMark (*99*) v.4 and AUGUSTUS (*100*) v.3.3.3 were trained for *de novo* gene prediction. To achieve this, RNA-seq reads were mapped to the assemblies (soft-masked with the *maskfasta* function of BEDtools (*101*) v2.28.0 [-soft]) using Hisat2 (*102*, *103*) v2.1.0 (--phred33, --very-sensitive, --max-intronlen 50000) and *ab initio* gene predictors were trained using BRAKER (*104*) v.2.1.5 in *etp* mode (development version). The MAKER (*105*) pipeline was then employed to integrate the *ab initio* gene predictions, along with evidence from SwissProt Viridiplantae protein sequences and a transcriptome assembly generated for each species with Trinity (*106*) v2.11.0 to improve gene annotations. In the first round of MAKER, genes were predicted with five sources of evidence, while soft-masking the genome with the species-specific TE library: (1) the Trinity transcriptome assembly, (2) Swissprot proteins, (3) the gff3 obtained with BRAKER, (3) the GeneMark HMM file, and (3) trained Augustus gene modes. The evidence alignments were turned into ‘hints’ and MAKER v3.01.03 was run iteratively two additional times using the transcriptomes as evidence. The resulting set of predicted genes were annotated with Pfam domains (*107*) using InterProScan (*108*) and the models were filtered selecting them if they had an annotation edit distance < 1 and/or PFAM domain. For *Hottonia*, GeneMark (*99*) v.4 and AUGUSTUS (*100*) v.3.3.3 were trained for *de novo* gene prediction using RNA-seq data mapped onto the assemblies (soft-masked with RepeatMasker (*98*)) using Hisat2 (*102*, *103*) v2.1.0 (--dta, --max-intronlen 100000) and ab initio gene predictors were trained using BRAKER (*109*) v3.0.1. Additionally, a set of protein sequences obtained by merging the OrthoDB protein data set for Viridiplantae (odb10; www.orthodb.org) with a high-quality *P. veris* protein set (see ref. (*27*) for details on how the protein data set was obtained) was used as input for homology-based annotation in BRAKER.

The completeness of the gene annotations was assessed with BUSCO (*88*) v.5.6.1 (-m proteins), using the 2,326 single-copy orthologs from the eudicot database (eudicots_odb10; creation date: 2024-01-08). Functional annotation was performed on proteomes with eggNOG-mapper (*110*) v2.1.12 using the eggNOG database (*111*) v5.

### Estimating recombination rate in *Hottonia palustris*

To investigate if the *S*-locus lies in a region of suppressed recombination in *H. palustris*, we estimated recombination rate along chromosome 9 using ReLERNN (*113*). This software employs recurrent neural networks to estimate recombination rates across genomic windows. ReLERNN v1.0.0 was run with the simulate, train, predict and bs-correct modules (default settings), using the resequencing data of 28 individuals as input. A mutation rate of 7 × 10^−9^ ^(^Ossowski et al. 2010) and generation time of two years were used. The final output contained recombination rate estimates in 40-kb windows; the window size was determined by ReLERNN, ensuring that each window contained between 50 and 1,750 SNPs.

### *S*-locus identification

We identified the *S*-loci of *A. vitaliana* and *H. palustris* using three different approaches. First, we identified morph-specific a *k*-mers and mapped them on the genome assemblies. Second, we used a population genomic approach to search for regions of high differentiation between individuals of different floral morphs. Both these approaches were adopted using WGS data from 28 individuals for *H. palustris*, while for *A. vitaliana* these analyses were run separately on 12 herbarium specimens and 24 individuals from populations of Canton Wallis. In each data set, pins and thrums were equally represented. Third, we mapped the *P. veris S*-genes on the genome assemblies of *H. palustris* and *A. vitaliana* to test whether homologs of these genes were contained in the newly identified *S*-loci.

*Identification of morph-specific k*-mer*s.* To identify the *S*-locus in *H. palustris* and *A. vitaliana*, we utilized a *k*-mer-based approach to detect morph-specific *k*-mers. This method was chosen because it does not rely on prior knowledge of morph genotypes and is sensitive to both SNPs and structural variation. First, 31-mers were counted in each sample using the *count* function in Jellyfish (*65*) v2.2.10. The resulting *k*-mers were filtered to retain only those with coverage between 2 and 240, thereby excluding *k*-mers likely arising from sequencing errors (low coverage) or repetitive regions (high coverage). Additionally, *k*-mers were retained only if they occurred in at least two samples. Finally, a *k*-mer was defined as morph-specific if it was present in *n*-2 samples of the corresponding morph (where *n* is the total number of samples for that morph) and none of the samples of the other morph. Morph-specific *k*-mers identified in *H. palustris* and *A. vitaliana* were then mapped on the thrum haplotype assembly of the respective species with BWA-MEM (*74*) v0.7.17. The resulting BAM files were sorted with SAMtools (*112*) v1.9-63 and coverage was calculated in 5-kb windows with Mosdepth (*113*) (--by 5000, --no-per-base).

*Population genomics analyses.* To complement the approach based on morph-specific *k*-mers aimed at identifying the *S*-locus of *A. vitaliana* and *H. palustris* (see above), we searched for genomic regions characterized by high divergence between pin and thrum samples and by a higher heterozygosity in thrums compared to pins. For both species, reads were aligned to the chromosome-scale scaffolds of the thrum-specific haplotype with BWA-MEM (*74*) v0.7.17. The resulting BAM files were sorted with SAMtools (*112*) v1.9-63 and duplicate reads were removed with the *MarkDuplicates* function of Picard v2.18.14 (http://broadinstitute.github.io/picard/) (REMOVE_DUPLICATES = true, ASSUME_SORTED = true, VALIDATION_STRINGENCY = SILENT). Variant calling was performed with the *mpileup* and *call* functions of Bcftools (*112*, *114*) v1.8. The resulting VCF files were filtered with VCFtools (*115*) v0.1.17 in order to keep only sites that represented biallelic SNPs, with a minimum quality of 30, present in at least 25% of samples (--remove-indels, --max-missing 0.25, --minQ 30, -- max-alleles 2), and with a sequencing coverage between 5 and 70 for *H. palustris* (--min-meanDP 5, --max-meanDP 70, --min-meanDP 5, --max-meanDP 70) and between 2 and 70 for *A. vitaliana* (--min-meanDP 2, -- max-meanDP 70, --min-meanDP 2, --max-meanDP 70). Fixation index (*F_ST_*) was calculated between pins and thrums in 5-kb non-overlapping windows using *popgenWindows.py* (https://github.com/simonhmartin/genomics_general), allowing for a minimum of 100 sites per window (-w 5000, -m 100, --writeFailedWindows). To compare heterozygosity between pins and thrums, we first estimated heterozygosity in 5-kb non-overlapping windows for each individual using *popgenWindows.py*, allowing for a minimum of 100 sites per window (-w 5000, -m 100, --writeFailedWindows, --analysis indHet). Secondly, we calculated the mean heterozygosity for pins and for thrums for each window. Finally, we subtracted the average heterozygosity in pins from the average heterozygosity in thrums for each window.

*Mapping of Primula S-genes*. To determine whether the *S*-loci of *H. palustris* and *A. vitaliana* evolved independently from the *P. veris S*-locus, we investigated whether these newly-identified *S*-loci contained homologs of the *P. veris S*-genes. We therefore mapped the amino acid sequences of the *P. veris S*-genes (*26*) on the *H. palustris* and *A. vitaliana* proteomes using BLASTp (*75*) (-evalue 1e-5).

The final coordinates of the *S*-loci are: in *A. vitaliana*, the *s*-haplotype is located at aviP_sc1:15,984,083-16,018,688 bp and the *S*-haplotype at aviT_sc5:984,089-1,039,657 bp; in *H. palustris*, the s-haplotype is located at Hpal_hap1_9:36,917,167-53,577,764 bp and the *S*-haplotype at Hpal_hap2_9:36,762,381-49,532,255 bp; in *H. inflata*, the region syntenic to the *S*-locus is located at Hinf010:26,847,990-37,146,126 bp.

### Differential gene expression analysis in *Androsace vitaliana*

To quantify gene expression in *A. vitaliana*, we used 36 RNA-seq samples comprising leaves and flowers from nine thrum and nine pin individuals, and a gene set created by concatenating the CDS of the pin and thrum haplotypes, containing a total of 102,726 CDS. Reads were first trimmed with Trimmomatic (*116*) v0.38, with the parameters recommended by Trinity (*106*) (ILLUMINACLIP:2:30:10, LEADING:5, TRAILING:5, SLIDINGWINDOW:4:5, MINLEN:25). The CDS file was indexed with the *index* function of Salmon (*117*) v1.4.0. The *quant* function of Salmon was then used to quantify gene expression (--gcBias --validateMappings). Read counts obtained with Salmon for leaf and flower samples were imported separately into R v4.3.3 (https://www.R-project.org/) using *tximport* (*118*) and a DESeqDataSet was created with the *DESeqDataSetFromTximport* function of the DESeq2 (*119*) v1.42.1 R/Bioconductor (*120*) package. Read counts were normalized using the default median of ratios method (*119*) and plotted for the *s*- and *S*-alleles of *AvCSE*, *AvEH*, and *AvCYP*.

### Synteny analyses

Syntenic genes were identified within each species and between each species pair using the MCScan tool of the JCVI toolkit (*121*, *122*) v1.3.6, following the GitHub manual (github.com/tanghaibao/jcvi/wiki/MCscan-(Python-version)). First, homologous genes were identified with *jcvi.compara.catalog ortholog* (--min_size=5, --dist=20, --no_strip_names) and the resulting anchor files were used to generate whole-genome dot plots with *jcvi.graphics.dotplot*. Then, simplified anchor files were generated with *jcvi.compara.synteny screen* (--minspan=30, --simple) and used to create whole-genome macrosynteny plots with *jcvi.graphics.karyotype*. Finally, we generated a list of syntenic genes with *jcvi.compara.synteny mcscan* (--iter=1), which was used to: a) generate microsynteny plots; b) estimate d_S_ between syntenic genes within and between species to estimate WGDs in *Androsace*; c) estimate d_S_ between the *S*- and *s*-haplotype of *H. palustris*.

### Identification of WGDs in *Androsace*

To identify WGDs in *Androsace* species, we estimated d_S_ between syntenic genes within and between *A. vitaliana* (thrum haplotype; 2n=4x=40), *A. wulfeniana (*2n=4x=40), and *A. septentrionalis (*2n=2x=20). For each estimate, syntenic genes were identified using MCScanX as outlined above. Then, d_S_ values were estimated using ParaAT (*123*) v2.0, which employs MUSCLE (*124*) v3.8.31 to align sequences and KaKs_Calculator (*125*) v2.0 to calculate d_S_. The resulting d_S_ distributions were then plotted in R v4.3.3 using the ggplot2 package. Additionally, we estimated the syntenic depth within and between the abovementioned *Androsace* genomes using the *jcvi.compara.synteny depth* function of MCScanX.

### Calculating DoS and performing MK test

To estimate whether heterozygous and hemizygous *S*-genes are undergoing different evolutionary trajectories, we calculated the direction of selection (*126*) (DoS) and performed a McDonald-Kreitman (*127*) (MK) test on *H. palustris*, *A. vitaliana*, and *P. veris* using the software *degenotate.py* (https://github.com/harvardinformatics/degenotate). This tool requires whole-genome sequencing (WGS) data from multiple individuals of the focal species, an outgroup, and the reference genome and gene annotation of the focal species as input.

For all species, reads were aligned to the chromosome-scale scaffolds of the thrum-specific haplotype with BWA-MEM (*74*) v0.7.17. The resulting BAM files were sorted with SAMtools (*112*) v1.9-63 and duplicate reads were removed with the *MarkDuplicates* function of Picard v2.18.14 (http://broadinstitute.github.io/picard/) (REMOVE_DUPLICATES = true, ASSUME_SORTED = true, VALIDATION_STRINGENCY = SILENT). Variant calling was performed with the *mpileup* and *call* functions of Bcftools (*112*, *114*) v1.8. The resulting VCF files were filtered with VCFtools (*115*) v0.1.17 in order to keep only sites that represented biallelic SNPs, with a minimum quality of 30 that did not overlap with repetitive elements (--remove-indels, --minQ 30, --max-alleles 2, --exclude-bed), and with a sequencing coverage below 200 for *H. palustris* and *A. vitaliana* (--max-meanDP 200, --max-meanDP 200), and between 5 and 70 for *P. veris* (--min-meanDP 5, --max-meanDP 70, --min-meanDP 5, --max-meanDP 70)

For *H. palustris*, the VCF file used as input for *degenotate.py* was generated by mapping sequencing reads from 28 *H. palustris* and 20 *P. veris* samples to the *H. palustris* genome assembly (pin haplotype). To estimate selection on heterozygous *S*-genes, we analyzed *S*- and *s*-alleles separately. For *s*-alleles, thrum *H. palustris* samples were removed from the VCF prior to running *degenotate.py*. For *S*-alleles, alleles fixed in *H. palustris* pins – i.e. alleles homozygous in all pin individuals, presumably representing alleles fixed in the *s*-haplotype – were removed from the VCF; this way, heterozygous alleles in thrums were converted into haploid alleles, likely representing *S*-alleles. Selection on hemizygous *S*-genes was assessed using a separate VCF created by aligning the same set of samples to the *H. palustris* thrum haplotype assembly. For *A. vitaliana*, we applied a similar approach using 14 herbarium samples of *A. vitaliana* along with the individuals used to generate the pin and thrum genome assemblies. As neither *P. veris* nor *H. palustris* contains orthologs of all three *A. vitaliana S*-genes (*P. veris* lacks the *AvCYP* ortholog, and *H. palustris* lacks the *AvEH* ortholog), we performed two separate analyses using *P. veris* (10 samples) and *H. palustris* (14 samples) as outgroups. For *P. veris*, we used 20 *P. veris* samples and 20 *P. vulgaris* samples, mapping them to the *P. veris* thrum haplotype assembly. Since all *P. veris S*-genes are hemizygous, there was no need to separate *S*- and *s*-alleles for this species.

### Testing for TE enrichment in *S*-loci

We tested whether S-loci of *A. vitaliana*, *H. palustris*, and *P. veris* were enriched in TEs compared to the genomic background and to their flanking regions. To assess whether *S*-loci were enriched in TEs compared to the genomic background, for each species we compared the *S*-locus TE content to that of 10,000 randomly sampled genomic windows across the genome of the same size as the *S*-locus, i.e. 34,605 bp for *A. vitaliana* (*s*-haplotype), 55,568 bp for *A. vitaliana* (*S*-haplotype), 16,660,597 bp for *H. palustris (s*-haplotype), 12,769,874 bp for *H. palustris (S*-haplotype), and 262,809 bp for *P. veris*. For each comparison, an empirical p-value was calculated as the proportion of random windows with TE content equal to or greater than that of the *S*-locus.

Since TE abundance may vary widely along the genome, we also tested whether *S*-loci were enriched in TEs compared to their flanking regions. For each species, we first defined *S*-flanking regions as regions of identical size upstream and downstream the *S*-locus, so that the sum of their lengths equaled the length of the *S*-locus: *A. vitaliana (s*-haplotype), aviP_sc1:15,966,780-15,984,082 bp and 16,018,689-16,035,991 bp; *A. vitaliana (S*-haplotype), aviT_sc5:956,305-984,088 bp and aviT_sc5: 1,039,658-1,067,441 bp; *H. palustris* (*s*-haplotype), Hpal_hap1_9:28,586,868-36,917,166 bp and Hpal_hap1_9: 53,577,765-61,908,063 bp; *H. palustris* (*S*-haplotype), Hpal_hap2_9:30,377,444-36,762,380 bp and Hpal_hap2_9:49,532,256-55,917,192 bp. Second, we generated BED files containing the coordinates of non-overlapping windows of species-specific sizes: 3 kb for *A. vitaliana*, 250 kb for *H. palustris*, and 20 kb for *P. veris*. Third, we estimated TE abundance in each window by using the *coverage* function of BEDtools (*101*) v2.28.0 on the TE annotation generated with RepeatMasker. Finally, the TE content of non-overlapping windows within each *S*-locus was compared to the windows from its flanking regions using a Wilcoxon rank-sum test in R v4.3.3 (https://www.R-project.org/).

### Analyses on *S*-locus expansion in *Hottonia palustris*

To test whether the large size of the *S*-locus in *H. palustris* was explained by an expansion of recombination suppression in *H. palustris* or the *S*-locus was originally larger and ‘shrank’ in *Primula*, we estimated d_S_ between syntenic orthologs of *H. palustris* - *P. veris* (n=15,964), *H. palustris* - *H. inflata* (n=18,830), and between the *S*- and *s*-haplotype of *H. palustris* (n=88); orthologous gene pairs and allele pairs were identified with MCScanX as outlined above, and d_S_ values were estimated using ParaAT (*123*) v2.0.

To test whether antagonistic selection was the main driving force behind the expansion of the *H. palustris* S-locus, we searched for evidence of evolutionary strata and for an enrichment of genes with antagonistic effects. To identify evolutionary strata, we plotted the d_S_ values obtained between the *S*- and *s*-alleles for the 88 *S*-genes present in both haplotypes, ordering genes by increasing d_S_ value. We then assessed whether the *S*-locus was enriched in genes showing morph-biased expression between floral morphs, using *P. veris* genes previously identified as morph-biased as proxies for antagonistic genes. We performed a t-test to assess whether the orthogroups containing *H. palustris S*-genes (n = 127) were enriched in antagonistic genes, against all orthogroups containing *H. palustris* genes (n = 14,050). This analysis was conducted separately for each tissue type in which morph-biased genes were identified in *P. veris* (corolla tube, style, whole flower), as well as across all tissues combined.

### Construction of gene families for phylogenetic analysis

Prior to performing phylogenetic analysis, we clustered genes into gene families (orthogroups) using OrthoFinder (*128*) v2.3.11 run with default parameters on 22 proteomes. These proteomes included: seven proteomes presented here (*A. vitaliana* [pin haplotype], *A. vitaliana* [thrum haplotype], *A. septentrionalis*, *A. wulfeniana*, *H. inflata*, *H. palustris* [pin haplotype], *H. palustris* [thrum haplotype]); proteomes from ten Ericales species, representing seven of the 22 Ericales families (*129*), selected for having chromosome-scale genome assemblies (*Actinidia chinensis* (*130*), *Aegiceras corniculatum* (*131*), *Camellia sinensis* (*132*), *Dyospiros oleifera* (*133*), *Gilia yorkii* (*134*), *Primula edelbergii* (*27*), *Primula veris* (*26*), *Rhododendron henanense* (*135*), *Vaccinium darrowii* (*136*), *Vitellaria paradoxa* (*137*)); proteomes from five additional species, selected for having high-quality genome assemblies and gene annotations and being widespread across the angiosperm phylogeny (*Amborella trichopoda* (*138*), *Arabidopsis thaliana* (*139*), *Solanum lycopersicum* (*140*), *Oryza sativa* (*141*), *Vitis vinifera* (*142*)). The final data set contained 733,259 total proteins, 674,132 (91.9%) of which were assigned to 40,194 orthogroups (**Supplementary Table 20)**.

The *S*-genes were contained in the following orthogroups: OG0006534 (*PvCCM*), containing 32 genes; OG0003769 (*PvGLO*), containing 43 genes; OG0000663 (*PvCYP*), containing 92 genes; OG0001331 (*PvPUM*), containing 70 genes; OG0000291 (*PvKFB*), containing 127 genes; OG0008416 (*AvCSE*), containing 28 genes; OG0005520 (*AvEH*), containing 35 genes. To avoid that some *S*-gene-containing orthogroups contained too few genes, thus hindering the phylogenetic analysis, we created “extended orthogroups” by aligning the amino acid sequences of *P. veris* and *A. vitaliana S*-genes against all orthogroup sequences with BLASTp (*75*) (-evalue 1e-25, -max_target_seqs 3) and concatenating the orthogroups having at least one match. The following orthogroups were concatenated: OG0002583, OG0003881, OG0005521, OG0007367, OG0008416, and OG0011811 for *AvCSE* (212 genes); OG0000010, OG0000663, OG0003314, OG0005243, and OG0013733 for *PvCYP* (619 genes); OG0005129, OG0005520, OG0008234, OG0017032, OG0035311, and OG0037733 for *PvPUM* (110 genes); OG0002710, OG0003769, OG0010238, and OG0011028 for *PvGLO* (143 genes); OG0001331, OG0002862,OG0009601, and OG0014179 for *PvEH* (160 genes). No additional orthogroups were found to contain homologs of *PvCCM* and *PvKFB*, other than OG0006534 and OG0000291, respectively. The sequences contained in these extended orthogroups were then used to generate gene phylogenies (see below).

### Phylogenetic analysis

The *S*-gene homologs obtained from genomes and contained in the extended orthogroups were then used to infer phylogenetic relationships within each gene family.

The *PvCCM* homologs obtained from genomes were aligned using MAFFT (*143*) v7.450 Auto algorithm. A preliminary phylogeny was estimated with FastTree (*144*) v2.1.11 using a GTR+G (4 categories) model. This analysis revealed no clear evidence of subdivision into several gene lineages, so all the sequences were aligned with OMM_MACSE (*145*, *146*) v12.01 for use in phylogenetic and dating analyses.

*AvCSE* is part of the larger class I carboxylesterase gene family, which also contains tannase and acetate esterase genes (*147*). One sequence of each of the three subfamilies were obtained from GenBank to serve as references (*CSE* from *Arabidopsis*, tannase from *Camellia*, acetate esterase from *Solanum*). These reference sequences were aligned alongside *AvCSE* homologs obtained from genomes, using MAFFT v7.450 Auto algorithm. A preliminary phylogeny was estimated with FastTree v2.1.11 using a GTR+G (4 categories) model. This analysis revealed that *CSE*, tannase and acetate esterase formed three separate lineages, in addition to six other lineages of class I carboxylesterase genes. We kept only the sequences of the *CSE* lineage, which included the *Androsace S*-locus sequence, while all other sequences were discarded.

Additional mRNA sequences from the *CYP72* clan (*CYP72*, *CYP709*, *CYP735*, *CYP734*, *CYP715*, *CYP721*, *CYP714*, and *CYP749* families), the *CYP86* clan (*CYP704*, *CYP94*, *CYP86*, and *CYP96* families) and the *CYP87* clan (*CYP87* family) of *Arabidopsis thaliana* and *Oryza sativa* were obtained from Genbank or EMBL, using the previously-published guide trees and classification (*148*, *149*). These reference sequences were aligned alongside *CYP* homologs obtained from genomes, using MAFFT v7.450 Auto algorithm. A preliminary phylogeny was estimated with FastTree v2.1.11 using a GTR+G (4 categories) model. Then, a subset of *CYP* homologs were selected to include only genes in the *CYP734* family, discarding all other sequences, except for reference sequences obtained directly from Genbank and EMBL, which were kept as outgroups. The reduced dataset was re-aligned with OMM_MACSE v12.01 for use in phylogenetic and dating analyses.

The *AvEH* homologs obtained from genomes were aligned using MAFFT v7.450 Auto algorithm. A preliminary phylogeny was estimated with FastTree v2.1.11 using a GTR+G (4 categories) model. This analysis revealed a division into three distinct gene lineages, each encompassing sequences from all analyzed species, consistent with the presence of three separate *EH* gene sub-families. Sequences from the two lineages that did not include the *Androsace S*-locus sequence were excluded. The retained sequences were re-aligned with OMM_MACSE v12.01 for use in phylogenetic and dating analyses.

Additional reference sequences of *GLO*/*PI* and *AP3*/*TM6*/*DEF* genes of *Arabidopsis thaliana* and *Solanum lycopersicum* were obtained from Genbank. These reference sequences were aligned alongside *GLO* homologs obtained from genomes, using MAFFT v7.450 Auto algorithm. A preliminary phylogeny was estimated with FastTree v2.1.11 using a GTR+G (4 categories) model. Then, a subset of homologs was selected to include only *GLO*/*PI* and *AP3*/*TM6*/*DEF* genes, discarding all other sequences. The reduced dataset was re-aligned with OMM_MACSE v12.01 for use in phylogenetic and dating analyses.

The *PvKFB* homologs obtained from genomes were aligned using MAFFT v7.450 Auto algorithm. A preliminary phylogeny was estimated with FastTree v2.1.11 using a GTR+G (4 categories) model. The phylogeny revealed an intricate history of lineage-specific duplications, as previously shown (*150*). However, the clade containing *S*-locus sequences of *Hottonia* and *Primula* contained other Ericales sequences placed according to known interspecific relationships. This clade was sister to another clade containing a similar set of Ericales species, suggesting an ancient duplication event shared across Ericales. These two sister clades were selected for further analysis, discarding all other sequences.

Additional mRNA sequences representing the diversity of plant *PUM*/*PUF* genes were obtained from Genbank, using a previously-published guide tree and classification (*151*). These reference sequences were aligned alongside *PvPUM* homologs obtained from genomes, using MAFFT v7.450 Auto algorithm. A preliminary *PUM* phylogeny was estimated with FastTree v2.1.11 using a GTR+G (4 categories) model. This analysis showed *S*-locus sequences to be nested within one of two sister subclades of *PUM* Clade II as previously identified (*151*), this clade showing evidence of a deep duplication shared across all flowering plants. Only the Clade II subclade including the *S*-locus sequences was retained for further analysis, discarding all other sequences.

Protein-coding gene alignments were generated using OMM_MACSE v12.01, which ensures frame-preserving alignments while accounting for insertions, deletions, and sequencing errors. Each gene was aligned independently, using the standard genetic code (code 1) and disabling pre- and post-filtering to ensure that all sequences and nucleotides were retained in the alignments. The dataset was partitioned by codon and the best models and partitioning scheme for each gene alignment was selected by BIC in PartitionFinder (*152*) v2.1.1.

Non-ultrametric phylogenetic trees were estimated with Bayesian inference in MrBayes (*153*) v.3.2.7a using the best models and partitions to avoid the constraints of molecular clock models. Each gene alignment was analyzed with two independent MCMC runs of 10 million generations, sampling every 2000 iterations, with convergence assessed in Tracer (*154*) v.1.7.2, discarding 20% of each run as burnin. The resulting posterior trees were then used as input for AleRax (*155*) to infer reconciled gene trees on the species tree without introducing biases from clock model assumptions at this stage.

For divergence time dating, separate BEAST (*156*) v.2.7.7 analyses were done on each gene family, using the optimal partitioning and models selected by PartitionFinder, but keeping clock and tree models linked (within each gene family). We used a Yule tree prior, with a prior on log-normal prior on birth rate with a standard deviation of 1.175 (to create a 95% highest probability density of about two orders of magnitude around mean) and a mean calculated according to the formula *lambda*=ln (*n*/2)/*t*, where *n* is the number of sequences in the alignment and *t* is the estimated root age. A root age of 139 Mya was used for angiosperms according to ref. (*157*), and *n* was estimated by counting the number of sequences in the largest subclade of each gene family phylogeny that contained a single copy of *Amborella*. Thus, an average birth rate prior was set to 0.0199 for *CCM*, 0.0268 for *CFB*, 0.0190 for *CSE*, 0.0270 for *CYP*, 0.0225 for *EH*, 0.0233 for *GLO*, 0.0327 for *KFB*, and 0.0246 for *PUM*. An optimized relaxed clock model was implemented with a log-normal prior on the rate parameter (ORCucldMean) with mean of 0.00615 substitution/My as previously calculated in *Primula veris* (*26*) and standard-deviation of 0.23 (giving a 95% highest probability density on the mean of 0.00382 to 0.00940 substitutions/My) based on the rate variation in other plants, as previously reported (*158*).

Uniform fossil calibrations were assigned on each gene family phylogeny whenever the phylogenetic relationships estimated by AleRax allowed unambiguous determination that a splitting event was due to speciation, rather than gene duplication within one species. The MRCA constraint in BEAST analyses was enforced as monophyletic only if it received >=95% support in the AleRax gene phylogeny reconciliation analysis. The minimum bound of age calibrations corresponded to the age of the fossil, and the maximum was set to 140 Mya which is the maximum bound for angiosperms estimated by ref. (*157*) using the bracketing method of Marshall (*159*). Stem dates were used instead of crown dates for the minimum calibration of Primulaceae and *Primula* because the species available in our analyses were nested within Primulaceae and *Primula*, and calibrating the crown of the clade recovered in our analyses would thus have overestimated its age. The fossil calibrations we used were: 1) Crown age of angiosperms: 136–140 Mya (angiosperm pollen fossil cited in ref. (*157*)); 2) Stem age of eudicotyledons: 125–140 Mya (tricolpate pollen fossil cited in ref. (*157*)); Crown age of Ericales: 89–140 Mya (flower fossil cited in ref. (*157*)); Stem age of Primulaceae: 66–140 Mya (flower fossil cited in Larson et al. (*160*)); Stem age of Primula: 16-140 Mya (seed fossil cited in Boucher et al. (*161*)). Two separate BEAST dating analyses were run of 50 million generations for each gene, using Coupled MCMC with three heated chains. Results were checked in Tracer v.1.7.2 to ensure convergence of the chains and effective sample sizes >200 for each parameter after discarding 20% of each run as burnin. Maximum clade credibility trees were constructed from the resulting trace files with LogCombiner.

## Acknowledgments

We thank Matthieu Charrier for helping with the sampling of wild *H. palustris* populations in France, Pedro Jiménez-Mejías for help with sampling of herbarium specimens of *A. vitaliana* at the Real Jardín Botánico Madrid Herbarium, and the Functional Genomics Center Zurich (FGCZ) of University of Zurich and ETH Zurich for providing the infrastructures and support in DNA sequencing with PacBio and Illumina platforms. We also thank Alex Bernhard for the flower photographs depicted in Fig. 1.

This study is part of the ERGA (European Reference Genome Atlas) pilot initiative. ERGA Hubs: We thank the Antwerp University Hospital Center of Medical Genetics and Jarl Bastianen for access to sequencing library quality control equipment. Sequencing was supported by the sequencing facility of the Department of Biology, University of Florence through the Departments of Excellence program funded by the Italian Ministry for University and Research. ERGA Commercial Partners: We would like to acknowledge and thank all supplier partners that have kindly donated kits, reagents to the ERGA pilot Library Preparation Hubs, specifically Dovetail Genomics, Part of Cantata Bio LLC (especially Mark Daly, Thomas Swale and Lily Shuie); Arima Genomics; PacBio; Integrated DNA Technologies (IDT); MagBio Genomics Europe GmbH; Zymo Research; Agilent Technologies; Fisher Scientific Spain; Illumina Inc.

We acknowledge financial support from the University of Zurich and the Swiss National Science Foundation (grant no. 175556) to EC, and from an UZH Forschungskredit (FK-19-103), a NSERC Postdoctoral fellowship award (532569–2019), and a NSERC Discovery Grant (RGPIN-2021-03117) to ÉL-B.

## Author contributions

GP, NY, EL-B, and EC conceptualized the study. ÉL-B, NY, BK, HW-S, GS, and MH performed fieldwork. ÉL-B collected herbarium samples. Nucleic acid extraction was performed by ÉL-B and IAG (*Androsace*) and NY (*Hottonia*). EMT performed genome size measurement. ÉL-B prepared libraries for nanopore sequencing. ÉL-B, NY and SG prepared Hi-C libraries. HGL, GD, and HS organized and performed RNA extractions and library preparations for *H. palustris*. HGL, CN, and MAD performed Omni-C® library preparation, sequencing, and demultiplexing. Genomes were assembled by ÉL-B (*Androsace*), NY (*H. palustris*), and NY and IAG (*H. inflata*). Gene annotation was performed by RRC, ÉL-B, and G.P. (*Androsace*), NY (*H. palustris*), and IAG. (*H. inflata*). TE annotation was performed by RRC and ÉL-B. (*Androsace*) and GP (*Hottonia*). ÉL-B wrote the scripts for the identification of morph-specific *k*-mers. Population genomic analyses for the identification of the *S*-locus were performed by ÉL-B, and GP (*Androsace*), and GP and NY *(H. palustris*). IAG estimated recombination rates in *H. palustris*. GP and IAG performed differential gene expression analyses in *A. vitaliana*. ÉL-B performed phylogenetic analyses. GP and EM-C performed analyses on selection. GP performed comparative genomic analyses, wrote the first draft of the manuscript, prepared all figures, and revised the text based on co-author feedback. All authors read, revised and approved the final paper.

## Supplementary Materials

Figures S1-S27 and Tables S1-S20 are provided as supplementary files.

