## Supplementary Figures for "Distinct genomic architectures but the same gene underlie the convergent evolution of a plant supergene"

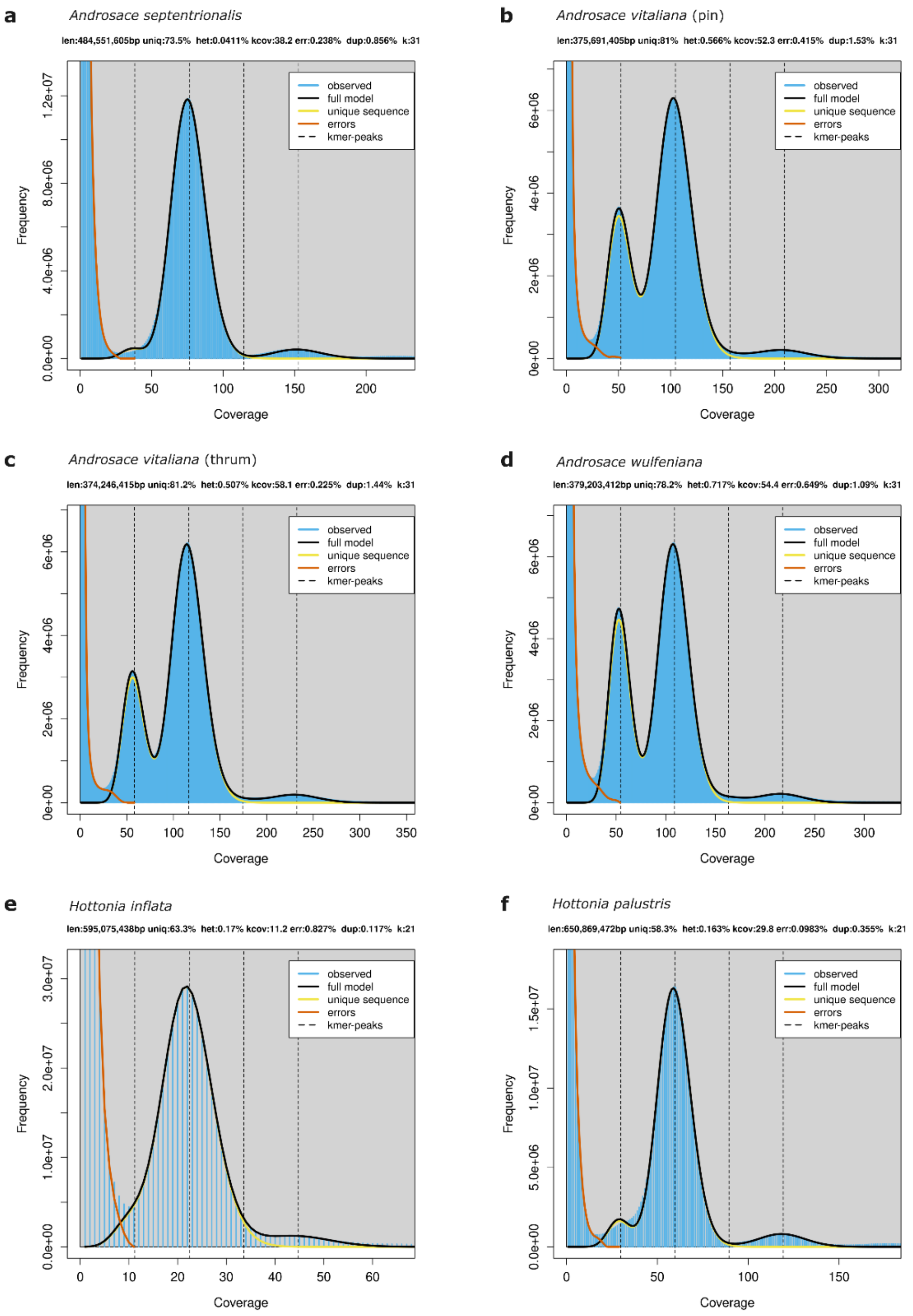

**Fig. S1: Genome profiles obtained with GenomeScope.**
*K*-mer density plots were obtained with GenomeScope using Illumina data and 31-mers for *A. septentrionalis* (**a**), *A. vitaliana* (pin and thrum haplotypes; **b** and **c**, respectively), and *A. wulfeniana* (**d**), and using PacBio HiFi data and 21-mers for *H. inflata* (**e**) and *H. palustris* (**f**).

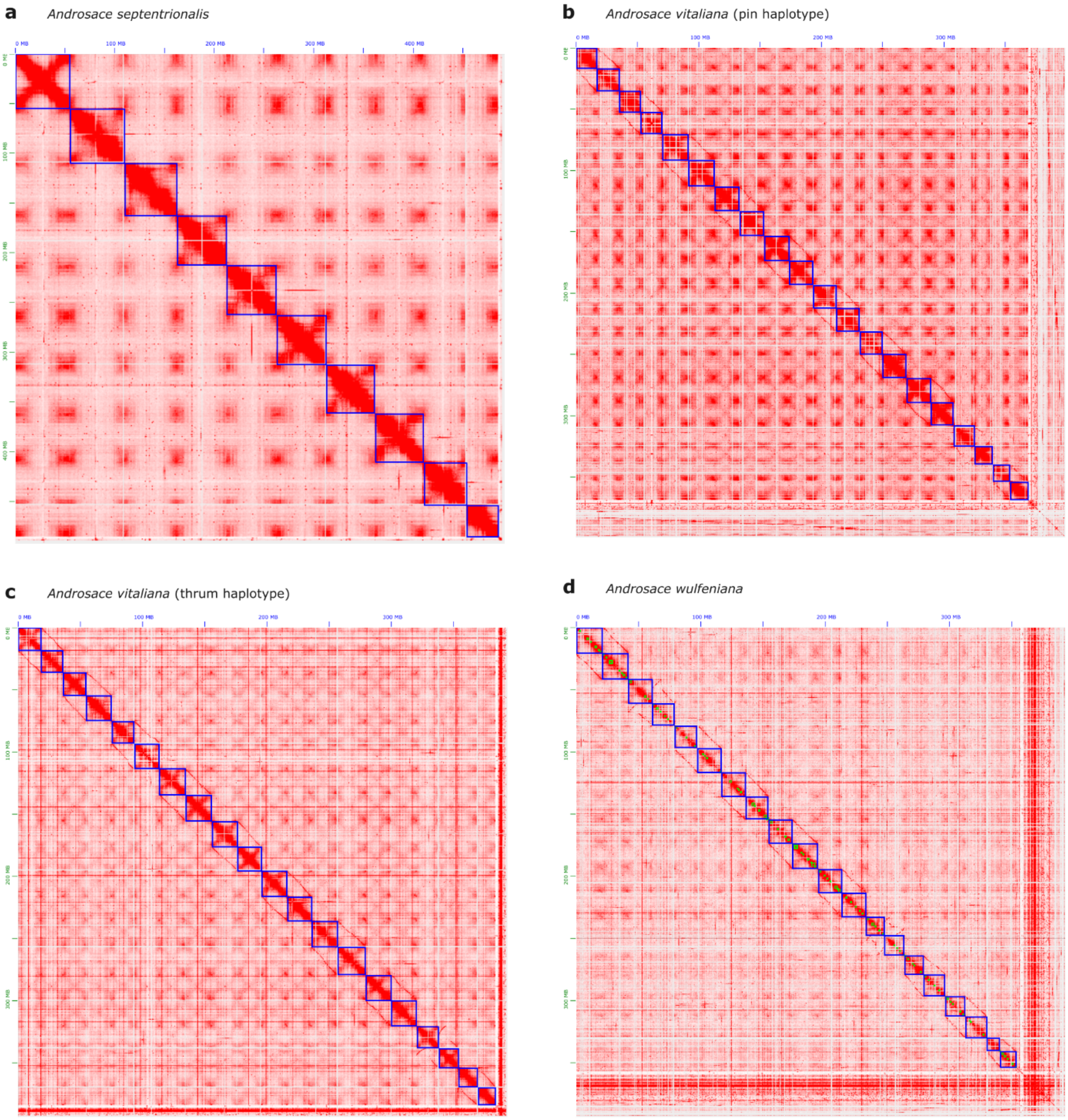
a
Androsace septentrionalis
b
Androsace vitaliana (pin haplotype)
c
Androsace vitaliana (thrum haplotype)
d
Androsace wulfeniana

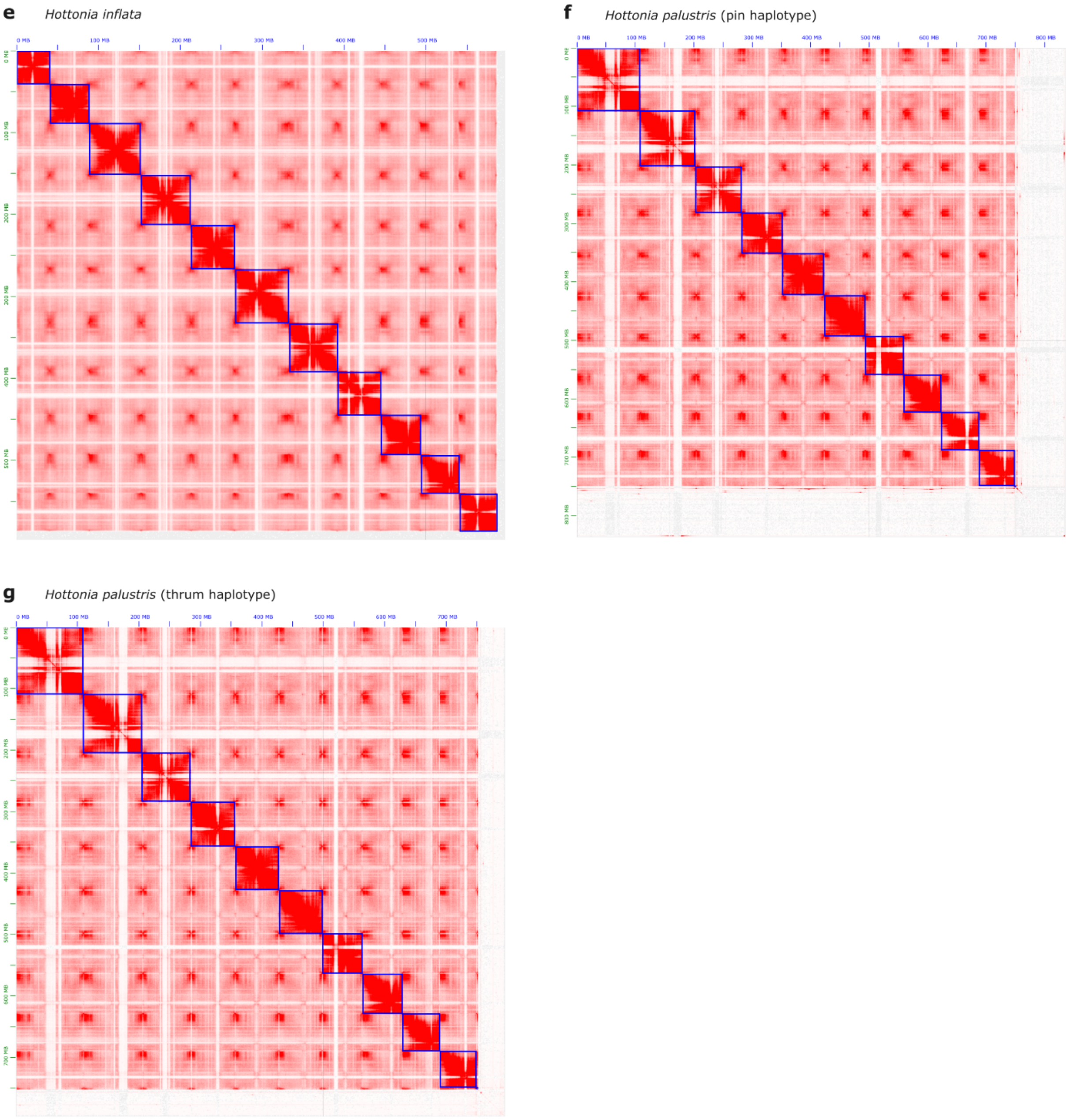

**Fig. S2: Hi-C contact maps for the seven genome assemblies presented here.**
Contact maps of Hi-C libraries for *H. palustris* (pin haplotype; **a**), *H. palustris* (thrum haplotype; **b**), *H. inflata* (**c**), *A. wulfeniana* (**d**), *A. vitaliana* (pin haplotype; **e**), *A. vitaliana* (thrum haplotype; **f**), and *A. septentrionalis* (**g**). Darker colors indicate higher density of chromatin contacts; chromosome-scale scaffolds are delimited by blue boxes.

**Fig. S3:**

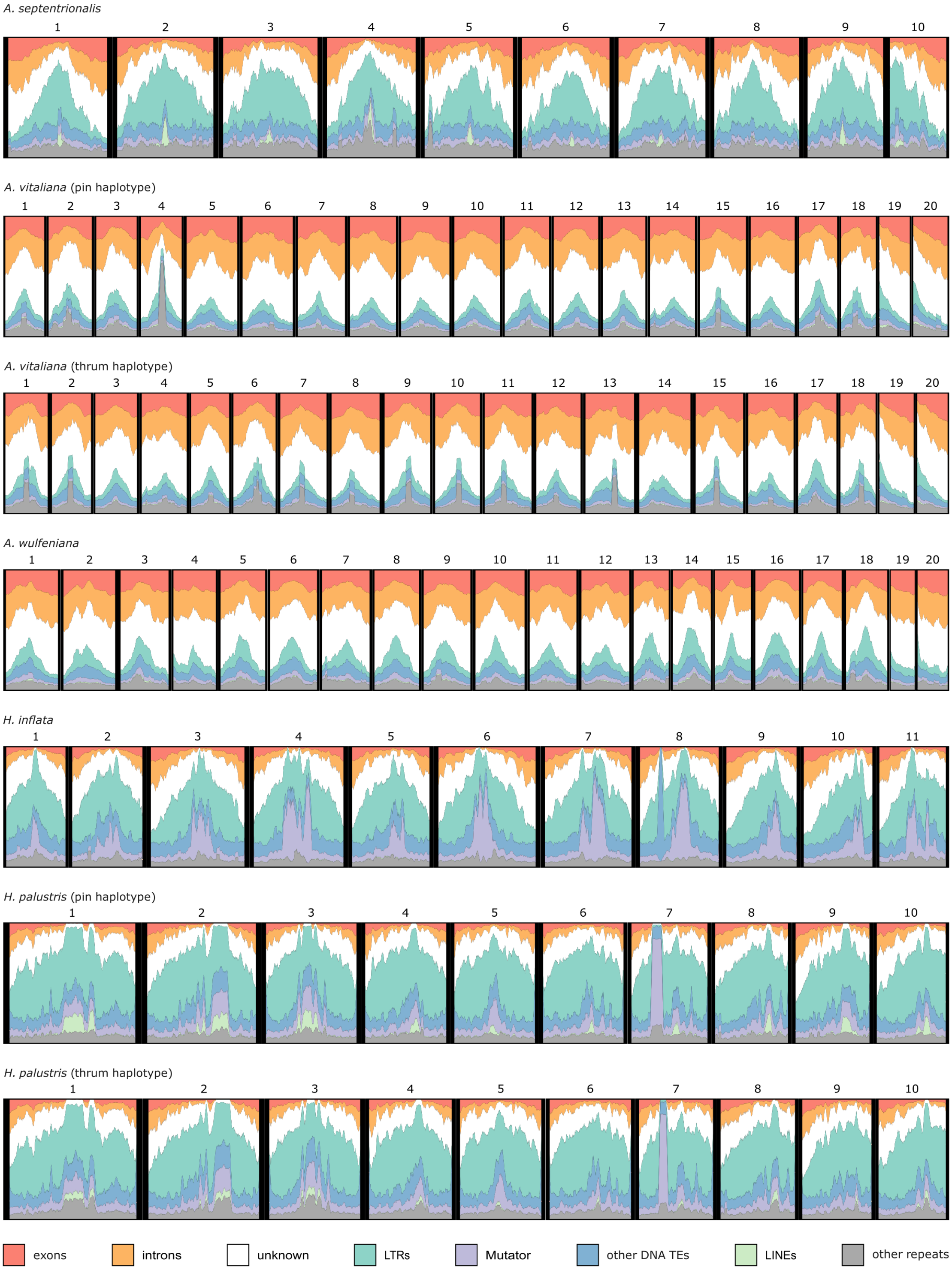

**Overview of the chromosome-scale assemblies presented in this study.**
Distribution of repeat and gene density across the seven genome assemblies (of five species) presented here, calculated in sliding windows (2-Mb width, 100-kb steps). Putative centromeric repeats are characterized by an enrichment of LINEs (light green) in *H. palustris* and *A. septentrionalis*, Mutator (purple) in *H. inflata*, and unclassified TEs (here together with "other repeats", gray) in *A. vitaliana*. The category "other repeats" includes both TEs and tandem repeats.

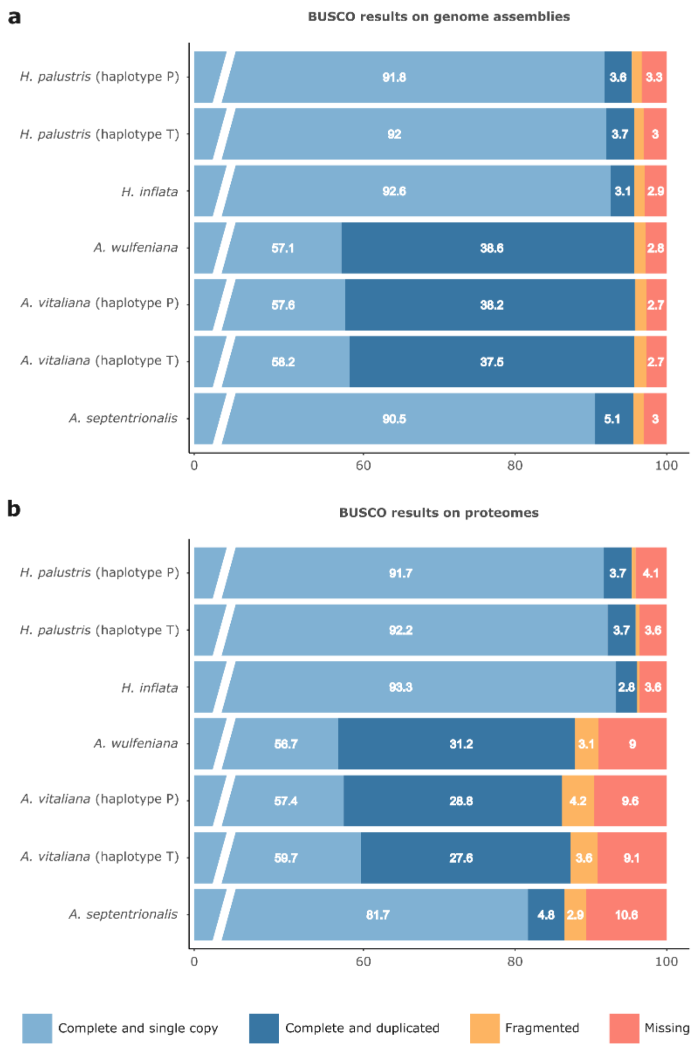

**Fig. S4: Genome and proteome evaluation by BUSCO.**
Completeness BUSCO scores for the genome assemblies (**a**), and respective proteomes (**b**) presented here, obtained using the eudicots_odb10 database, which contains 2,326 single-copy orthologs. Numbers in each bar represent the percentage of genes in each category. To ease visualization, percentages are not reported for fragmented genes that accounted for less than 2% of the total. The total amount of complete BUSCO genes for each species is given by the sum of "Complete and single copy" (light blue) and "Complete and duplicated" (dark blue).

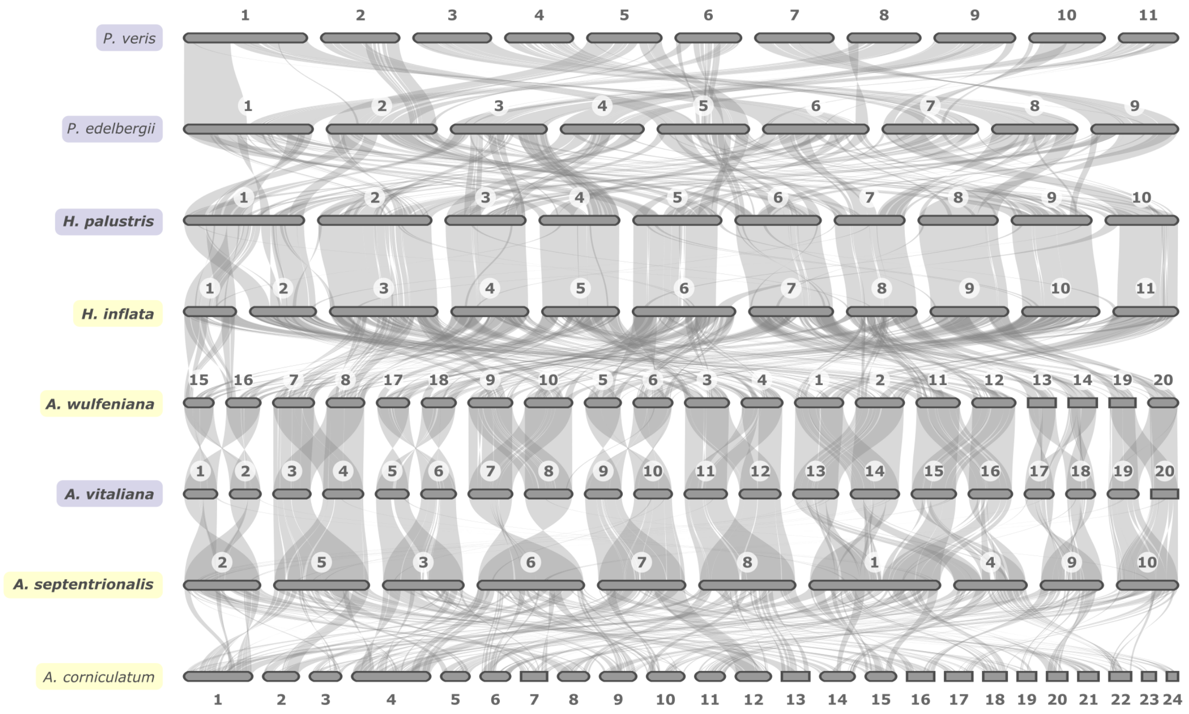

**Fig. S5: Lack of synteny across Primulaceae.**
Whole-genome synteny plots among Primulaceae. Distylous and non-distylous species are highlighted in purple and yellow, respectively; species whose genome assemblies were generated in this study are boldfaced.

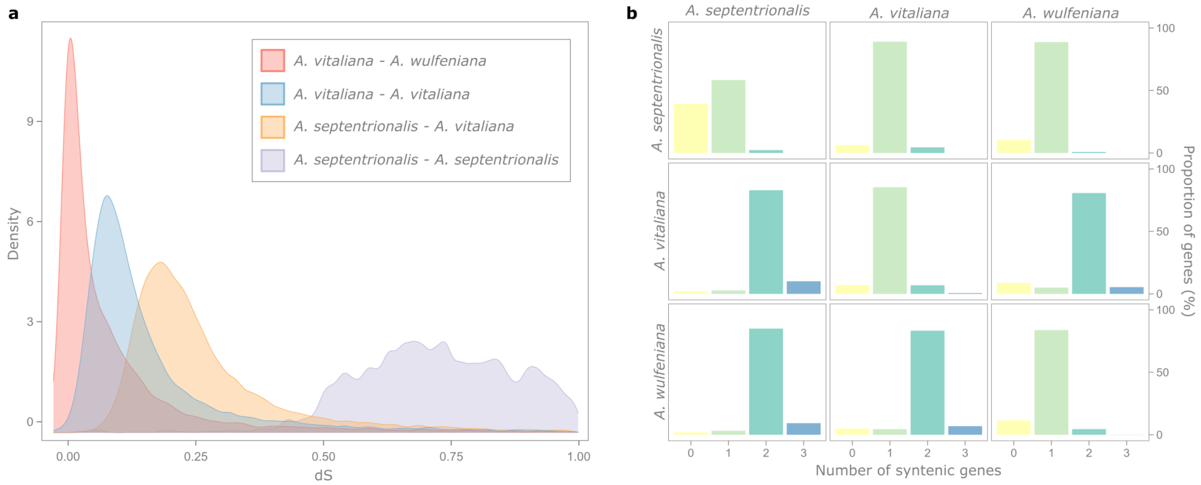

**Fig. S6: Evidence for a WGD shared by *A. vitaliana* and *A. wulfeniana*.**
**a.** Distributions of $d_S$ calculated between syntenic orthologs of *A. vitaliana* - *A. wulfeniana* (n=13,078, calculated only for one subgenome; red curve), *A. septentrionalis* - *A. vitaliana* (n=16,414; orange curve), and between syntenic paralogs within *A. vitaliana* (n=28,899; blue curve) and *A. septentrionalis* (n=5,567; purple curve). The WGD we observe in *A. vitaliana* (blue curve) is shared with *A. wulfeniana* (i.e. it occurred before the divergence between these two species, here represented by the red curve), but not with *A. septentrionalis*, as it is more recent than the divergence between *A. vitaliana* and *A. septentrionalis* (orange curve). *Androsace septentrionalis* only shows signatures of an ancient WGD, likely corresponding to the *Pv-α* WGD event previously identified in a study on *P. veris*[26]. **b.** Bar plots indicating the number of syntenic genes present in each species comparison within *Androsace*. Each box indicates how many homologous syntenic genes in the genome of the species on the x-axis can be found in the genome of the species on the y-axis. Most genes of *A. septentrionalis* have two paralogs in both *A. vitaliana* and *A. wulfeniana*, indicating that a WGD occurred after the divergence between *A. septentrionalis* and the common ancestor of *A. vitaliana* and *A. wulfeniana*.

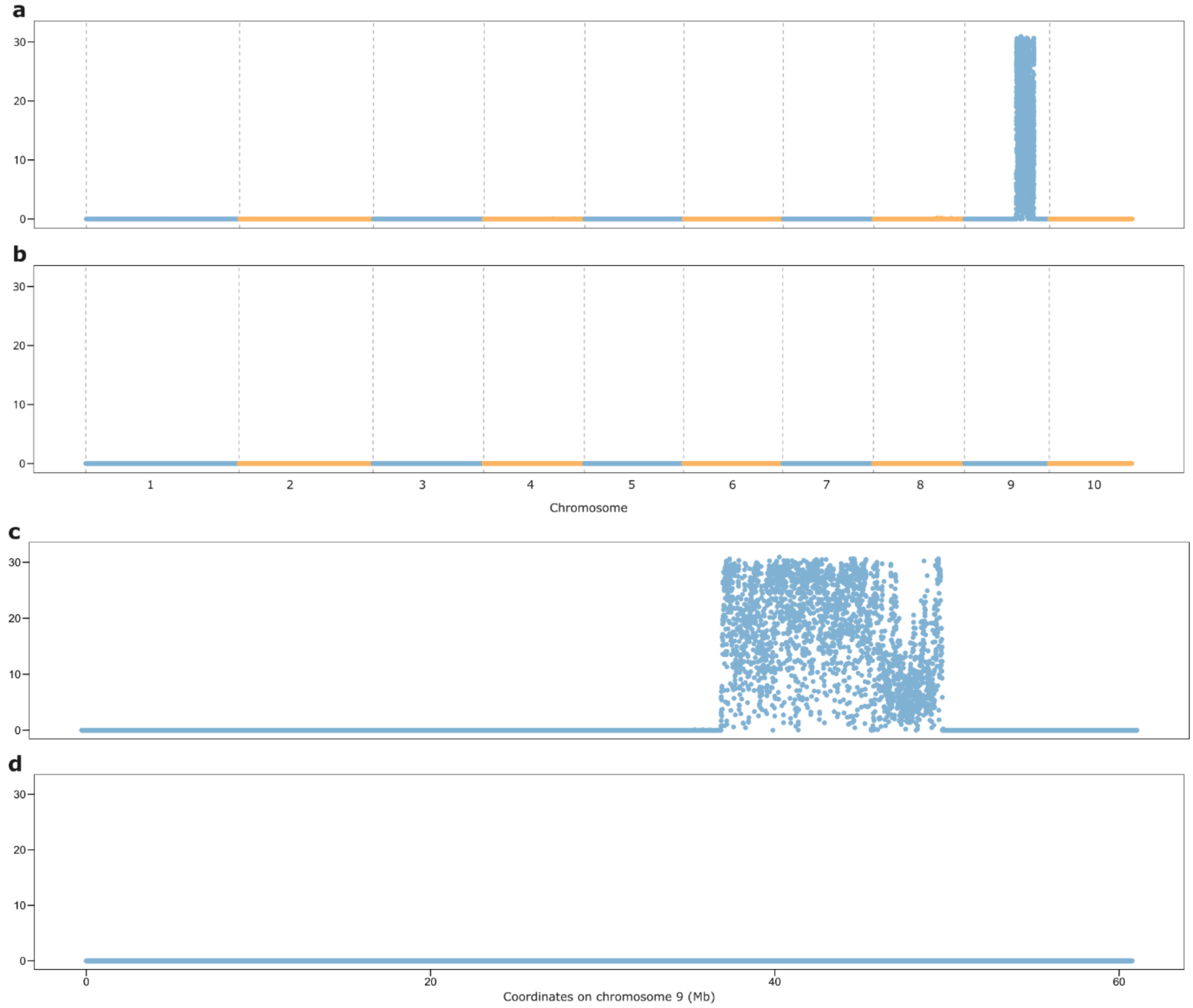

**Fig. S7: Distribution of morph-specific *k*-mers in *Hottonia palustris.***
Morph-specific *k*-mers were identified from 28 samples of a population collected in the Burgwies pond in Zurich. **a-b.** Percentage of sequence covered by thrum-specific (**a**) and pin-specific (**b**) *k*-mers (calculated in 5-kb windows) in the *H. plaustris* assembly (thrum). **c-d.** Zoom-in on chromosome 9, showing the distribution of thrum-specific (**c**) and pin-specific (**d**) *k*-mers. A region on chromosome 9 (36.76-49.53 Mb) is enriched in thrum-specific *k*-mers (**a** and **c**). Conversely, there is no region enriched in pin-specific *k*-mers (**b** and **d**).

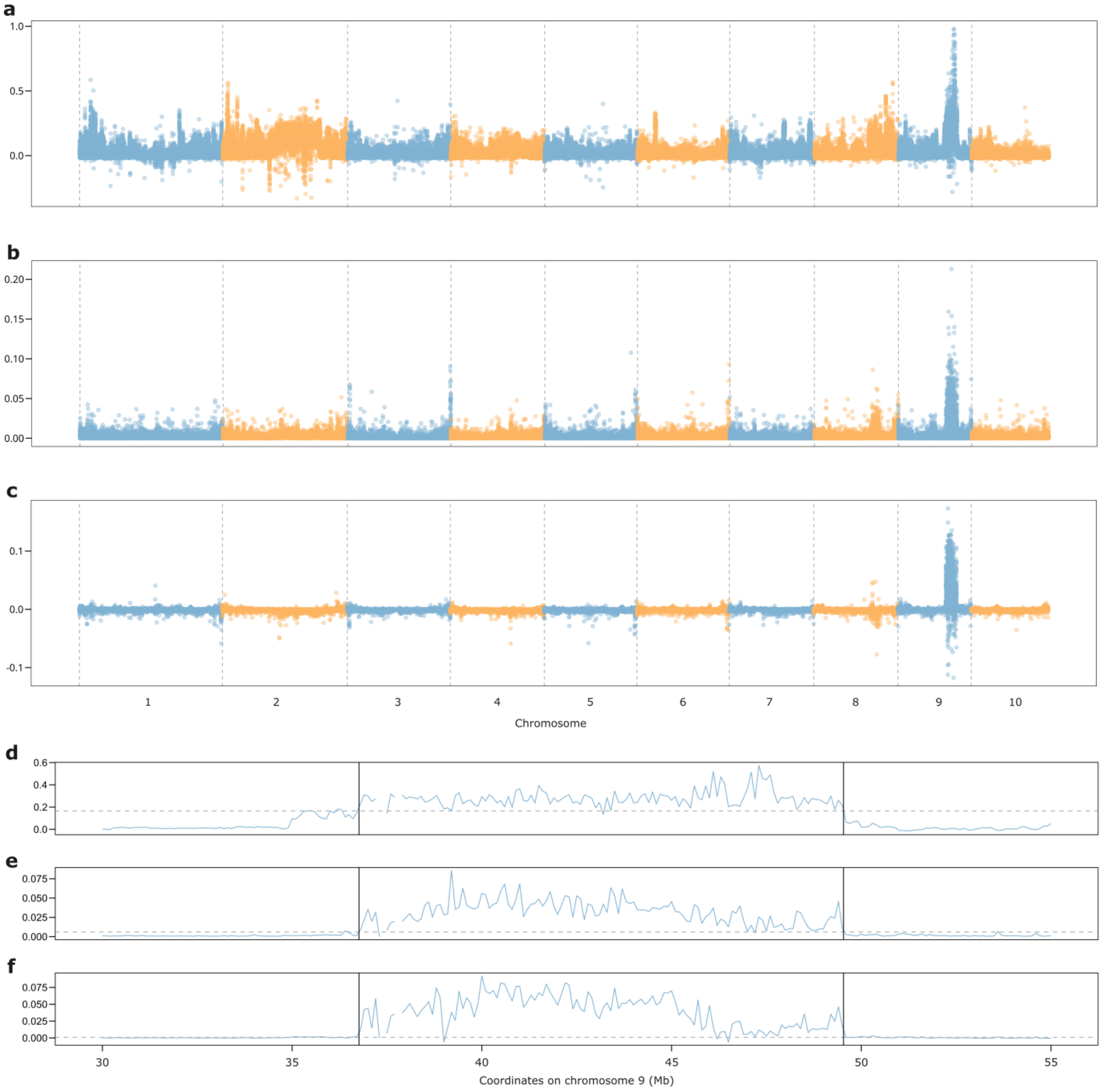

**Fig. S8: Population genetic evidence for the localization of the *S*-locus in *Hottonia palustris*.**
**a-c.** $F_{ST}$ (**a**), $D_{XY}$ (**b**), and morph-biased heterozygosity (**c**; defined as heterozygosity in thrums minus heterozygosity in pins), calculated in 5-kb windows across the *H. palustris* genome (thrum haplotype) using a population of 14 pins and 14 thrums. A peak is visible on chromosome 9 for each distribution. **d-f.** Zoom-in on the region of chromosome 9 (30-55 Mb), characterized by elevated $F_{ST}$ (**d**), $D_{XY}$ (**e**), and morph-biased heterozygosity (**f**); black vertical lines delimit the *S*-locus; a horizontal dashed line corresponds to the 95th percentile of each distribution, calculated on the whole genome.

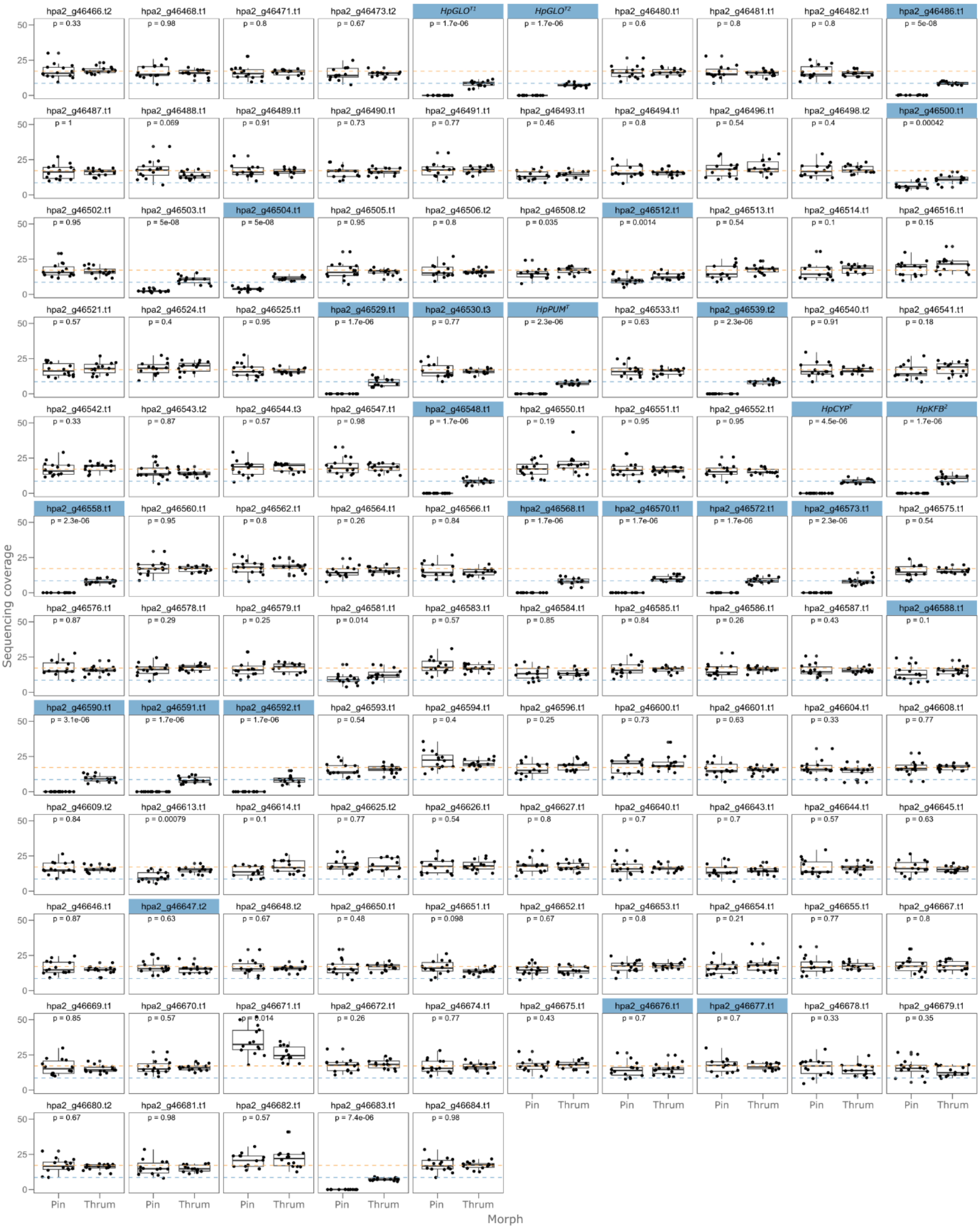

**Fig. S9: Sequencing coverage of *Hottonia palustris S*-haplotype genes.**
Box plots showing sequencing coverage calculated for 14 pins and 14 thrums on the 115 *S*-haplotype genes of *H. palustris*. The mean coverage calculated for all genes on chromosome 9 is represented by an orange dashed horizontal line; a blue dashed horizontal line marks the coverage expected for a hemizygous gene (i.e. half the mean coverage); p-values represent pairwise comparisons between groups, conducted using the Wilcoxon rank-sum test. Genes specific to the *S*-haplotype are highlighted in blue; these include the orthologs of *Primula S*-genes: *HpGLO*$^{T1}$ (*hpa2_g46474.t1*), *HpGLO*$^{T2}$ (*hpa2_g46478.t1*), *HpPUM*$^{T}$ (*hpa2_g46531.t1*), *HpCYP*$^{T}$ (*hpa2_g46555.t1*), and *HpKFB*$^{T}$ (*hpa2_g46556.t1*). Genes specific to the *S*-haplotype that show a similar coverage in the two morphs likely represent genes that pseudogenized in in the *S*-haplotype.

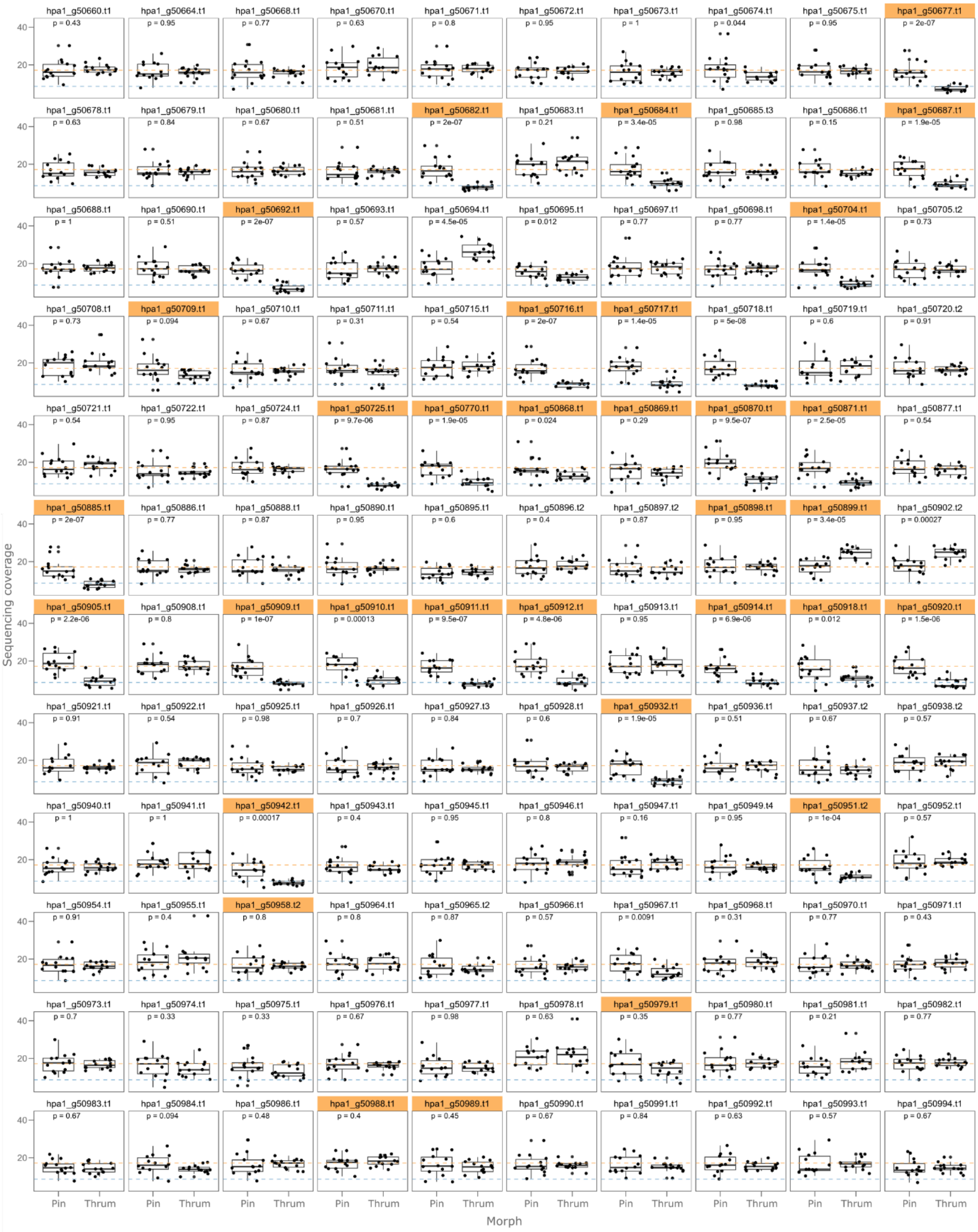

**Fig. S10: Sequencing coverage of *Hottonia palustris s*-haplotype genes.**
Box plots showing sequencing coverage calculated for 14 pins and 14 thrums on the 120 *s*-haplotype genes of *H. palustris*. The mean coverage calculated for all genes on chromosome 9 is represented by an orange dashed horizontal line; a blue dashed horizontal line marks the coverage expected for a hemizygous gene (i.e. half the mean coverage); p-values represent pairwise comparisons between groups, conducted using the Wilcoxon rank-sum test. Genes specific to the *s*-haplotype are highlighted in orange. Genes specific to the *s*-haplotype that show a similar coverage in the two morphs likely represent genes that pseudogenized in in the *s*-haplotype.

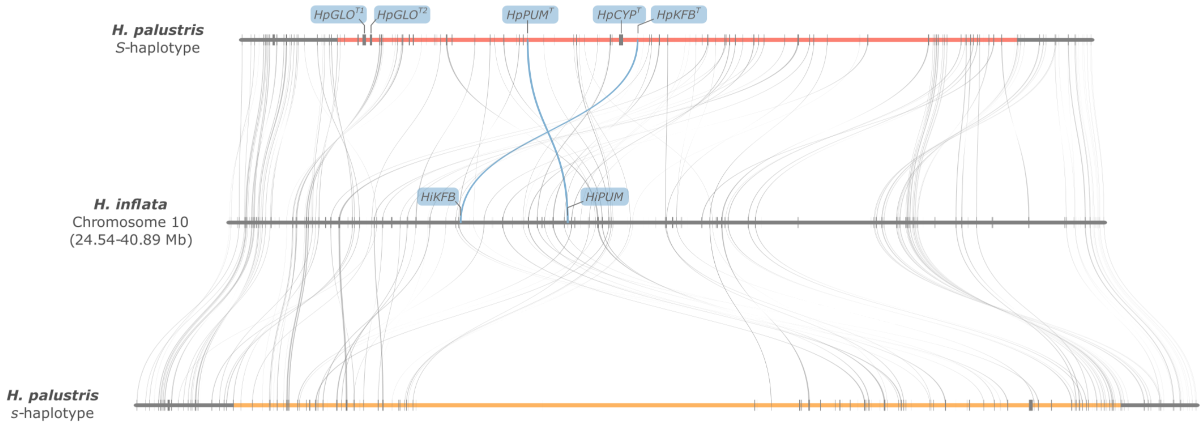

**Fig. S11: The *Hottonia palustris S*-locus shows suppressed recombination and is rearranged compared to *H. inflata.***

Microsynteny plot between the *H. palustris S*- and *s*-haplotypes (top and bottom, respectively), and their syntenic region in *H. inflata* (center). The *H. palustris S*- and s-haplotypes are highlighted in red and orange, respectively; gene names are reported for orthologs of *Primula S*-genes. *Hottonia inflata* contains a degenerated *S*-locus lacking orthologs of *GLO*$^{T}$ and *CYP*$^{T}$, indicating that this species represents a secondary loss of distyly. Furthermore, this plot shows that several rearrangements occurred in this genomic region since the divergence between *H. palustris* and *H. inflata*.

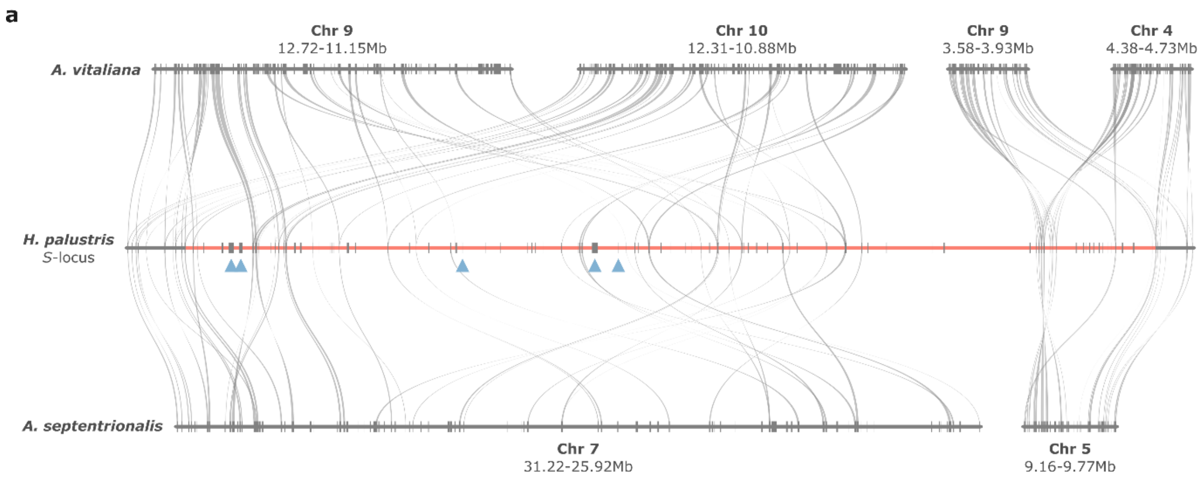

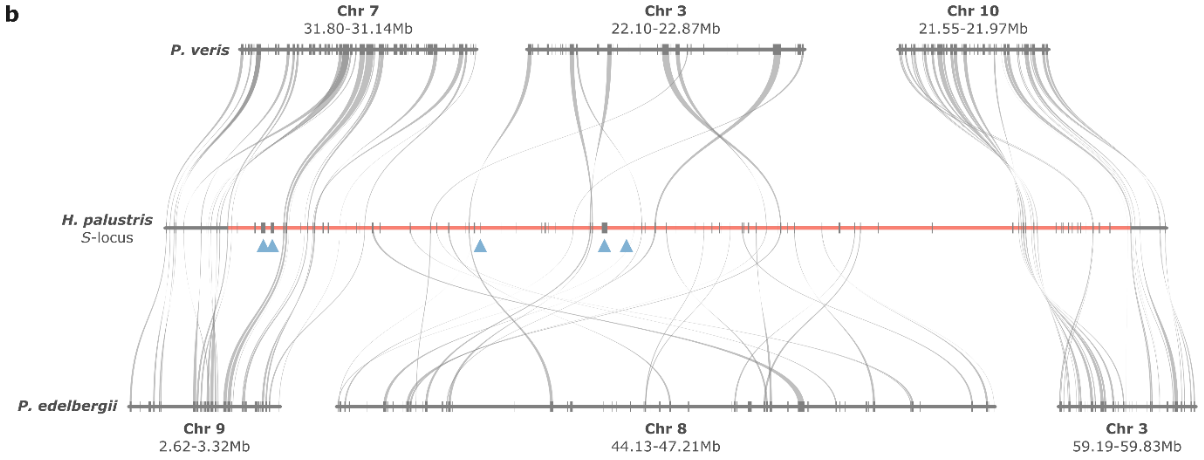

**Fig. S12: The *Hottonia palustris S*-locus lies in a region not syntenic with *Androsace* nor *Primula*.**
**a.** Microsynteny plot between the *H. palustris S*-locus (*S*-haplotype; center), *A. vitaliana* (thrum haplotype; top) and *A. septentrional*is (bottom). **b.** Microsynteny plot between the *H. palustris S*-locus (*S*-haplotype; center), *P. veris* (top) and *P. edelbergii* (bottom). The region containing the *S*-locus in *H. palustris* (chromosome 9: 36.00-50.07 Mb) has the *S*-locus colored red and the five hemizygous orthologs of the *Primula S*-genes marked by blue triangles (left to right: $HpGLO^{T1}$, $HpGLO^{T2}$, $HpPUM^{T}$, $HpCYP^{T}$, and $HpKFB^{T}$); none of these five genes is contained in a region syntenic with any other Primulaceae. Instead, parts of the *H. palustris S*-locus are syntenic with multiple regions in different chromosomes in each comparison, indicating that the *S*-locus: i) translocated after the divergence between *Hottonia* and *P. veris*/*P. edelbergii* and ii) underwent several structural rearrangements in *H. palustris*, incorporating additional genes.

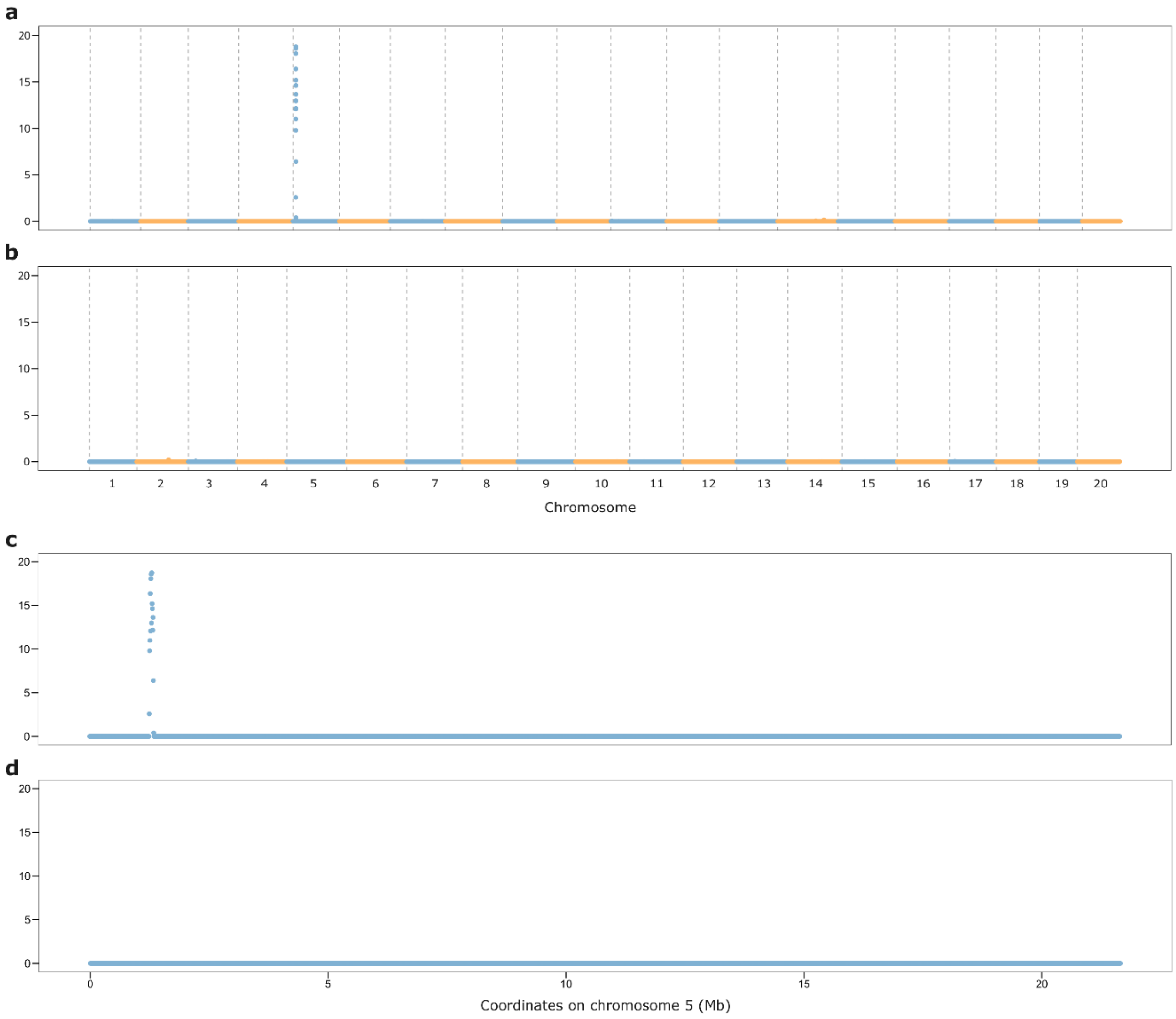

**Fig. S13: Distribution of morph-specific *k*-mers in *Androsace vitaliana* (Wallis samples).**
Morph-specific *k*-mers were identified from 24 samples (12 pins and 12 thrums) of a population collected in the Wallis canton (Switzerland). **a-b.** Percentage of sequence covered by thrum-specific (**a**) and pin-specific (**b**) *k*-mers (calculated in 5-kb windows) in the *A. vitaliana* assembly (thrum). **c-d.** Zoom-in on chromosome 5, showing the distribution of thrum-specific (**c**) and pin-specific (**d**) *k*-mers. A region on chromosome 5 (950-1,100 kb) is enriched in thrum-specific *k*-mers (**a** and **c**). Conversely, there is no region enriched in pin-specific *k*-mers (**b** and **d**).

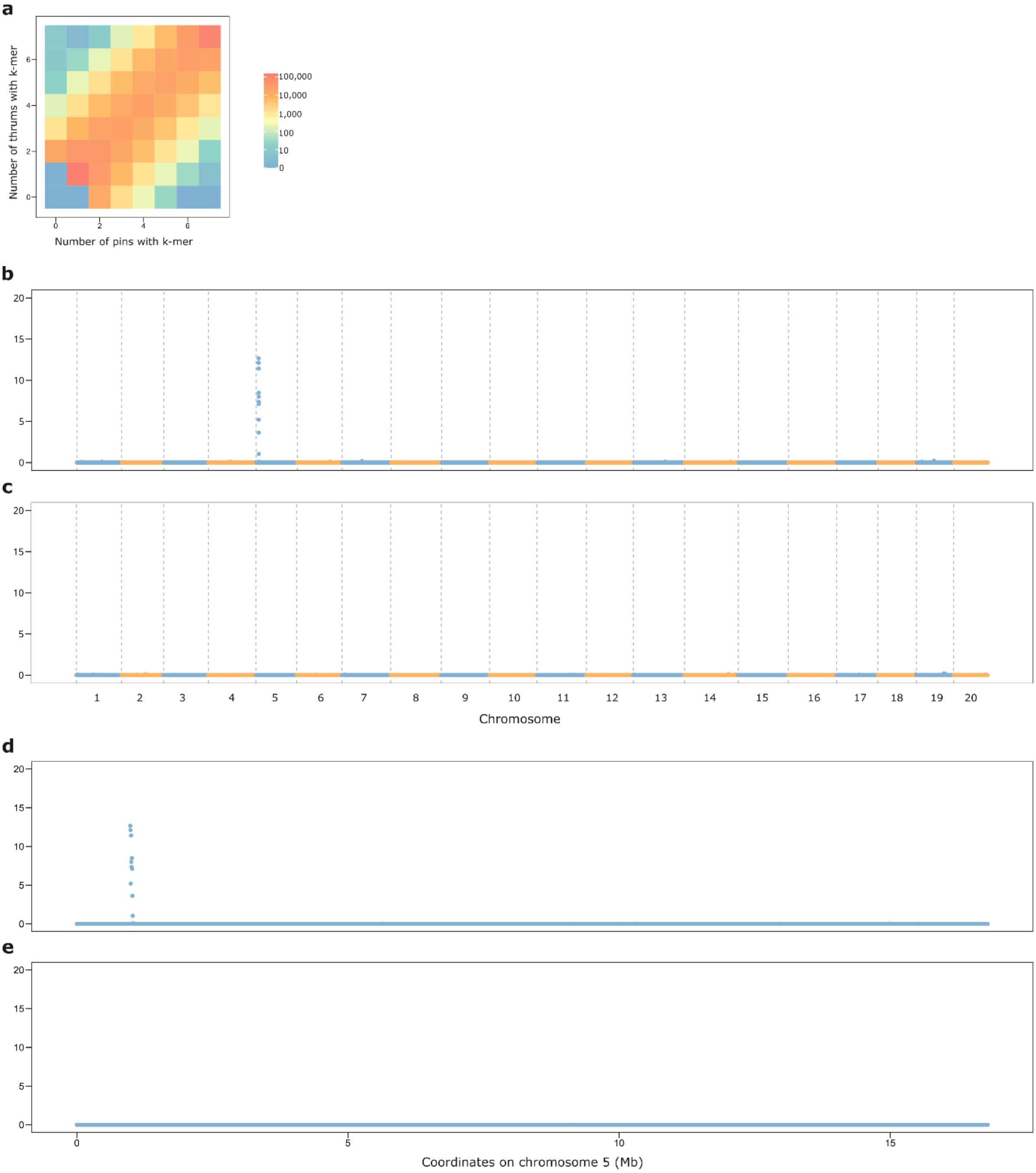

**Fig. S14: Distribution of morph-specific *k*-mers in *Androsace vitaliana* (herbarium samples).** Morph-specific *k*-mers were identified from 14 samples (7 pins and 7 thrums) whose DNA was extracted from herbarium specimens. a-b. Percentage of sequence covered by thrum-specific (**a**) and pin-specific (**b**) *k*-mers (calculated in 5-kb windows) in the *A. vitaliana* assembly (thrum). **c-d.** Zoom-in on chromosome 5, showing the distribution of thrum-specific (**c**) and pin-specific (**d**) *k*-mers. A region on chromosome 5 (950-1,100 kb) is enriched in thrum-specific *k*-mers (**a** and **c**). Conversely, there is no region enriched in pin-specific *k*-mers (**b** and **d**).

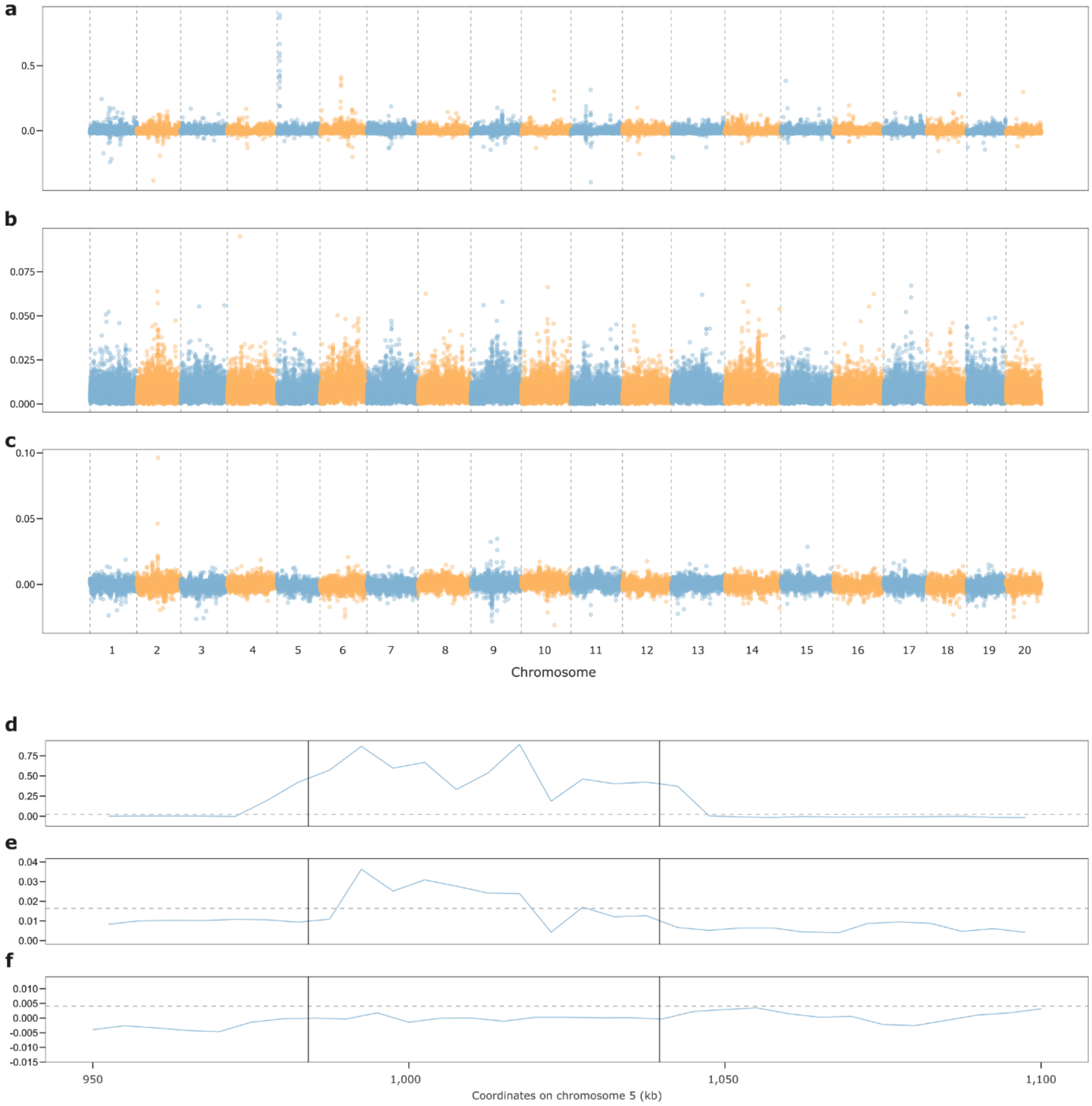

**Fig. S15: Population genetic evidence for the localization of the *S*-locus in *Androsace vitaliana* (Wallis samples).**
**a-c.** $F_{ST}$ (**a**), $D_{XY}$ (**b**), and morph-biased heterozygosity (**c**; defined as heterozygosity in thrums minus heterozygosity in pins), calculated in 5-kb windows across the *A. vitaliana* genome (thrum haplotype) using 24 samples (12 pins and 12 thrums) of a population collected in the Wallis canton (Switzerland). A peak is visible on chromosome 5 for the $F_{ST}$ distribution. **d-f.** Zoom-in on the region of chromosome 5 (950-1,100 kb), characterized by elevated $F_{ST}$ (**d**). Black vertical lines delimit the *S*-locus; a horizontal dashed line corresponds to the 95th percentile of each distribution, calculated on the whole genome. Values above the 95th percentile were detected in this region for $F_{ST}$ (**d**) and $D_{XY}$ (**e**), but not for the morph-biased heterozygosity (**f**). This is likely due to the small size of the *S*-locus in A. vitaliana (ca. 55 kb) and to its high TE content. Indeed, higher heterozygosity in thrums was observed when looking at single SNPs (Fig. 3 in the main text).

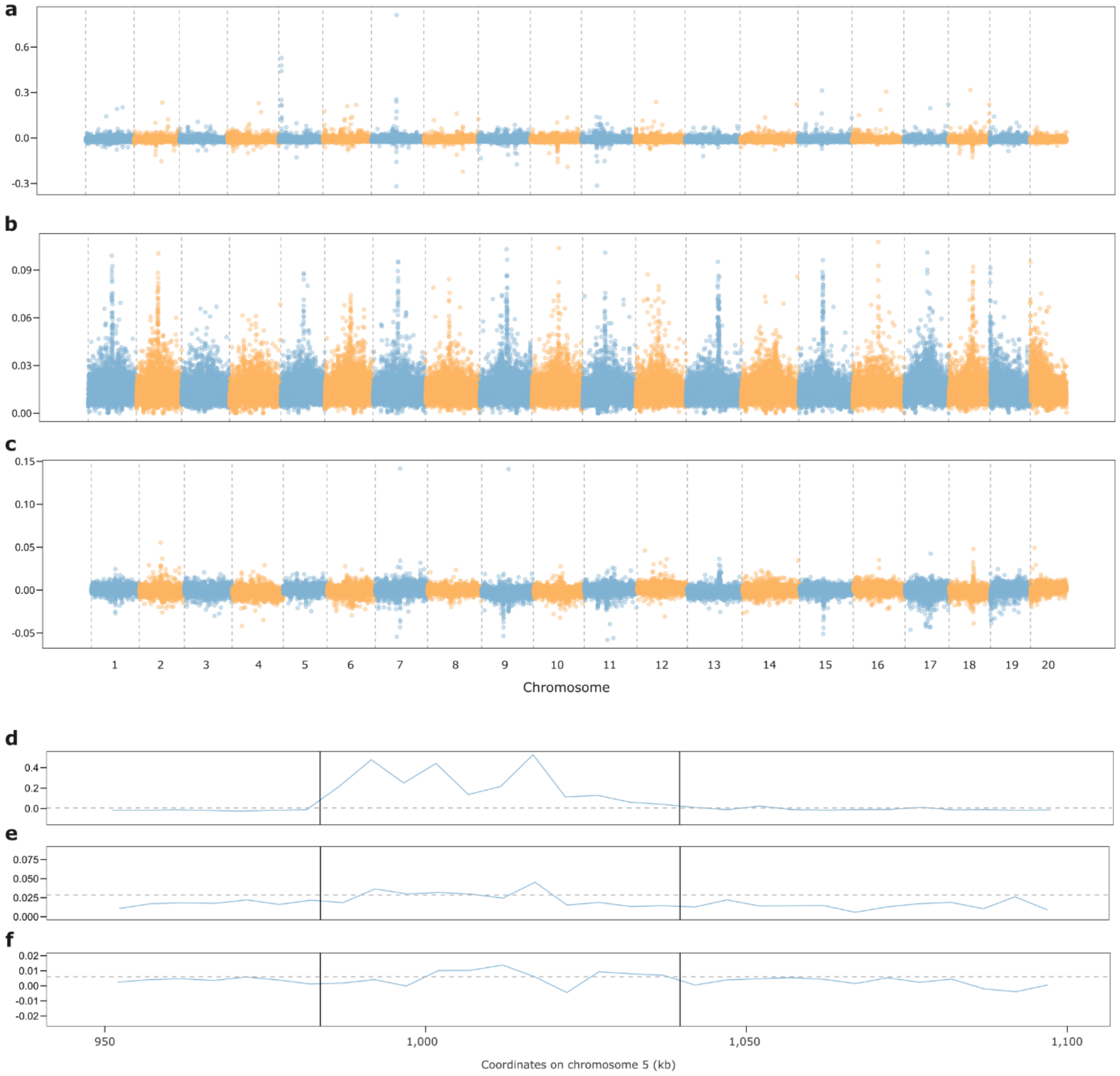

**Fig. S16: Population genetic evidence for the localization of the *S*-locus in *A. vitaliana* (herbarium samples).**
**a-c.** $F_{ST}$ (**a**), $D_{XY}$ (**b**), and morph-biased heterozygosity (**c**; defined as heterozygosity in thrums minus heterozygosity in pins), calculated in 5-kb windows across the *A. vitaliana* genome (thrum haplotype) using 14 samples (7 pins and 7 thrums) whose DNA was extracted from herbarium specimens. A peak is visible on chromosome 5 for the $F_{ST}$ distribution. **d-f.** Zoom-in on the region of chromosome 5 (950-1,100 kb), characterized by elevated $F_{ST}$ (**d**). Black vertical lines delimit the *S*-locus; a horizontal dashed line corresponds to the 95th percentile of each distribution, calculated on the whole genome. Values above the 95th percentile were detected in this region for $F_{ST}$ (**d**) and $D_{XY}$ (**e**), but not for the morph-biased heterozygosity (**f**). This is likely due to the small size of the *S*-locus in A. vitaliana (ca. 55 kb) and to its high TE content. Indeed, higher heterozygosity in thrums was observed when looking at single SNPs (Fig. 3 in the main text).

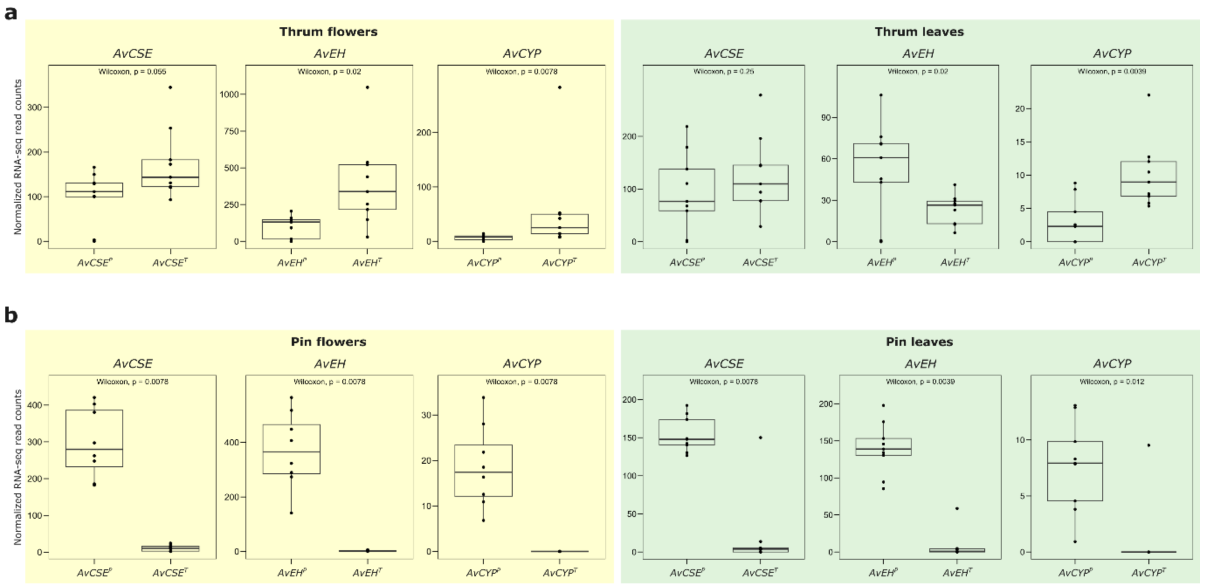

**Fig. S17: *Androsace vitaliana S*-alleles are upregulated in flowers compared to *s*-alleles.**
Box plots comparing the expression of *s*- (left) and *S*- (right) alleles in flowers (yellow box) and leaves (green box). **a.** Results from nine thrum flower samples and nine thrum leaf samples. In flowers, $AvEH^T$ and $AvCYP^T$ were upregulated compared to $AvEH^P$ and $AvCYP^P$, while *AvCSE* alleles were expressed at the same level; in leaves, $AvEH^P$ was upregulated compared to $AvEH^T$, *AvCSE* alleles were expressed at the same level, and both *AvCYP* alleles were expressed at extremely low levels. **b.** Results from eight pin flower samples and nine pin samples. In pins, no expression was detected for *S*-allelels, as expected since pins only carry the *s*-haplotype; this result proves that the difference in expression detected in thrums is not due to a bias in RNA-seq read mapping but rather represents a true biological difference.

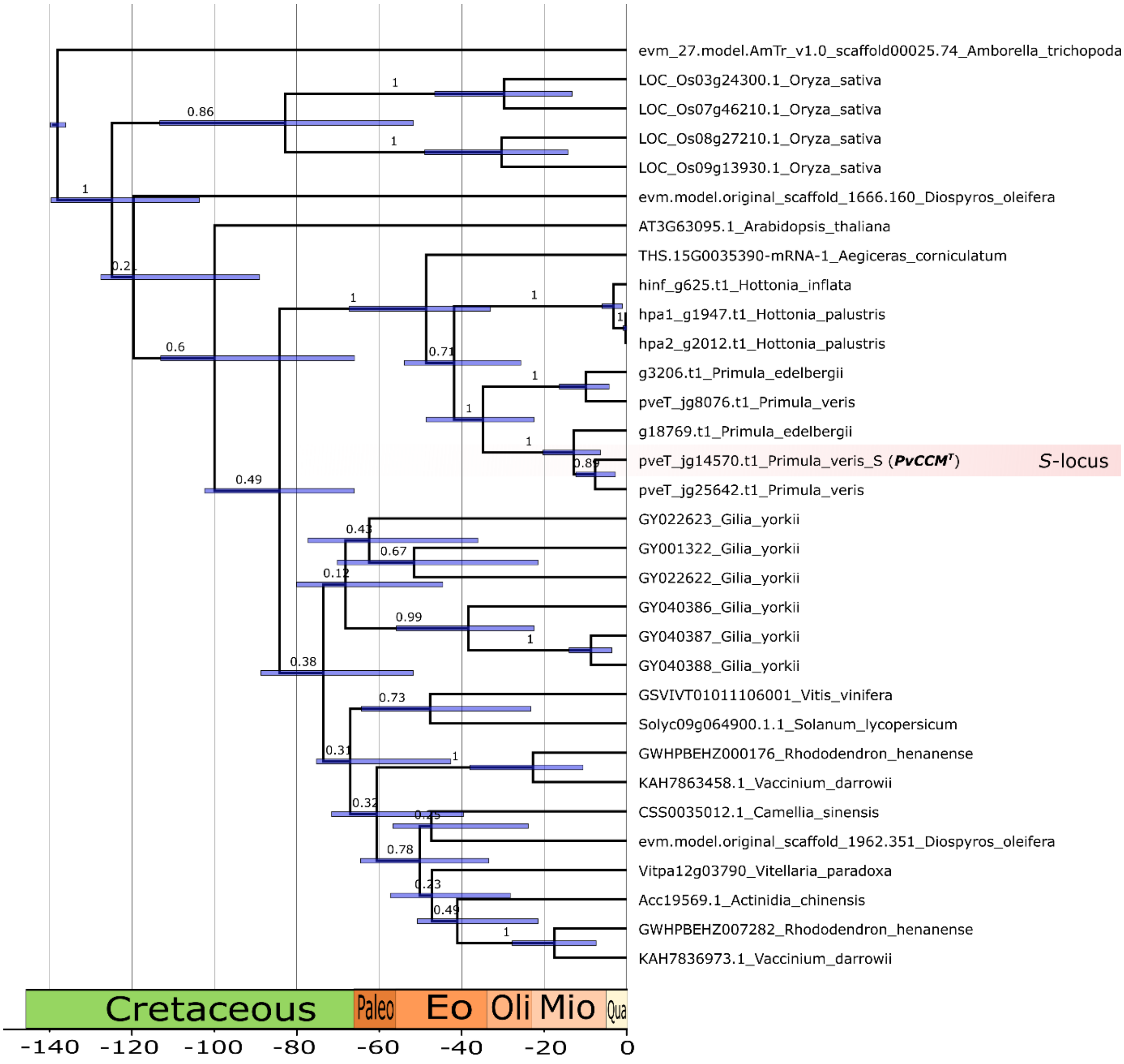

**Fig. S18: Phylogeny of the *S*-gene *CCM$^T$* and close homologs.**
Bayesian chronogram of *CCM$^T$* genes in selected select genomes. Bottom scale bar indicates time before present in million years (My), with boxes indicating geological periods (pre-Cenozoic) or epochs (Cenozoic). Blue bars at nodes represent 95% Bayesian credibility intervals around age estimates. Branch labels represent posterior probabilities for the subtended clade.

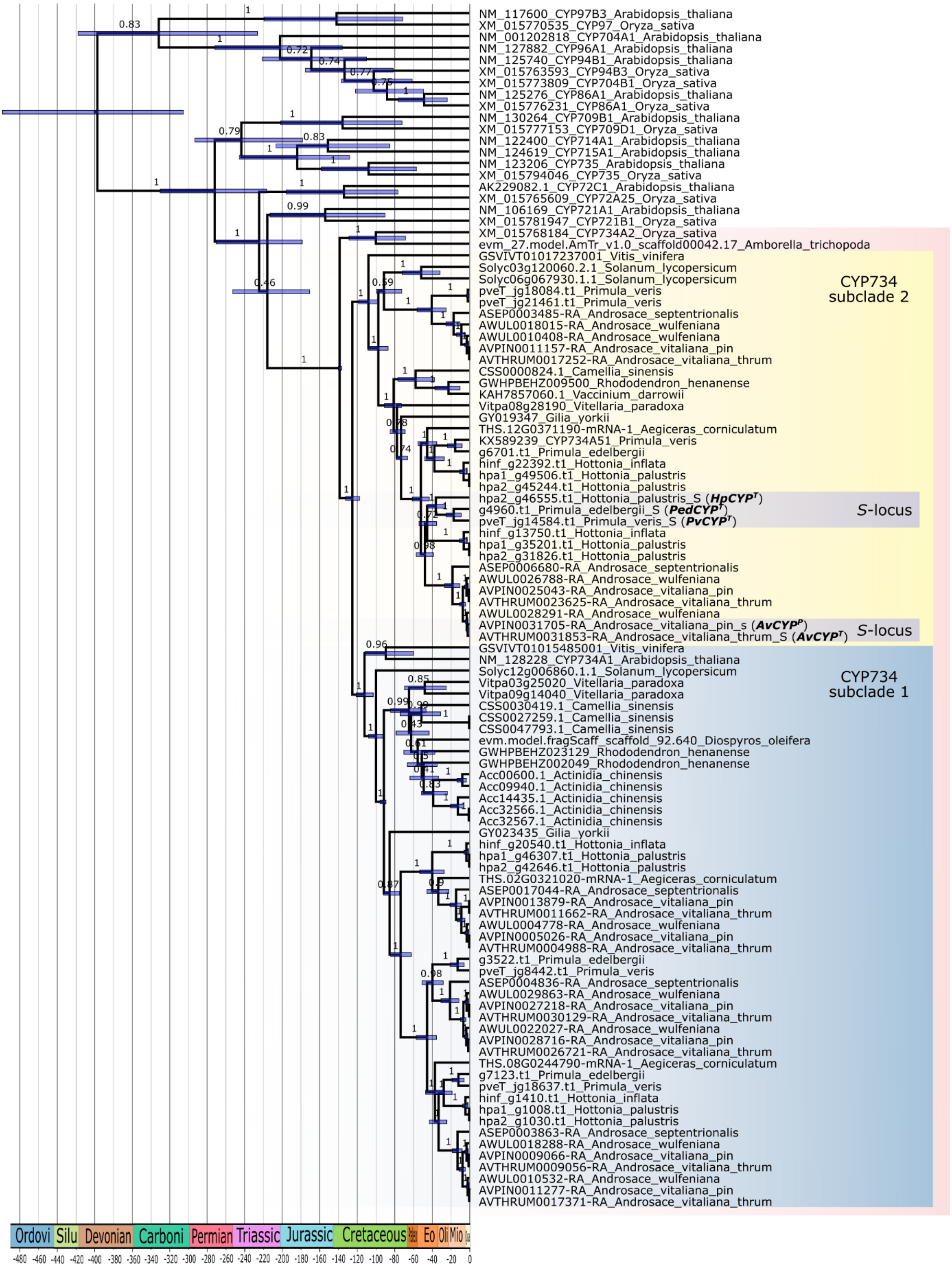

**Fig. S19: Phylogeny of the *S*-gene *CYP*$^{T}$ and close homologs.**
Bayesian chronogram of *CYP*$^{T}$ genes in selected select genomes. Bottom scale bar indicates time before present in million years (My), with boxes indicating geological periods (pre-Cenozoic) or epochs (Cenozoic). Blue bars at nodes represent 95% Bayesian credibility intervals around age estimates. Branch labels represent posterior probabilities for the subtended clade.

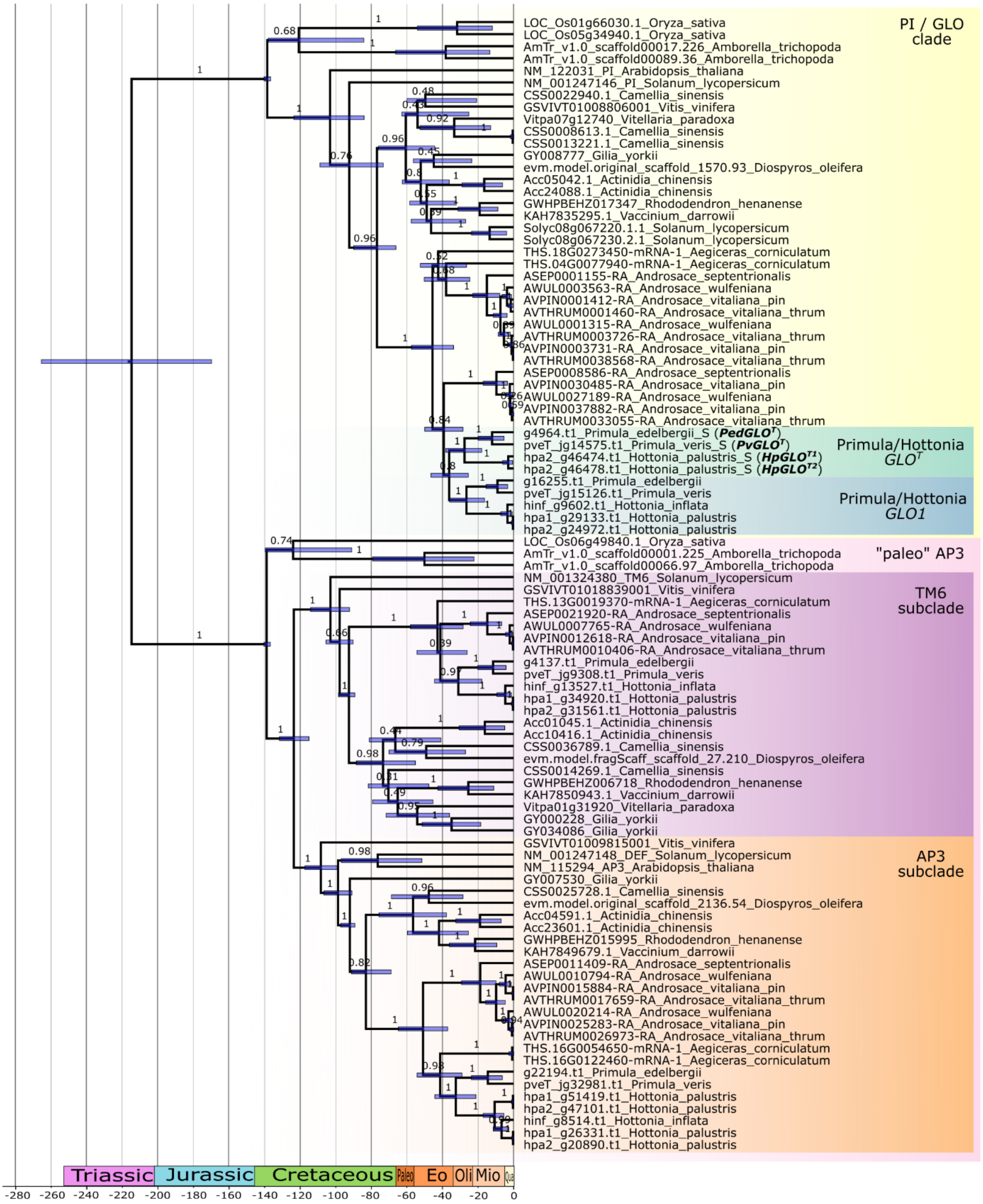

**Fig. S20: Phylogeny of the *S*-gene *GLO*$^T$ and close homologs.**
Bayesian chronogram of *GLO*$^T$ genes in selected select genomes. Bottom scale bar indicates time before present in million years (My), with boxes indicating geological periods (pre-Cenozoic) or epochs (Cenozoic). Blue bars at nodes represent 95% Bayesian credibility intervals around age estimates. Branch labels represent posterior probabilities for the subtended clade.

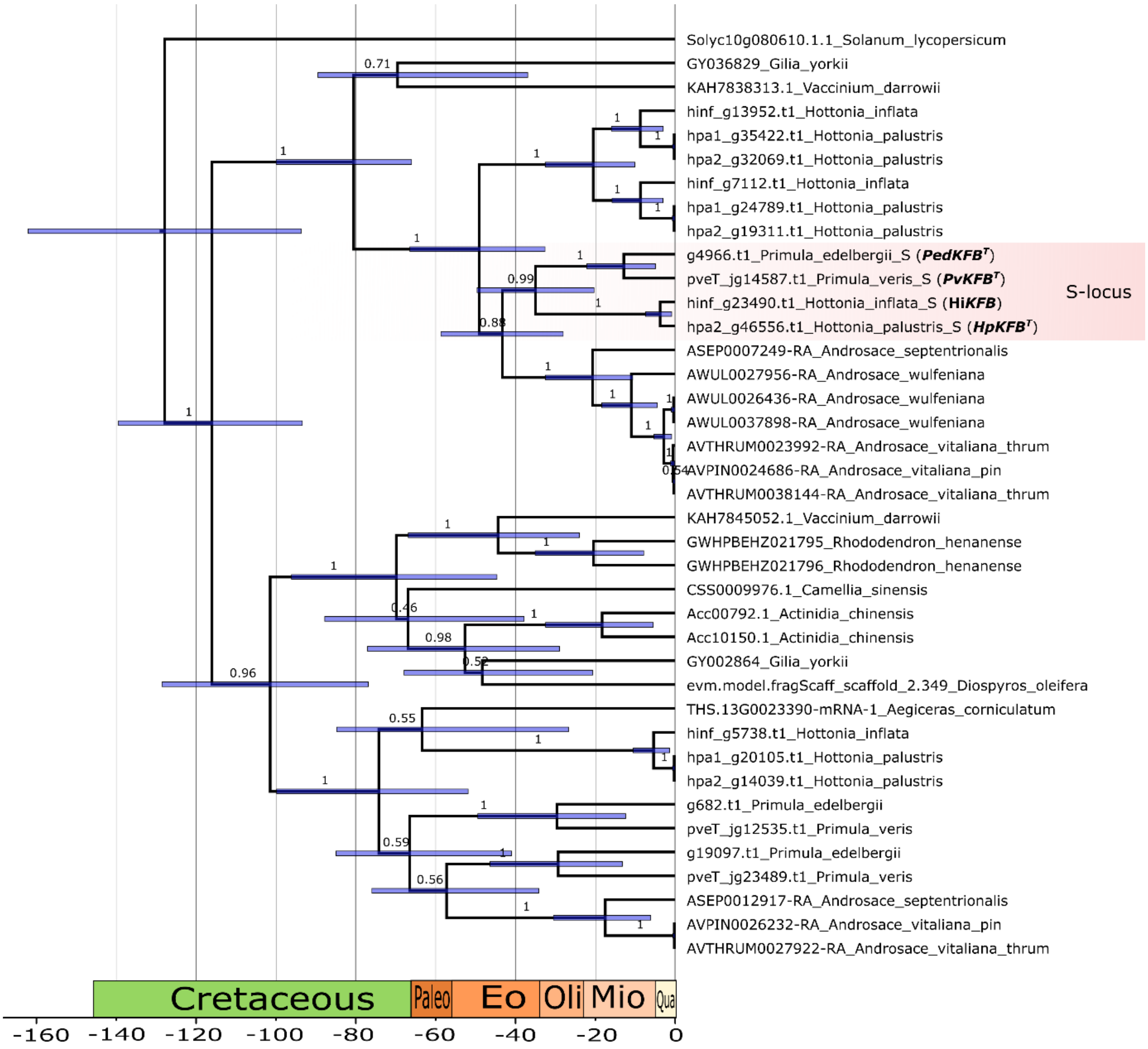

**Fig. S21: Phylogeny of the *S*-gene *KFB*$^T$ and close homologs.**
Bayesian chronogram of *KFB*$^T$ genes in selected select genomes. Bottom scale bar indicates time before present in million years (My), with boxes indicating geological periods (pre-Cenozoic) or epochs (Cenozoic). Blue bars at nodes represent 95% Bayesian credibility intervals around age estimates. Branch labels represent posterior probabilities for the subtended clade.

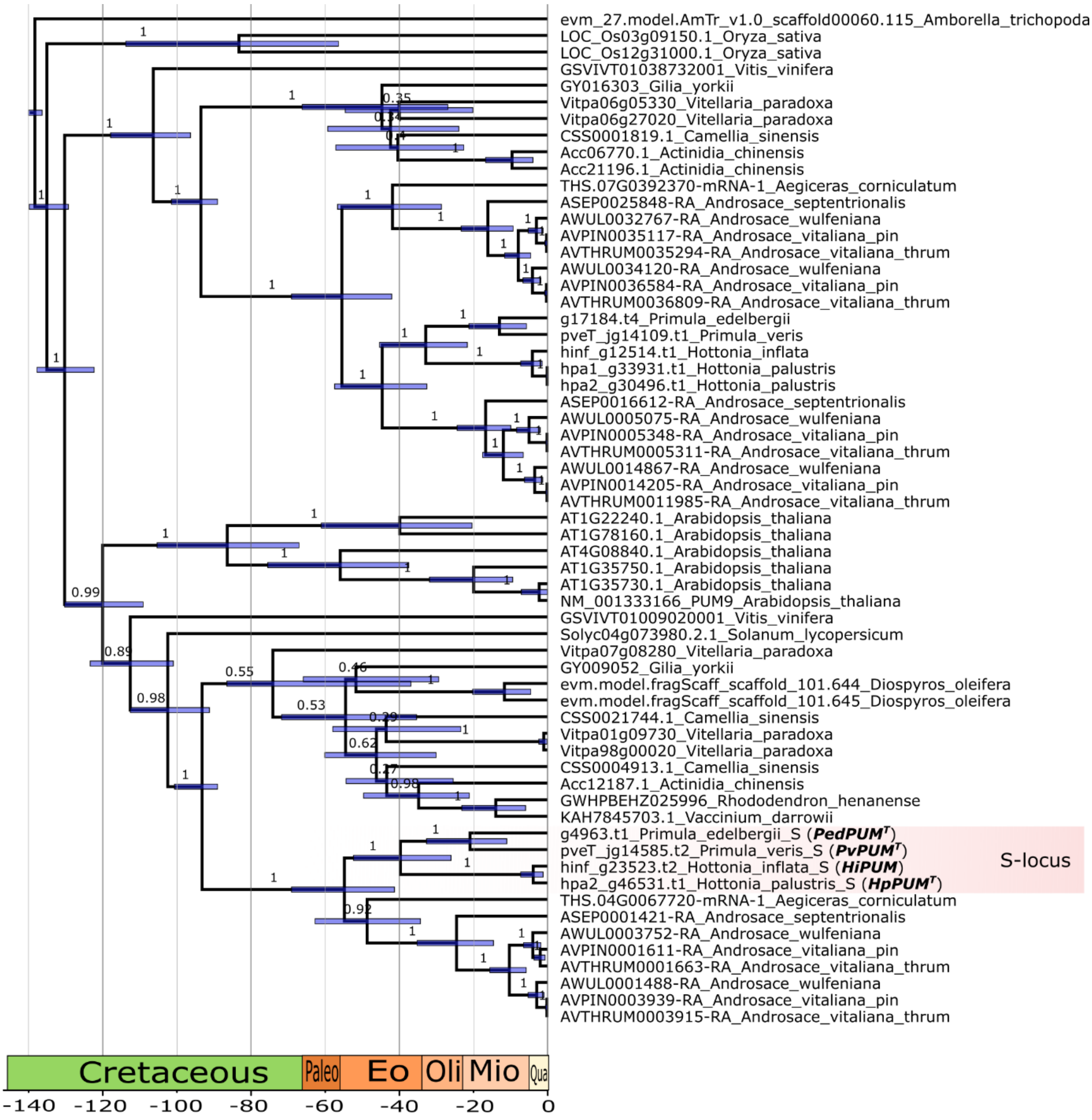

**Fig. S22: Phylogeny of the *S*-gene $PUM^T$ and close homologs.**
Bayesian chronogram of $PUM^T$ genes in selected select genomes. Bottom scale bar indicates time before present in million years (My), with boxes indicating geological periods (pre-Cenozoic) or epochs (Cenozoic). Blue bars at nodes represent 95% Bayesian credibility intervals around age estimates. Branch labels represent posterior probabilities for the subtended clade.

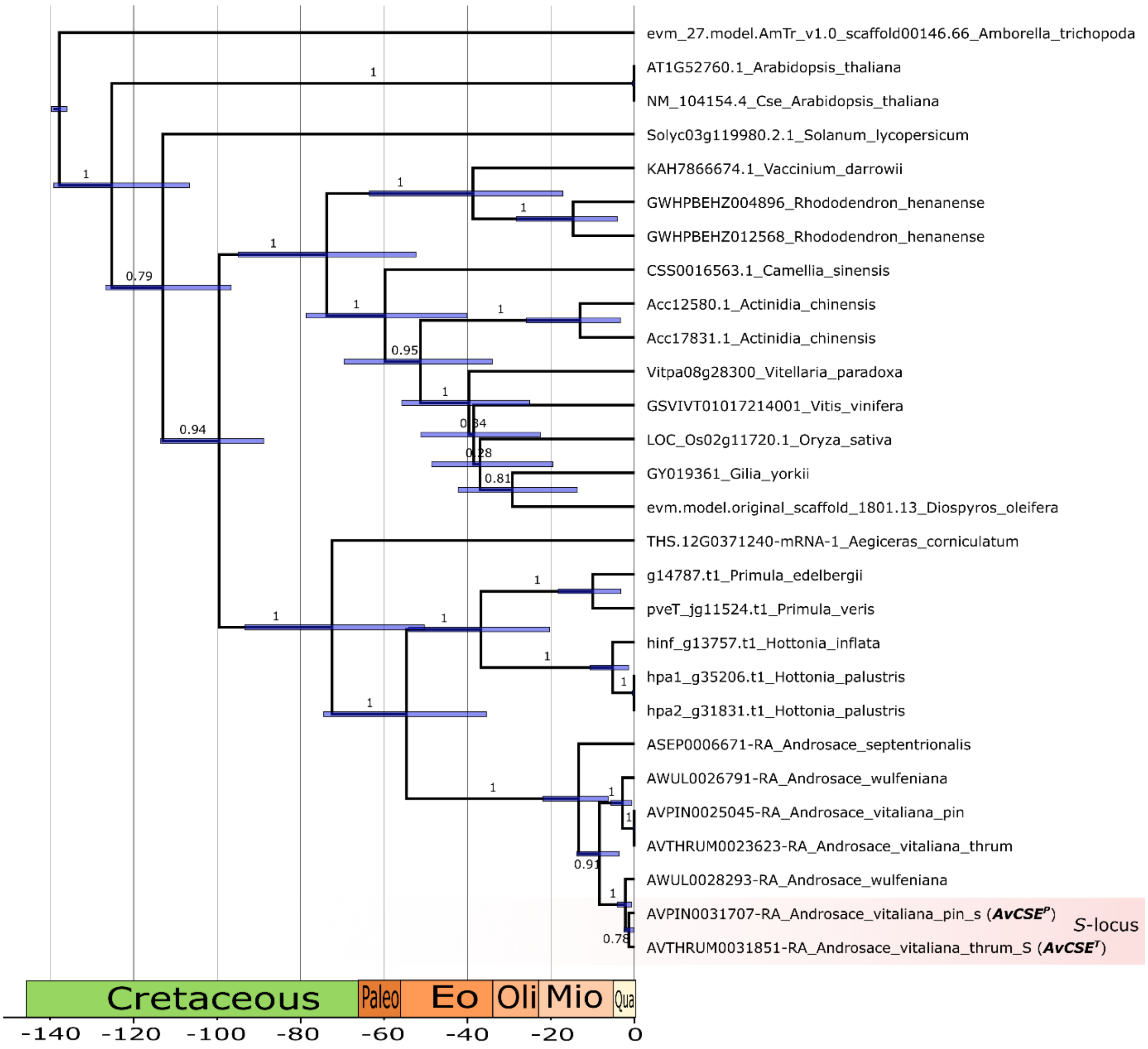

**Fig. S23: Phylogeny of the *S*-gene $AvCSE^T$ and close homologs.**

Bayesian chronogram of $AvCSE^T$ genes in selected select genomes. Bottom scale bar indicates time before present in million years (My), with boxes indicating geological periods (pre-Cenozoic) or epochs (Cenozoic). Blue bars at nodes represented 95% Bayesian credibility intervals around age estimates. Branch labels represent posterior probabilities for the subtended clade.

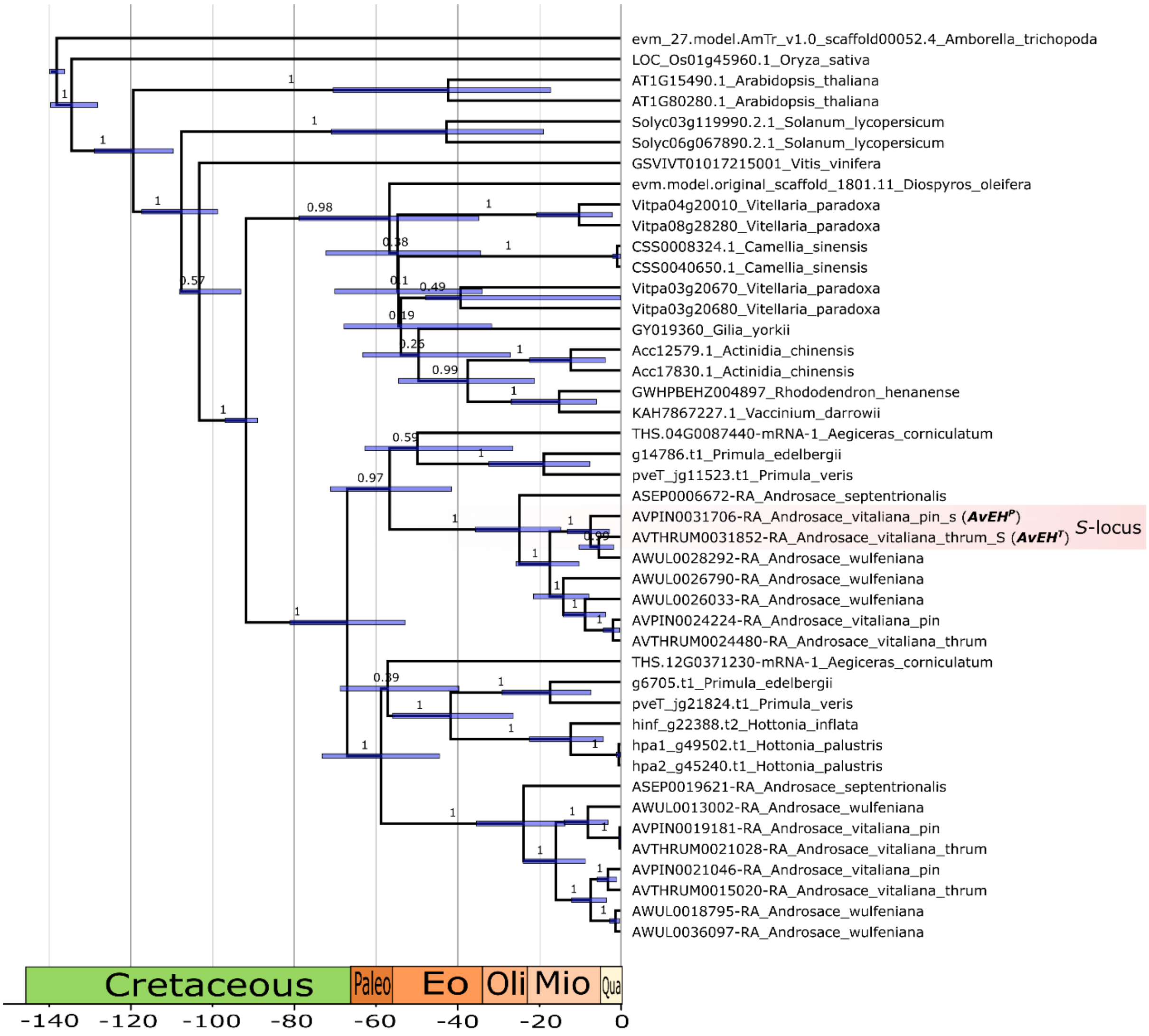

**Fig. S24: Phylogeny of the *S*-gene $AvEH^{T}$ and close homologs.**

Bayesian chronogram of $AvEH^{T}$ genes in selected select genomes. Bottom scale bar indicates time before present in million years (My), with boxes indicating geological periods (pre-Cenozoic) or epochs (Cenozoic). Blue bars at nodes represented 95% Bayesian credibility intervals around age estimates. Branch labels represent posterior probabilities for the subtended clade.

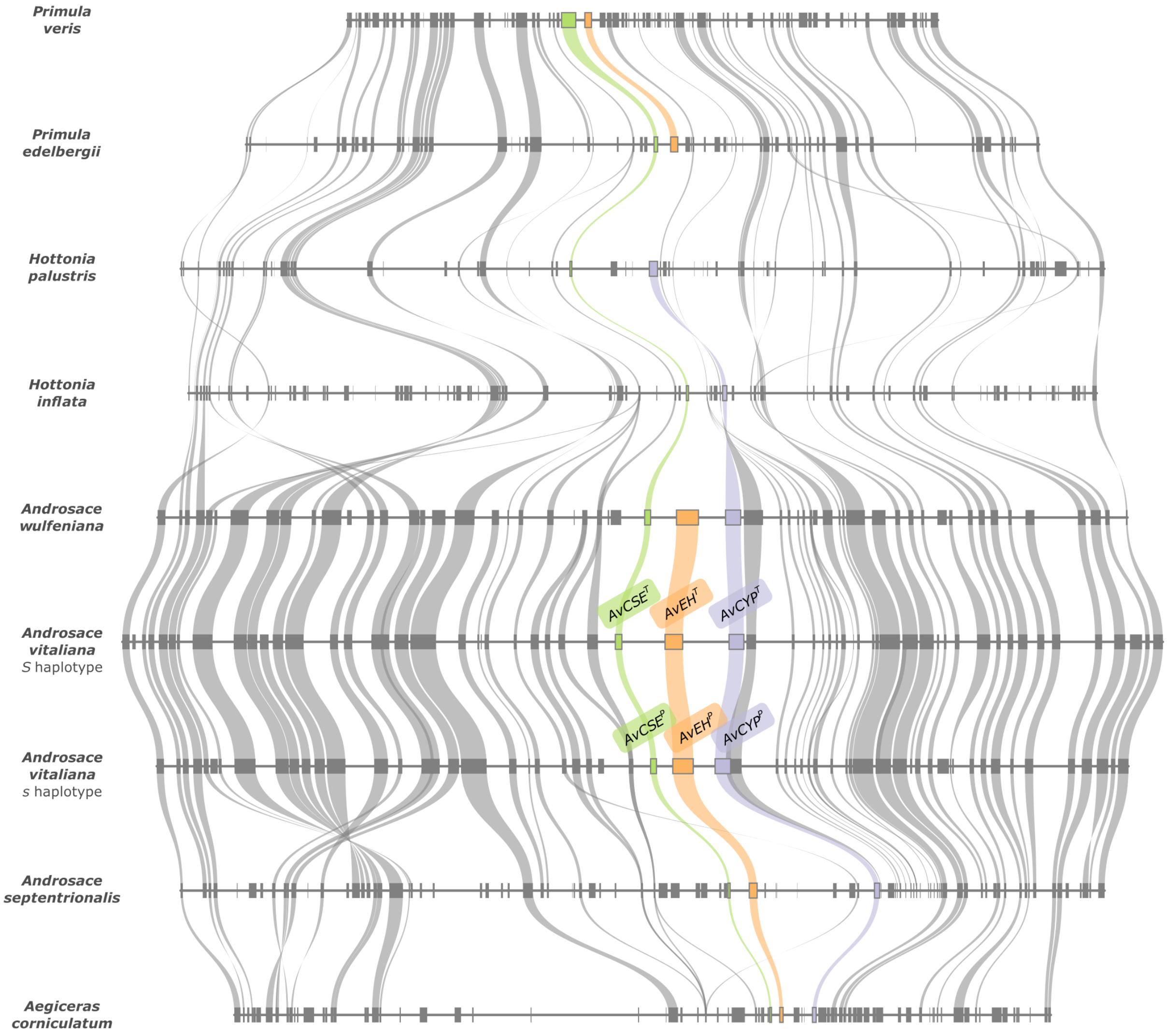

**Fig. S25: The *Androsace vitaliana S*-genes were ancestrally colocalized.**
Microsynteny plot across nine Primulaceae genome assemblies, showing that the *A. vitaliana S*-genes (*AvCSE*, *AvEH*, and *AvCYP*, colored in green, orange, and purple, respectively) are contained in a genomic region syntenic across Primulaceae. The orthologs of the three *A. vitaliana S*-genes are present in this region in all *Androsace* species and in the outgroup *Aegiceras corniculatum*, while *Primula* and *Hottonia* species lack orthologs of *AvCYP* and *AvEH*, respectively.

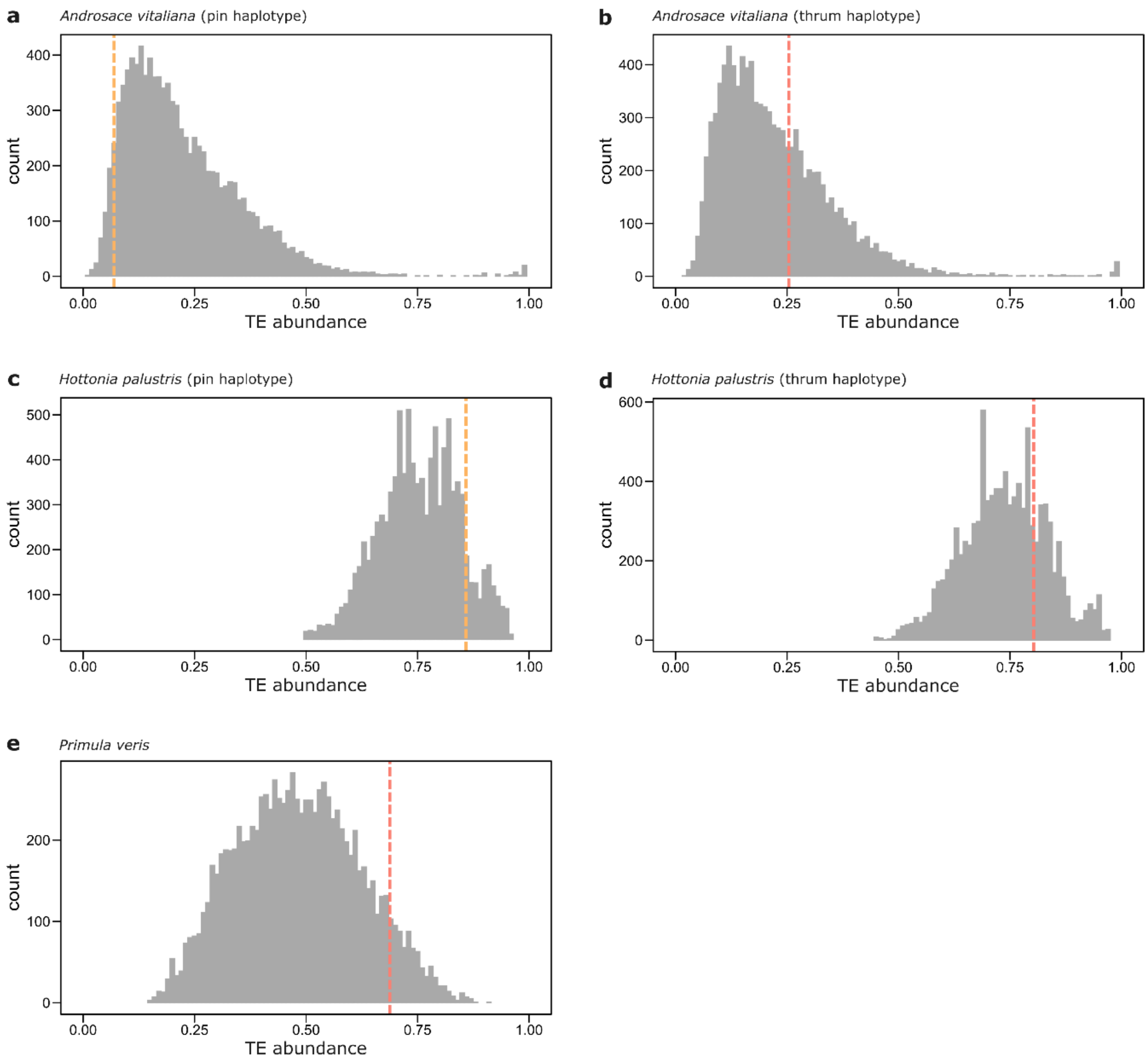

**Fig. S26: Comparison of TE abundance between *S*-loci and genomic background.**
Histograms showing the distribution of TE abundance across 10,000 genomic windows (each matching the size of the *S*-locus) randomly sampled from the genome. Vertical dashed lines indicate TE abundance within the *S*-locus (orange: *s*-haplotype; red: *S*-haplotype). For each species, we assessed whether the *S*-locus was significantly enriched in TEs by calculating an empirical p-value, defined as the proportion of random windows with TE content equal to or greater than that of the S-locus. No significant enrichment of TEs in the *S*-locus was detected.

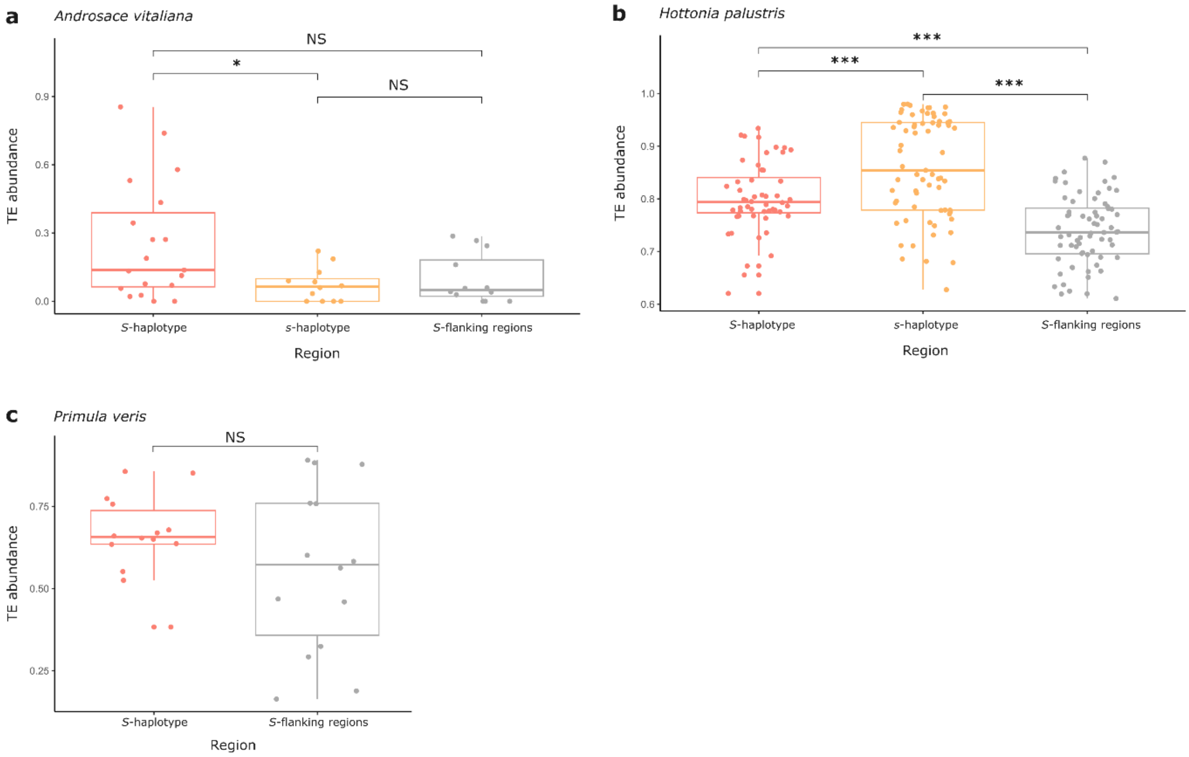

**Fig. S27: Comparison of TE abundance between *S*-loci and their flanking regions.**
Box-plots indicating, for each species, TE abundance calculated in windows within the *S*-locus (*s*- and *S*-haplotypes) and in its flanking regions, defined as regions upstream and downstream the *S*-locus, whose length combined equals the *S*-locus length (*, $0.01 < p < 0.05$; ***, $p < 0.001$; NS, non-significant; Wilcoxon rank-sum test). **a.** *A. vitaliana* (3-kb windows): *S*-haplotype (red, n=19); *s*-haplotype (orange, n=12); *S*-flanking regions (grey, n=12). **b.** *H. palustris* (250-kb windows): *S*-haplotype (red, n=52); *s*-haplotype (orange, n=67); *S*-flanking regions (grey, n=68). **c.** *P. veris* (20-kb windows): *S*-haplotype (red, n=14); *S*-flanking regions (grey, n=14).

**Supplementary Tables (separate file)**
All supplementary tables (Table S1-S20) are included in a separate file.
